# Utilizing Heteroatom Types and Numbers from Extensive Ligand Libraries to Develop Novel hERG Blocker QSAR Models Using Machine Learning-based Classifiers

**DOI:** 10.1101/2023.08.05.552121

**Authors:** Safa Haddad, Lalehan Oktay, Ismail Erol, Kader Şahin, Serdar Durdagi

## Abstract

The human Ether-à-go-go Related Gene (hERG) channel plays a crucial role in membrane repolarization. Any disruptions in its function can lead to severe cardiovascular disorders like long QT syndrome (LQTS), which increases the risk of serious cardiovascular problems such as tachyarrhythmia and sudden cardiac death. Drug-induced LQTS is a significant concern and has resulted in drug withdrawals in the past. The main objective of this research study is to pinpoint crucial heteroatoms present in ligands that initiate interactions leading to effective blocking of the hERG channel. To achieve this aim, ligand-based quantitative structure-activity relationships (QSAR) models were constructed using extensive ligand libraries, considering the heteroatom types and numbers, and their associated hERG channel blockage pIC_50_ values. Machine learning-assisted QSAR models were developed to analyze the key structural components influencing compound activity. Among various methods, the KPLS method proved to be the most efficient, allowing the construction of models based on eight distinct fingerprints. The study delved into investigating the influence of heteroatoms on the activity of hERG blockers, revealing their significant role. Furthermore, by quantifying the effect of heteroatom types and numbers on ligand activity at the hERG channel, six compound pairs were selected for molecular docking. Subsequent molecular dynamics (MD) simulations and MM/GBSA calculations per residue were performed to comprehensively analyze the interactions of the selected pair compounds.

## 1. INTRODUCTION

Despite advancements in diagnosis and treatment, cardiovascular diseases (CVD) continue to be the leading cause of death worldwide, as highlighted by the World Health Organization (WHO). The mortality rate related to CVD is a significant concern, as seen in countries like Russia, where the annual rate is 614 deaths per 100,000 individuals, placing it among the highest worldwide [1]. The human Ether-à-go-go Related Gene (hERG), which encodes voltage-gated potassium ion channels, plays a critical role in the repolarization of cardiac action potential in human cardiomyocytes [2]. These potassium channels (K_V_) are integral membrane proteins that serve vital functions in various physiological processes. They are involved in generating nerve impulses, regulating neuronal excitability, controlling cardiac pacemaking, and modulating muscular contractility. The channels are composed of homotetramers, each with six trans-membrane segments (S1-S6). The Voltage Sensing Domain (VSD) comprises segments S1 to S4, while the Pore Domain (PD) consists of S5, S6, the P-Loop, and the Selectivity Filter (SF) that facilitates the permeation of K^+^ ions (Figure 1).

**Figure 1.**
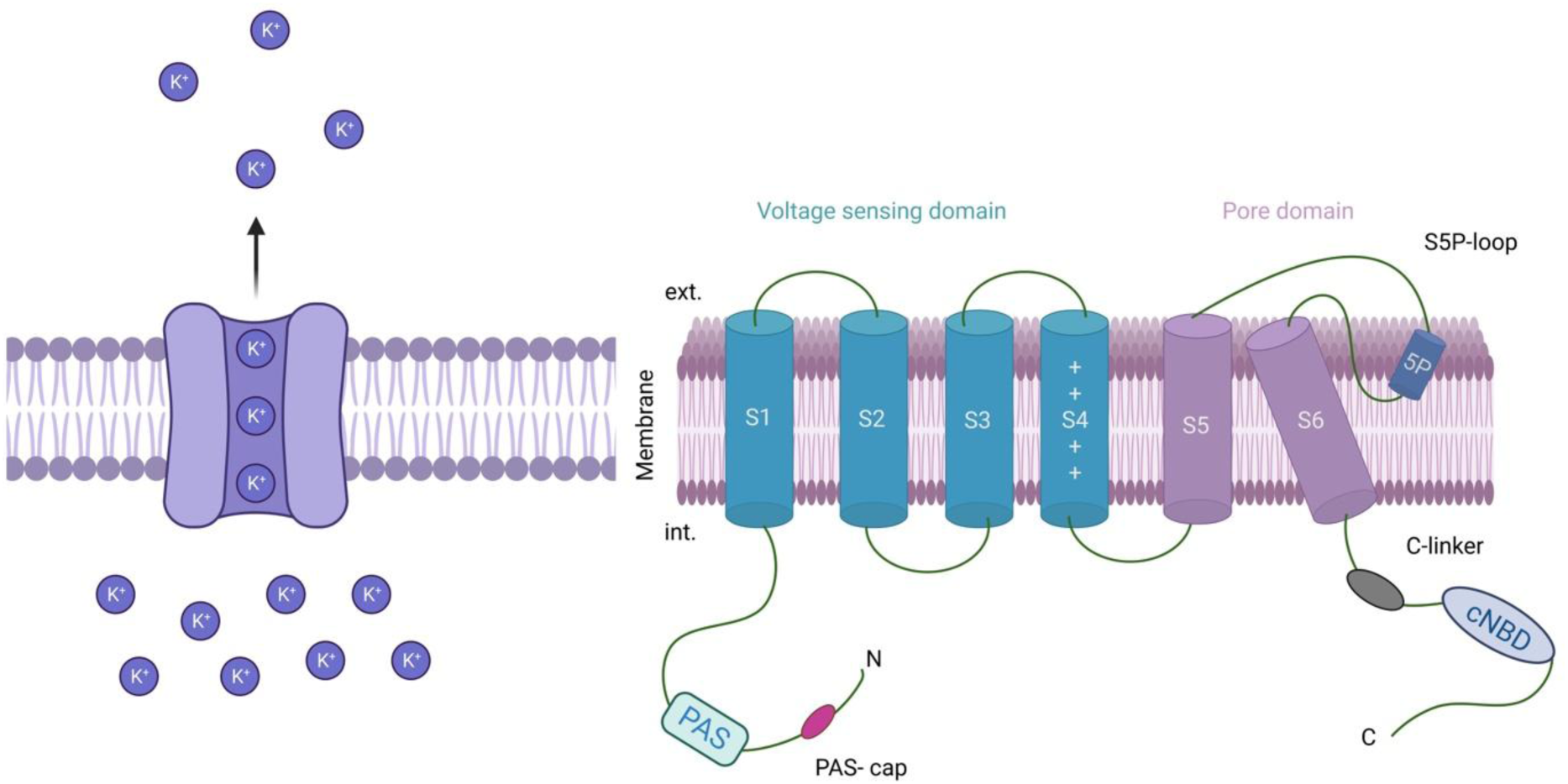
Kv11.1 (hERG) channel. While S1-S4 helices form voltage sensing domain, S5-S6 helices form pore domain region of the channel.

Prolonged ventricular repolarization caused by hERG channel blockade can increase the risk of cardiac damage, evident by an extended QT interval on electrocardiography (ECG). Paradoxically, the use of antiarrhythmic medications to prolong QT intervals poses the one of the highest risk of potentials. This risk has been also observed across various drug classes, including antihistamines, antipsychotics, antibiotics, and gastrointestinal stimulants [3]. Notably, medications such as cisapride (a serotonin receptor agonist), quinidine (an antiarrhythmic), astemizole, and terfenadine (both antihistamines) have either been withdrawn from the market or highly restricted due to their potential cardiotoxicity associated with hERG channel blockage [4]. These regulatory actions emphasize the significance of hERG-related risks and underscore the need to monitor and control the use of medications that may interfere with cardiac repolarization. Consequently, the hERG potassium channel has become a crucial therapeutic target, leading to the prioritization of screening drugs for their interaction with the hERG channel during early stages of drug design. International Conference on Harmonization (ICH) regulations now require the evaluation of drug candidates for their ability to block the hERG channel during preclinical testing [5]. This evaluation has become a pivotal aspect in the initial phases of drug research to ensure the safety and efficacy of new drugs, particularly concerning their potential to cause adverse effects on cardiac repolarization [6]. By conducting rigorous screening of hERG channel function, researchers can identify and eliminate drug candidates with unfavorable interactions, ultimately reducing the risk of cardiotoxicity and enhancing patient safety. In recent years, multiple mechanisms associated with QT prolongation have been discovered, highlighting the complexity beyond hERG channel blockade. Despite the long-standing recommendation by international regulatory agencies since 2005 to assess the inhibitory effect of new drugs on the hERG channel in preclinical settings [7], research continues to uncover additional factors influencing QT prolongation.

The comprehensive evaluation of QT prolongation risk, utilizing the multiple ion channel effect model that considers sodium and calcium channel blocking alongside hERG channel assay, proves to be more robust than relying solely on hERG assay for assessing drug safety. However, a significant number of currently available drugs undergo evaluation using only the hERG assay during the preclinical stage. As a result, efforts have been made to assess the risk of drug-induced QT prolongation at the preclinical and clinical level. Despite initiatives like *crediblemeds.org*, which aims to provide an updated list of QT-prolonging drugs, there are still instances where drugs with unknown risks of QT prolongation continue to be prescribed [8].

Presently, the assessment of hERG-associated cardiac toxicity involves employing *in vitro* and *in vivo* techniques to study the impact of potential hERG channel blockers and understand their effects on channel permeability. While *in vivo* experiments offer comprehensive drug evaluations, their high cost, inefficiency, and conflict with the 3R principle (which advocates for replacement, reduction, and refinement in animal studies) highlight the need to minimize their use. In recent years, *in vitro* assays have made progress in terms of duration and cost-effectiveness. However, these methods have also limitations in investigating the underlying structural mechanisms responsible for observed outcomes. To address this challenge, computational techniques have emerged as a promising approach in drug development for evaluating the hERG-blocking potential of small compounds before conducting experiments [9]. Computational methods provide insights into the structural basis of hERG channel blockade and can complement experimental approaches, contributing to a more efficient and informed drug discovery process. By utilizing computational modeling and simulations, researchers can prioritize and optimize potential drug candidates, reducing reliance on resource-intensive *in vivo* experiments and ultimately accelerating the development of safer and more effective medications.

*In silico* strategies, such as machine learning-based classifiers and structure-based modeling, offer valuable and reliable complements to experimental approaches in addressing the issue of hERG-associated cardiac toxicity. These computational methods leverage large datasets and advanced algorithms to analyze the structural and functional properties of hERG channels and their interactions with potential blockers. By training machine learning models on experimental data, researchers can accurately predict the hERG-blocking potential of novel compounds. Additionally, structure-based modeling techniques enable detailed exploration of the binding interactions between compounds and hERG channels, providing insights into the mechanisms of channel blockade. Integrating *in silico* approaches with experimental studies not only enhances the understanding of hERG-associated cardiac toxicity but also facilitates the identification of safer and more effective drug candidates in a more efficient and cost-effective manner [1].

In this study, ligand-based QSAR models were constructed using data on heteroatom types and numbers derived from extensive ligand libraries, incorporating information on chemical constitution and hERG channel blockage (pIC_50_ values). The QSAR models, developed using various methods, proved effective in identifying essential structural components influencing compound activity. The investigation of heteroatoms highlighted their role in the hERG activity of blockers. By quantifying the impact of heteroatom types and number in a compound on pIC_50_ value, six compound pairs were selected for docking, followed by molecular dynamics (MD) simulations and MM/GBSA calculations for a comprehensive analysis.

## 2. METHODS

The overall methodology of this study is summarized in Figure 2, which gives an overview of the techniques employed during the study.

**Figure 2.**
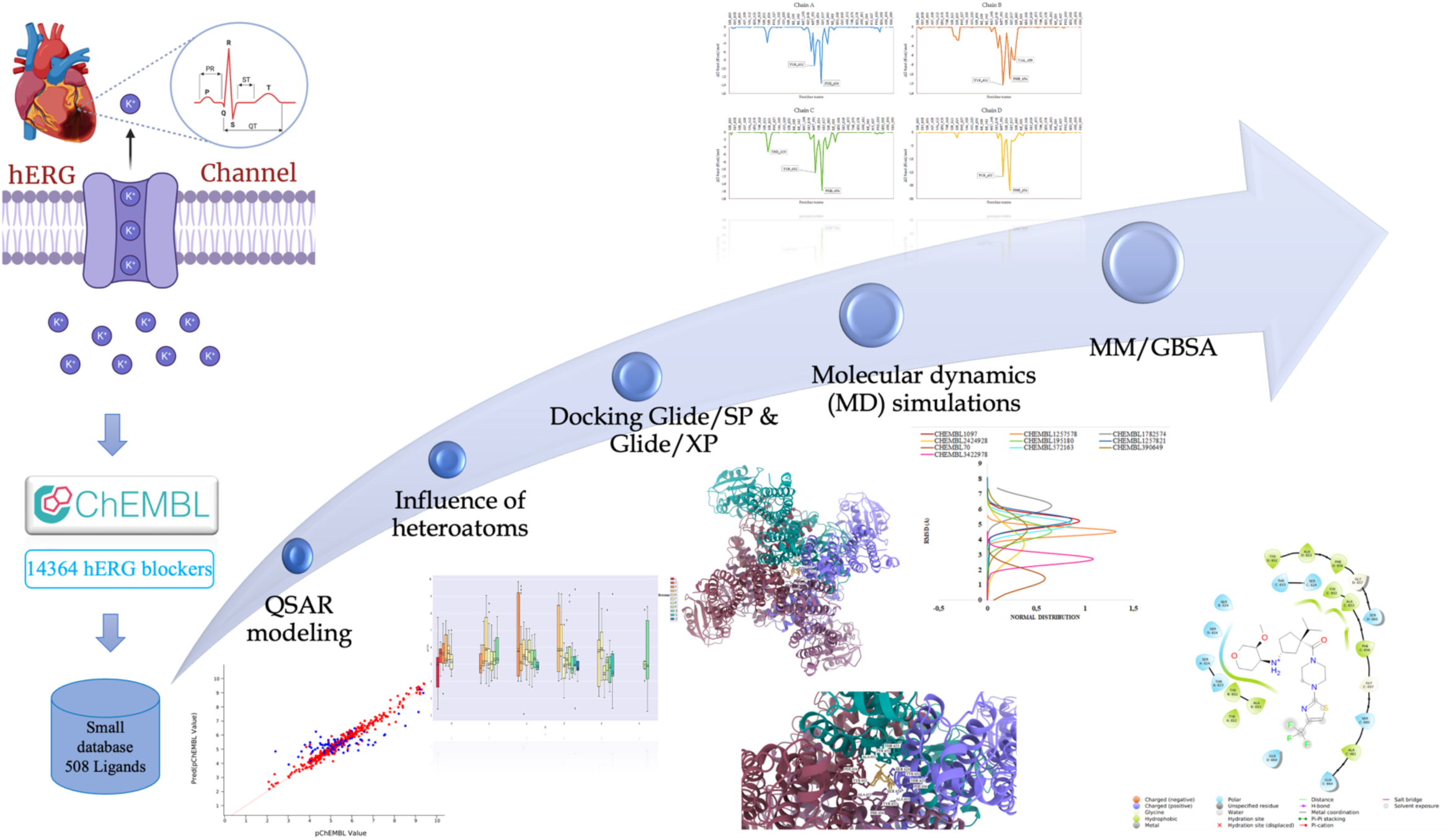
Overview of the methodology employed in this study.

### 2.1 Construction of ligand databases from ChEMBL

The ChEMBL database (https://www.ebi.ac.uk/chembl/) was employed to gather structural and biological activity data of known hERG blockers. As of November 2022, the database contained 14,364 available compounds with their corresponding IC_50_ values at the hERG channel. To standardize the activity values, the IC_50_ values were transformed into pIC_50_ by taking the negative logarithm (base 10) of each IC_50_ value. pIC_50_ is a widely used pharmacological indicator that quantifies the potency of a substance by measuring the concentration of ligand required to inhibit a specific biological activity. To manage the extensive database, a subset of 508 compounds was selected among 14,364 compounds. In order construct subset database, a normal distribution curve was generated, encompassing a range of pIC_50_ value spanning from 2.07 to 9.85 (Figure S1). This subset included compounds with varying levels of activity, providing a balanced representation of low, moderate, and high activity within the small database.

### 2.2 Building QSAR models

Schrodinger’s AutoQSAR module was employed to develop QSAR models, initially using DeepChem [10]. Machine-learning methods were used to create predictive models for the target data. To compare its efficiency with DeepChem, a similar process was repeated using the *traditional* QSAR method. Additionally, CANVAS cheminformatics package was utilized to build QSAR models. Physiochemical descriptors were calculated from the molecular properties of the 508 structures to enhance model generation. For model generation, Multiple Linear Regression (MLR), Partial Least-Squares (PLS) Regression, and Kernel-Based PLS Regression (KPLS) were employed. The statistical results of the constructed models, such as R² and Q², were assessed to determine the most effective method. KPLS models were developed using various types of fingerprints. The models that exhibited high scores for each fingerprint were selected, and scatter plots were used for visualization and atomic contributions analysis to identify influential substructures affecting compound activity.

### 2.3 Investigating the heteroatom types and numbers

To identify heteroatom types and numbers that significantly influence hERG blockage, we utilized the CANVAS cheminformatics package on each ligand structure. To ensure statistical reliability, only sets containing more than 5 compounds were considered for subsequent analyses. Median values were calculated for each dataset, and a Python script in the Spyder environment [11] was executed to perform correlation analysis. The script examines how changes in specific heteroatom types and numbers, in the presence of other heteroatoms, correlate with changes in pIC_50_ value. The resulting graphs illustrate the correlation between pIC50 value and the number and types of heteroatoms. Comparisons and calculations were conducted between sets with the same number of other heteroatoms, while observing the variations in the specific heteroatom’s count within the structure. To determine significant effects, the number of other heteroatoms was kept constant, and the impact of a particular heteroatom count was examined. If a substantial effect of at least around one log unit was detected between the median pIC_50_ values of the compared sets, the compound with the highest pIC_50_ was selected from the set with the higher median value, while the compound with the lowest pIC_50_ was chosen from the set with the lowest median value.

### 2.4 Ligand Preparation

The LigPrep module of Maestro molecular modeling package was employed to prepare the 508 ligands. During the ligand preparation process, the protonation states of the compounds were calculated within a pH range of 7.0 ± 2.0 using the Epik module. Compounds with chiral centers generated up to four distinct stereoisomers, and ionization states were considered for each molecule. To ensure accurate representation of molecular characteristics, the OPLS3e force field was applied.

### 2.5 Protein preparation

The cryo-EM structure of the hERG K^+^ channel (PDB ID: 5VA1) was obtained from the RCSB Protein Data Bank [12]. In this structure, the channel is in an open-like state, and the voltage sensors exhibit a depolarized shape. Despite its relatively small size, the central cavity contains four deep hydrophobic pockets, which may explain the heightened sensitivity of the hERG channel to diverse ligand structures. To enable molecular docking and MD simulations, the channel was prepared using the Protein Preparation tool in the Maestro molecular modeling suite [13]. Before the MD simulations, three potassium ions and two water molecules were added to the selectivity filter (SF) of the channel. Additionally, missing side chains and loops were filled. The Maestro’s Prime module [14], [15] was employed to incorporate the missing loops and side chains into the residues. PROPKA [16], [17] was used to add hydrogen atoms to the protein at physiological pH, ensuring accurate ionization states of amino acid residues. The structural optimization was then performed using the OPLS3 force field with a convergence criterion of 0.3 Å RMSD for heavy atoms.

### 2.6 Receptor grid generation

The generation of a receptor grid is a crucial step in docking studies. It involves creating a three-dimensional grid that represents the binding site of the receptor, which is necessary to evaluate the binding energy of ligands during docking simulations. The Receptor Grid Generation tool was utilized to identify the binding pocket (active site) in the structure. [18] The grid was generated by specifying key residues (i.e., Thr623, Ser624, Tyr652, Ala 653, and Phe656) located at the centroid of selected residues. (Figure 3) The coordinates for the center of the grid were determined as (2.86. −5.97. −1.27) in the (x, y, z) coordinate system, respectively.

**Figure 3.**
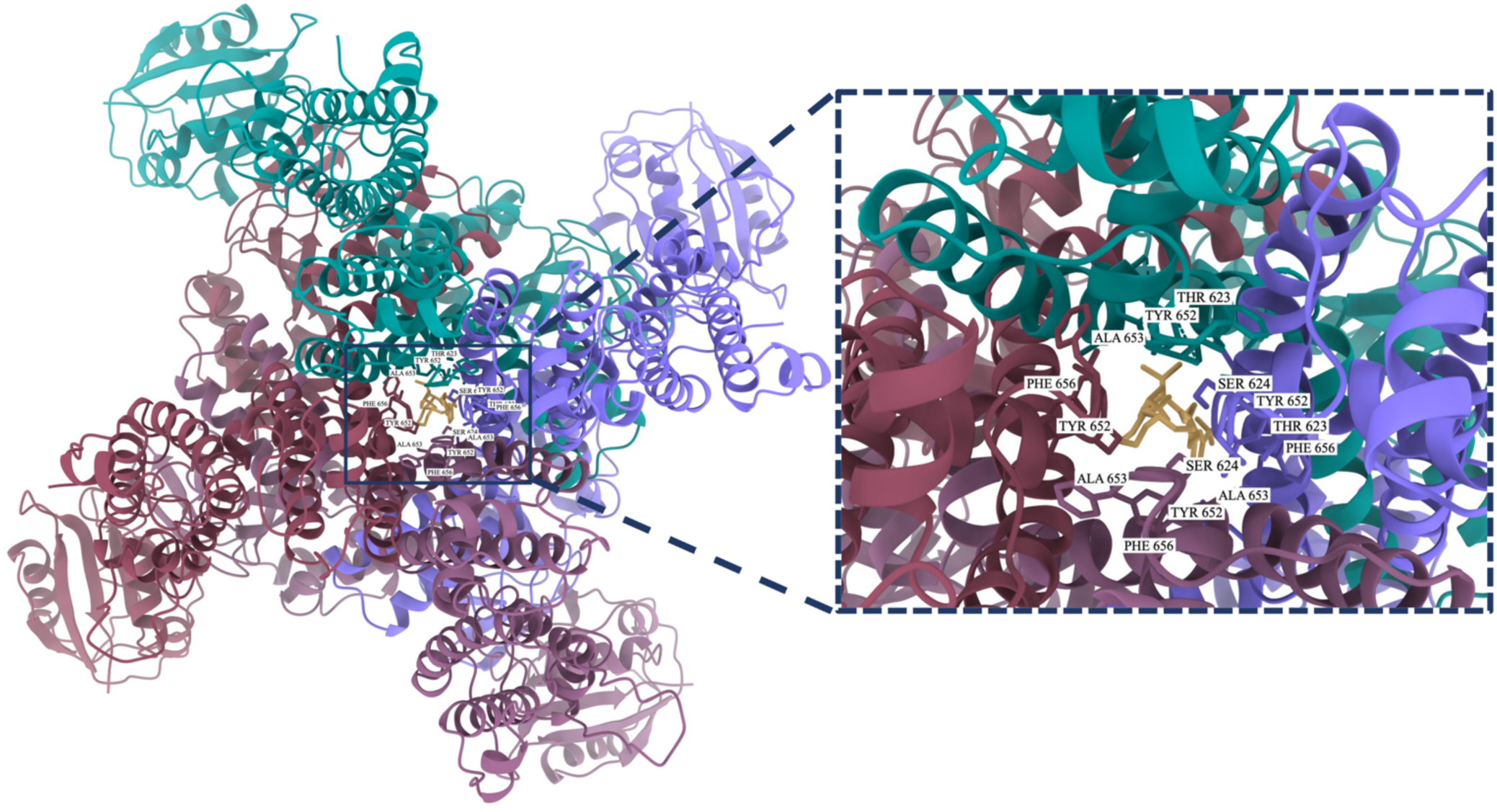
Docking pose of CHEMBL1782574 in the central cavity of the hERG channel.

### 2.7 Molecular docking

The docking simulations were conducted using the grid-base Glide docking algorithm [19]– [21], which systematically explores the binding site of the receptor. The ligands underwent hierarchical filtration to assess the complementarity of the ligand-receptor system. Ligands that passed this phase underwent energy minimization and were assigned scores. Two docking protocols, Glide/SP and Glide/XP, were employed in this study. Standard parameters, including ligand sampling with nitrogen inversions and ring conformations, bias sampling of torsion for amides, and post-docking minimization of 5 poses for each ligand were considered. On the other hand, Glide/XP performed deeper sampling, beginning with SP sampling before utilizing its own anchor-and-grow technique. It utilized a more complex scoring algorithm, demanding higher ligand-receptor shape complementarity. The Glide/XP docking protocol employed standard settings with flexible ligand sampling, Epik state penalties incorporated into the docking score, and post-docking minimization of 10 poses. The docked poses were ranked based on docking scores, and only the top-scoring poses were selected for further analysis using MD simulations.

### 2.8 Molecular dynamics (MD) simulations

The molecules with the highest docking scores were subjected to MD simulations using the Desmond. [22] The membrane-embedded structure (PDB ID: 5VA1) was obtained from the OPM database. [23] The system builder placed the 5VA1 protein in an orthorhombic solvation box using the TIP3P solvent model [24] and a POPC membrane model at 310 K, Figure 4. The simulations were conducted in the NPT ensemble at 310 K with a pressure of 1.01325 bar, maintained using a Nose-Hoover thermostat [25] and Martyina-Tobias-Klein barostat. [26] The system was balanced by adding Cl^−^ ions and 0.15 M of NaCl solution to achieve a pH of 7.4. Prior to the simulations, the Desmond software performed energy minimization and relaxation of the structure. Each MD simulation conducted for 200 ns generating 1000 frames. The relevant MD simulation data were collected individually and saved in trajectory files.

**Figure 4.**
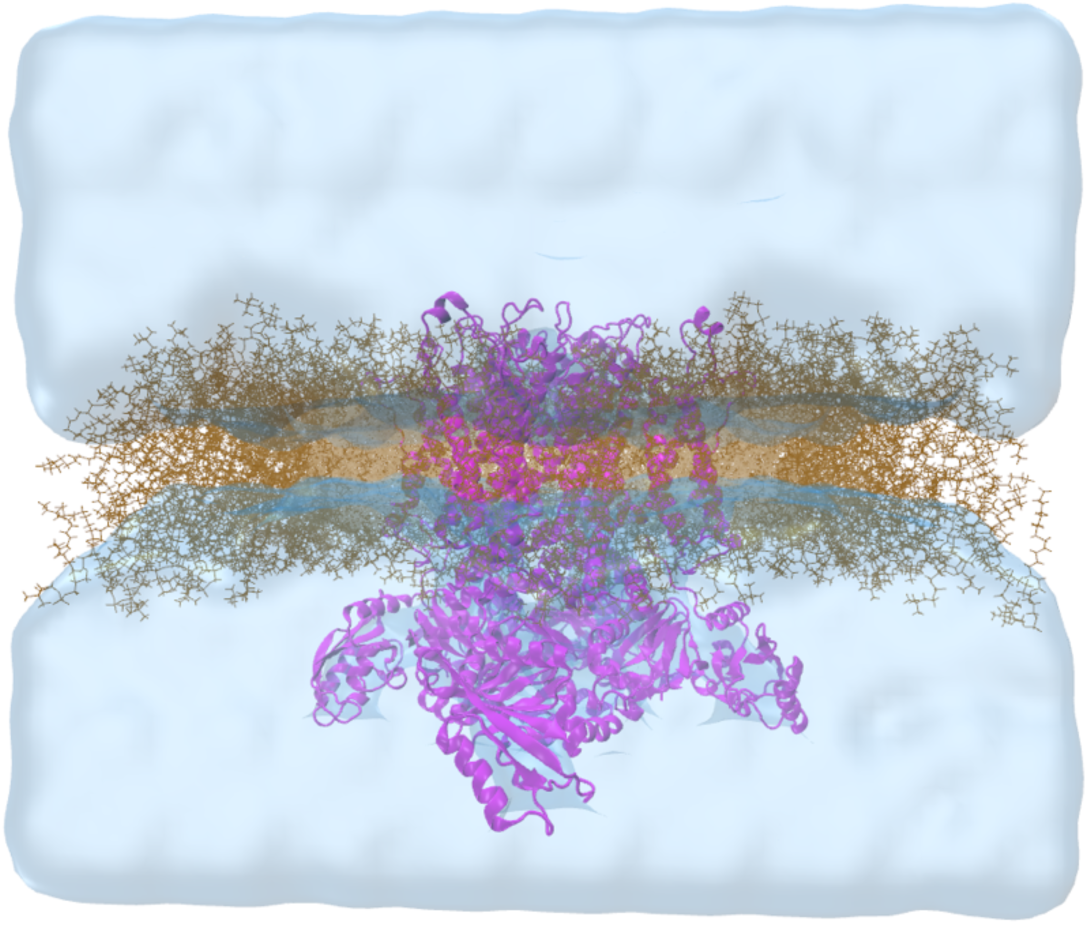
hERG channel (PDB, 5VA1) merged with POPC membrane bilayer. Figure represents water molecules as surface.

### 2.9 Molecular mechanics generalized Born surface area (MM/GBSA) calculations

To calculate binding free energies and understand the main differences between weak and strong inhibitors, MM/GBSA analysis was performed using Maestro’s Prime module. [14], [15] A systematic approach was employed, wherein MM/GBSA calculations were performed using one frame out of every ten frames. The dielectric constant was defined using the VSGB 2.0 implicit solvation model, with the exterior network considered as a water system with a constant dielectric constant and the interior dielectric constant allowed to vary. After the calculations for each complex, the average MM/GBSA value and standard deviations for each compound were computed. Furthermore, to gain insight into the contribution of individual residues in the four chains of the hERG channel to the inhibitory activity and frequency of ligand-protein contacts, per-residue MM/GBSA analysis was conducted to calculate binding free energies on a residue level. This analysis provides more detailed information about the role of specific residues in the interaction between the ligands and the protein.

## 3. Results and discussion

The main objective of this study is to create ligand-based QSAR models by utilizing heteroatom types and numbers from extensive ligand libraries, which encompass information on the chemical structure of ligands and their respective hERG channel blockage IC_50_ values. Furthermore, the study seeks to explore atomic-level features that enhance the affinity with the hERG channel and influence the inhibitory activity of compounds.

A dataset consisting of 14364 hERG blocker compounds was collected from the ChEMBL database for this study. In order to manage easily the constructed models, a smaller subset database is considered among 14364 hERG blockers. For this aim, evenly distributed biological activity of compounds is selected which involve 508 compounds. (Figure S1) To ensure the reliability of the collected IC_50_/pIC_50_ values, they were also cross-checked from scientific literature. These 508 compounds were selected to generate a normal distribution curve, which provides insights into the distribution of hERG blocking potential within the dataset. The pIC_50_ values for these compounds ranged from 2.07 to 9.85. QSAR models were constructed to investigate the relationship between the structure and activity of hERG blocker compounds, providing valuable insights into the crucial structural elements that influence their activity. Various QSAR modeling techniques were employed to assess their effectiveness on the given dataset. The results revealed that the models developed using *traditional* methods exhibited elevated R² and Q² values, indicating superior precision and predictability, albeit with lower or moderate Ranking score values (Table S1). Similarly, models developed using the *DeepChem* method also performed well in terms of R² and Q² values, but with lower Ranking score values (Table S2). When constructing the QSAR models, having a reliable objective function, or “score,” becomes essential to distinguish between effective (i.e., stable) and ineffective QSAR models. In AutoQSAR, the quality of the models is evaluated based on their performance on specific training and test sets. The accuracy of a model is represented by a value between 0 and 1, where 1 indicates perfect predictions and 0 signifies entirely incorrect predictions. The model M is then scored accordingly:

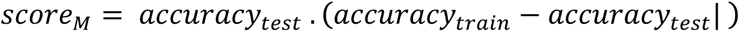

This formula is designed to provide favorable scores to models that demonstrate high accuracy on both the training and test sets. Conversely, it penalizes models that exhibit low accuracy on either or both sets, as well as models that show substantial discrepancies in accuracy between the two sets. [10]

Consistency in the effectiveness of the models was observed across various descriptor sets and molecular fingerprints. Among different methods compared, the KPLS approach outperformed PLS and MLR (Table S3). Consequently, QSAR models utilizing machine learning-based classifiers were developed using the KPLS method along with eight different fingerprints (*atom pairs, atom triplets, dendritic, linear, MACCS, mol_print, Radial,* and *topological*). To assess the impact of each fingerprint type on the statistical scores of the constructed QSAR models, the CANVAS tool was employed. The same dataset containing 508 hERG blocker compounds with their corresponding pIC_50_ values was used in CANVAS. The results showed that the KPLS approach, when using *atom triplets*, *MACCS*, and *Radial fingerprints*, yielded low Q^2^ values (Table 1). However, other fingerprints like *dendritic, topological, mol_print, linear*, and *atom pairs* produced both high R^2^ and Q^2^ values. The models with superior statistical results for each fingerprint type are presented in Table 1. Overall, these findings underscore the significance of successful QSAR modeling techniques, particularly the KPLS approach, in elucidating the structure-activity relationships of hERG blocker compounds. These models offer valuable insights for designing potent and precise molecules. The use of different fingerprints and modeling methods allows for a comprehensive analysis, leading to a better understanding of the dataset’s characteristics. The QSAR models displayed a strong correlation between predicted and observed pIC_50_ values. (Figure 5) These models effectively predicted the activity of both the training and test sets, demonstrating their efficacy in predicting the potency of hERG blocker compounds.

**Table 1.**
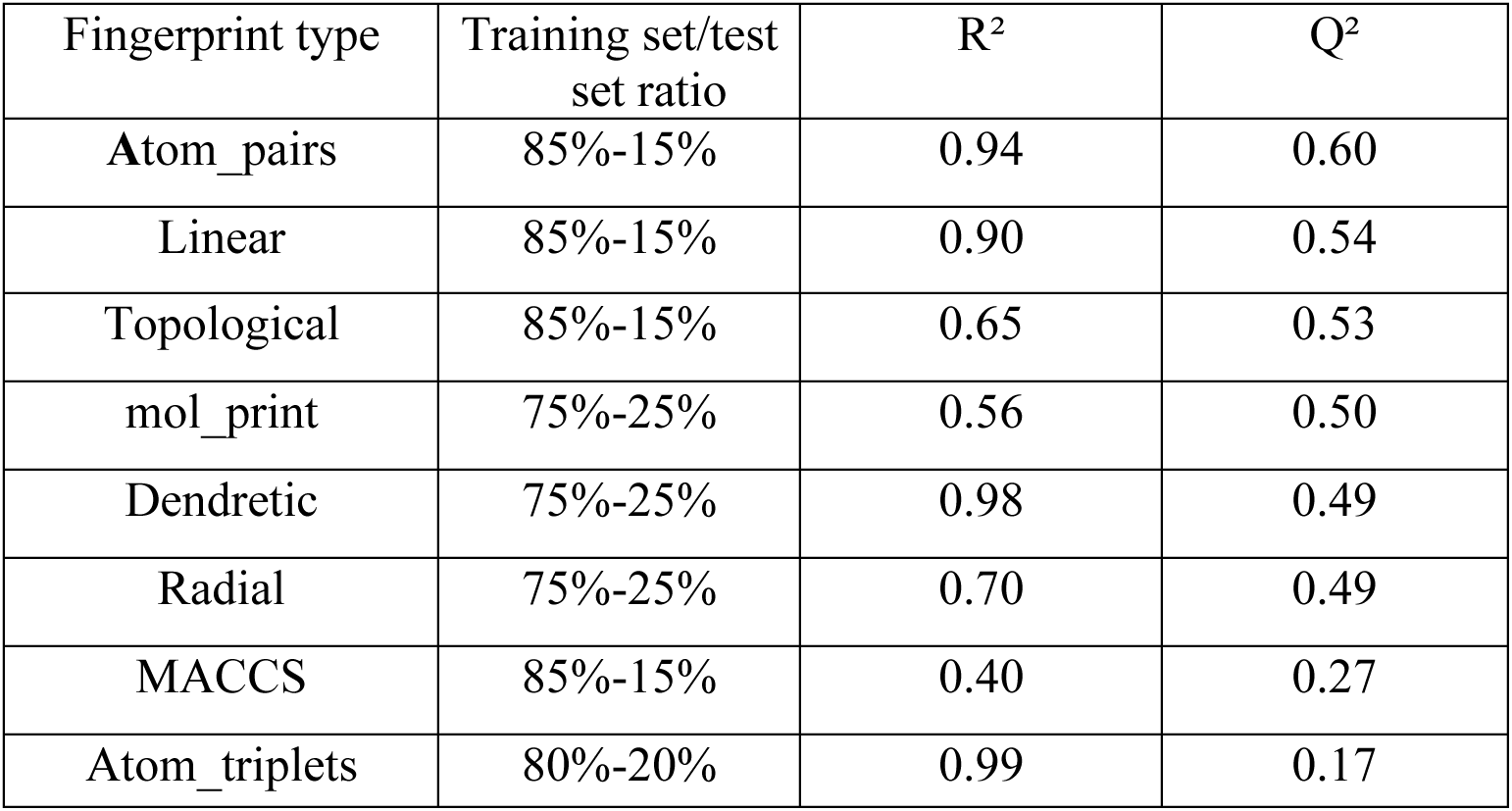
Performance evaluation of top KPLS-based fingerprint models generated by CANVAS.

| Fingerprint type | Training set/test set ratio | R<sup>2</sup> | Q<sup>2</sup> |
| --- | --- | --- | --- |
| Atom_pairs | 85%-15% | 0.94 | 0.60 |
| Linear | 85%-15% | 0.90 | 0.54 |
| Topological | 85%-15% | 0.65 | 0.53 |
| mol_print | 75%-25% | 0.56 | 0.50 |
| Dendretic | 75%-25% | 0.98 | 0.49 |
| Radial | 75%-25% | 0.70 | 0.49 |
| MACCS | 85%-15% | 0.40 | 0.27 |
| Atom_triplets | 80%-20% | 0.99 | 0.17 |

**Figure 5.**
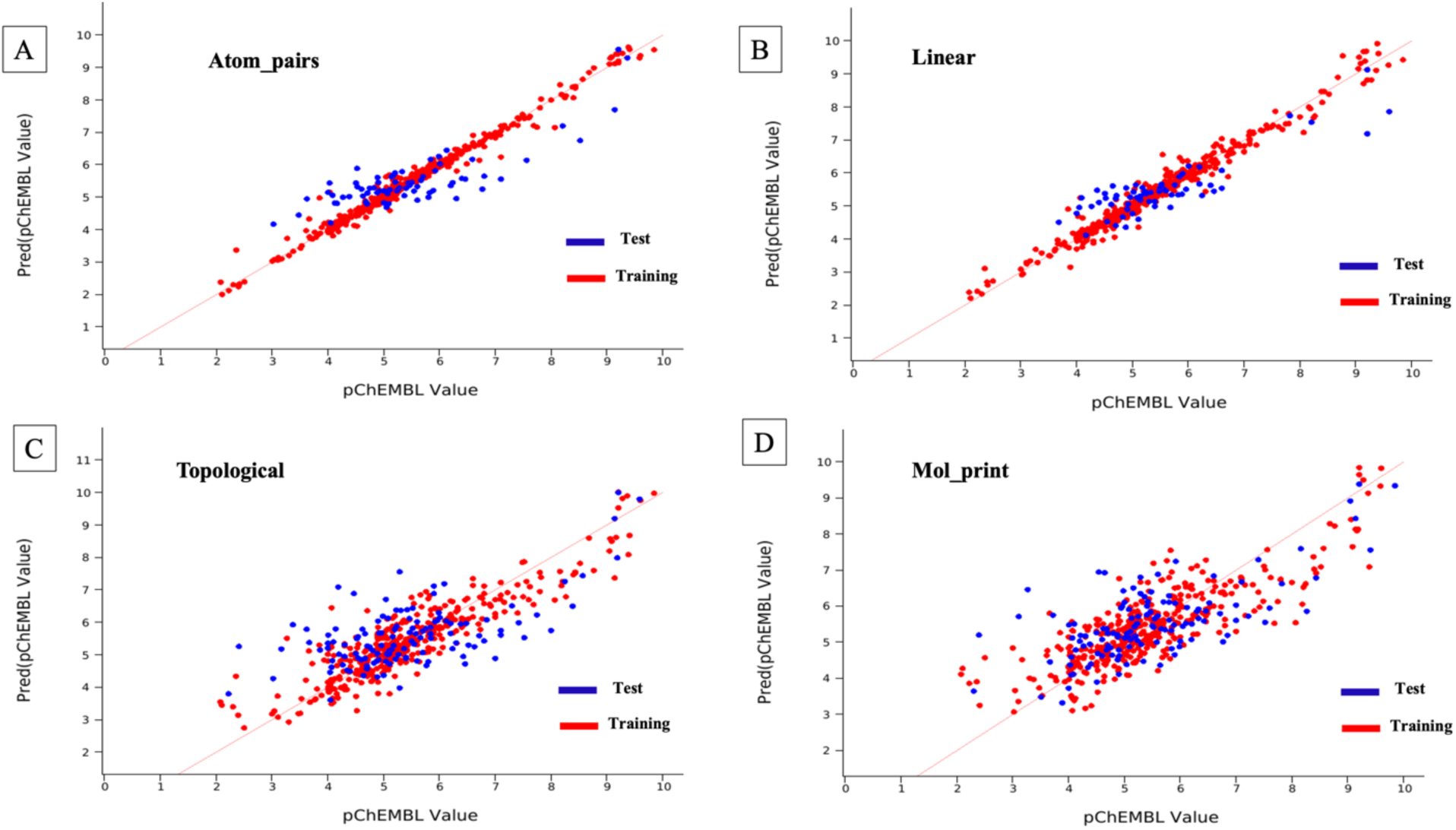
The scatter plot of observed and predicted pIC_50_ values of the hERG blockers based on KPLS model with A) *atompairs*, B) *linear*, C) *topological*, D) *mol_print* fingerprints.

Additionally, an investigation into the influence of heteroatoms on the functionality of hERG inhibitors was undertaken, focusing on the quantity and variety of heteroatoms present in the compounds. The LigFilter descriptors were computed using the CANVAS [27], [28], excluding carbon and hydrogen atoms. This allowed for the determination of the number of heteroatoms in a dataset comprising 508 compounds (Tables S4-S7), providing valuable insights into the heteroatom composition within the compounds under study.

The analysis revealed that oxygen, nitrogen, chlorine, and sulfur atoms were particularly important heteroatoms in the 508 selected compounds, modulating the inhibition activity of hERG blockers. Oxygen atoms exhibited an inverse relationship with activity, as an increase in their number within the compounds led to decreased inhibition (Figure 6). Similarly, an increase in chlorine atoms resulted in a slight decrease in activity (Figure S2). On the other hand, the presence of sulfur atoms displayed a positive correlation with activity, indicating that as the sulfur atom content increased within a compound, the pIC_50_ value also tended to increase (Figure S3). However, the presence of nitrogen atoms did not show a discernible impact on the biological activity, with no clear correlation between the pIC_50_ value and the number of nitrogen atoms in the compound. Nevertheless, a closer examination revealed that an increase in the number of nitrogen atoms from 1 to 3 within a compound resulted in a decrease in pIC_50_ values, but when the number of nitrogen atoms exceeded 3, the trend in pIC_50_ values showed an increase. Furthermore, when there were more than 5 nitrogen atoms in a compound, the pIC_50_ trend was not stable (Figure S4). These findings provide valuable insights into the structure-activity relationships of hERG blockers, particularly concerning the influence of heteroatom types and numbers.

**Figure 6.**
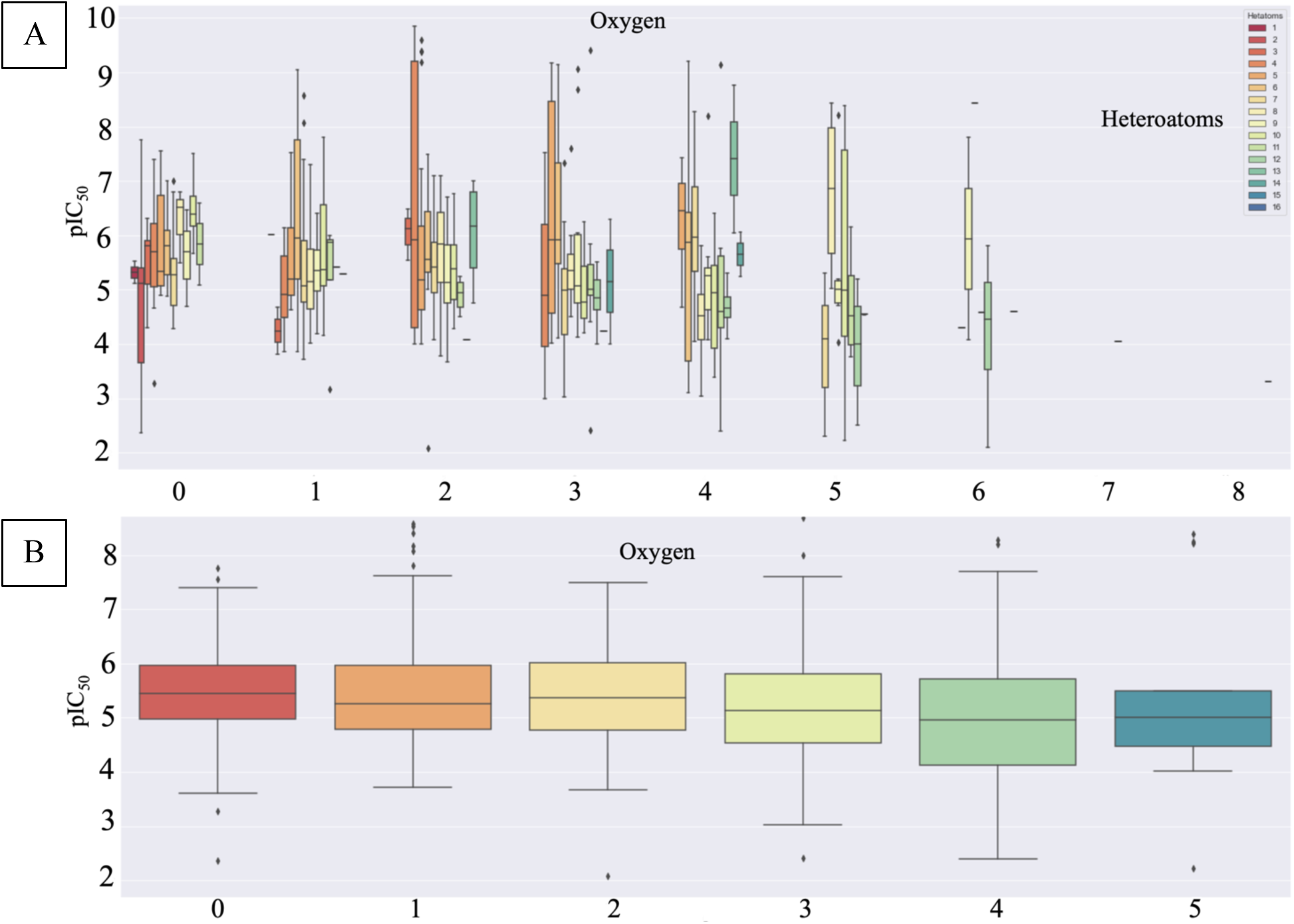
Influence of heteroatoms on the activity of the hERG blockers. (A) the effect of the number of oxygen atoms to the hERG blockage activity (pIC_50_). (B) Detailed analyzes on the influence of the oxygen atoms on the pIC_50_ values.

To further investigate the correlation between heteroatoms and pIC_50_ value, additional analyses were conducted by grouping compounds with the same number of a specific heteroatom type. This approach allowed for a more targeted analysis of how biological activity (pIC_50_) evolves with changes in the quantity of a particular heteroatom type, while keeping the total count of heteroatoms in the molecular structure constant. Figures 6B, S2B, S3B, and S4B depict these grouped analyses, offering a clearer understanding of the relationship between heteroatoms and activity. Further statistical analysis was performed to examine the quantitative impact of heteroatoms on pIC_50_ activity, focusing on changes of approximately 1 log unit or more for the effect of a specific atom on the activity. To ensure reliable sampling, a threshold of five compounds in a cluster was set, and clusters containing fewer than five compounds were excluded from the analysis. By evaluating the increase or decrease in the number of a specific heteroatom while keeping the number of other heteroatoms constant, significant effects on activity were observed. To gain a deeper understanding of these effects, compounds with the highest and lowest pIC_50_ values were selected from each set, encompassing both strong and weak inhibitors, providing a comprehensive view of the differences influencing the activity. As a result, six pairs of molecules were identified, each pair consisting of a strong and weak inhibitor (Table 2).

**Table 2.** Quantitative analysis of the influence of heteroatoms on pIC_50_ values.

| Molecular Formula of the compound representing the **weakest** inhibition within the set and its ChEMBL ID | pIC<sub>50</sub> value of the compound representing the **weakest** inhibition within the set | Heteroatom type and number representing the **weakest** inhibition within the set | Total number of heteroatom representing the **weakest** inhibition within the set | Mean pIC<sub>50</sub> value for the **weak** inhibition set | Molecular Formula of the compound representing the **strongest** inhibition within the set | pIC<sub>50</sub> value of the compound representing the **strongest** inhibition within the set | Heteroatom type and number representing the **strongest** inhibition within the set | Total number of Heteroatom representing the **strong** inhibition within the set | Mean pIC<sub>50</sub> value for the **strong** inhibition set | Log unit difference between median values |
| --- | --- | --- | --- | --- | --- | --- | --- | --- | --- | --- |
| **C17H19NO3**<br>(70) | 3 | 1 N | 4 | 4.89 | **C22H26N2O2**<br>(1257821) | 9.85 | 2 N | 4 | 5.92 | 1.03 |
| **C15H23N3O2**<br>(1097) | 4 | 3 N | 5 | 5.175 | **C26H31NO4**<br>(572163) | 7.42 | 1 N | 5 | 6.1 | 0.93 |
| **C15H23N3O2**<br>(1097) | 4 | 3 N | 5 | 5.175 | **C20H24N2O2S**<br>(1257578) | 9.59 | 2 N | 5 | 6.17 | 0.99 |
| **C23H35F3N4O3S** (1782574) | 2.41 | 4 N | 11 | 4.81 | **C26H26F3N5O3**<br>(3422978) | 9.41 | 5 N | 11 | 5.76 | 0.95 |
| **C24H26FN5O4**<br>(2424928) | 3.51 | 0 S | 10 | 4.96 | **C28H31F3N6S**<br>(390649) | 7.5 | 1 S | 10 | 5.9 | 0.94 |
| **C15H23N3O2**<br>(1097) | 4 | 0 Cl | 5 | 5.22 | **C24H27ClN2OS**<br>(195180) | 7.52 | 1 Cl | 5 | 6.72 | 1.5 |

This approach enabled a comprehensive investigation of the discrepancies among these molecules and facilitated subsequent analyses. To gain deeper insights into the impact of the type and number of heteroatoms in these selected representative compound pairs, molecular docking analyses were carried out using the Glide/SP and Glide/XP algorithms. These chosen molecules were docked to the pore domain region of the hERG channel. The outcomes of these simulations, comprising the ligand-protein interactions (Figure 7) and docking scores obtained from Glide/SP and Glide/XP, are presented in Table 3. Following the selection of the top docking scores, the compounds were subjected to further analysis using MD simulations. A total of 6 pairs, representing both strong and weak inhibitors, were utilized in 200 ns all-atom MD simulations. The resulting trajectories from each simulation were collected and subjected to analysis. To assess the structural stabilities of each ligand pair at the binding pocket, the root-mean-square deviation (RMSD) of the ligands was measured in comparison to their initial positions (i.e., Glide/XP). A higher ligand RMSD value generally indicates a deviation from the initial binding pose, while a lower RMSD value suggests that the molecule maintains its initial conformation throughout the simulations. As expected, the results from Figure 8 validate these assumptions, as the RMSD values consistently remain lower for the strong inhibitors and higher for the weak inhibitors. However, it is worth noting that the pair of molecules CHEMBL70 (pIC_50_, 3) and CHEMBL1257821 (pIC_50_, 9.85) deviates from this trend. These findings provide valuable insights into the structural dynamics and binding characteristics of the inhibitors, emphasizing the significance of heteroatom types and numbers in achieving a strong or weak inhibitory profile. The ligand-protein interactions were analyzed through the generation of a simulation interaction diagram. Throughout the 200 ns all-atom MD simulations, various contacts between the ligand and the protein were observed and recorded. Figures S5-S10 present detailed interactions between the ligand atoms and the protein, focusing on interactions that occurred for more than 15% of the simulation time. These analyses shed light on the complex interactions between the ligands and the protein, further contributing to our understanding of the structure-activity relationships and binding mechanisms of the inhibitors.

**Figure 7.**
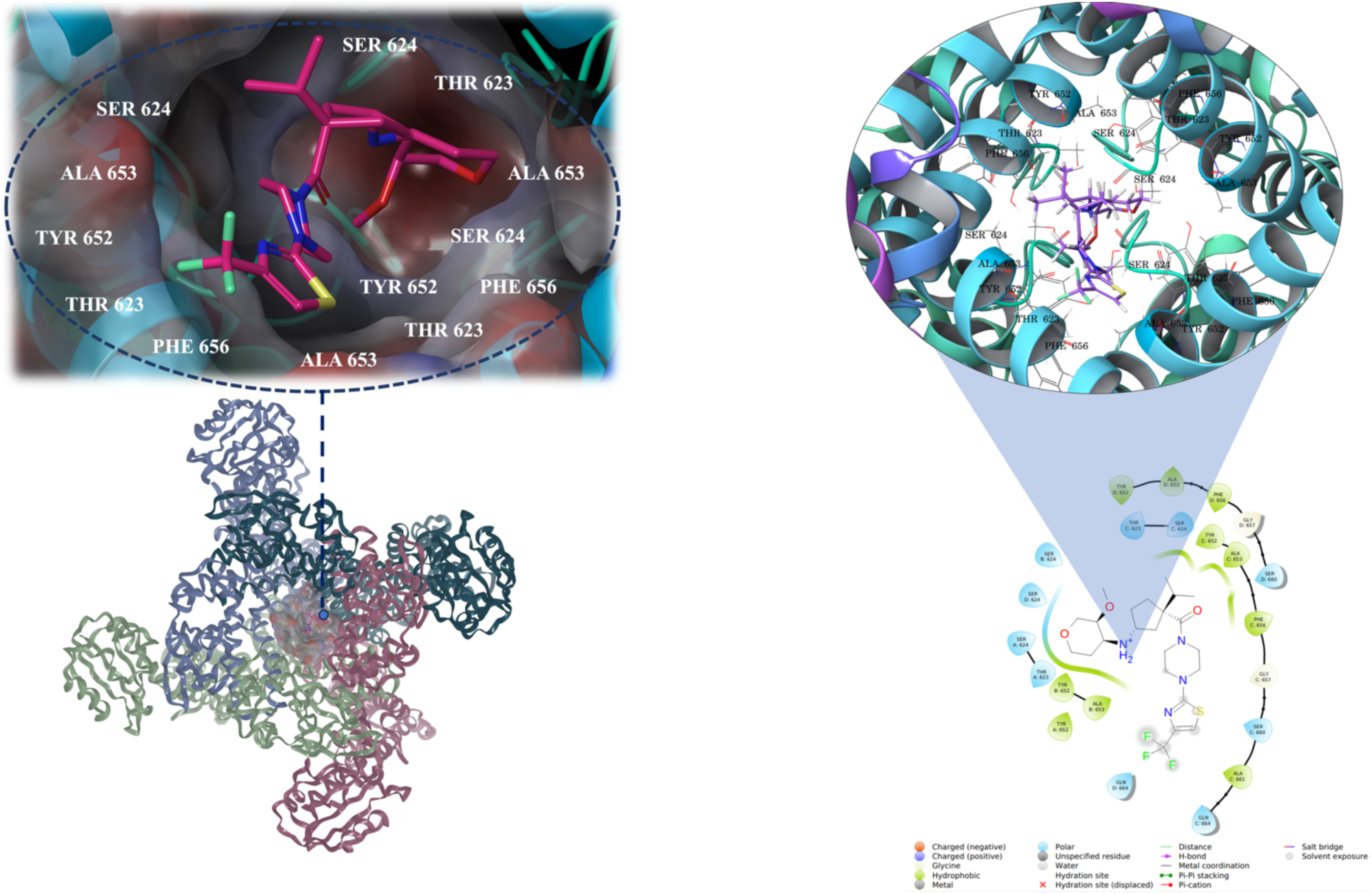
Investigating ligand-protein interactions within the intracavitary pore domain of the channel: Examination of the docking pose of CHEMBL1782574 compound with 2D and 3D insights.

**Table 3.** Docking scores from Glide/SP and XP of top selected ligands in the hERG binding site (central cavity).

| Pair Number | ChEMBL ID | pIC<sub>50</sub> | Glide (kcal/mol) |  |
| --- | --- | --- | --- | --- |
|  |  |  | SP | XP |
| Pair 1 | CHEMBL70 | 3.00 | -5.33 | -5.14 |
|  | CHEMBL1257821 | 9.85 | -6.84 | -6.12 |
| Pair 2 | CHEMBL572163 | 7.42 | -6.34 | -4.12 |
|  | CHEMBL1097 | 4.00 | -3.95 | -5.21 |
| Pair 3 | CHEMBL1257578 | 9.59 | -6.79 | -5.87 |
|  | CHEMBL1097 | 4.00 | -3.95 | -5.21 |
| Pair 4 | CHEMBL1782574 | 2.41 | -5.34 | -4.42 |
|  | CHEMBL3422978 | 9.41 | -7.08 | -6.29 |
| Pair 5 | CHEMBL2424928 | 3.51 | -6.69 | -5.21 |
|  | CHEMBL390649 | 7.5 | -4.92 | -5.24 |
| Pair 6 | CHEMBL1097 | 4.00 | -3.95 | -5.21 |
|  | CHEMBL195180 | 7.52 | -6.84 | -5.01 |

**Figure 8.**
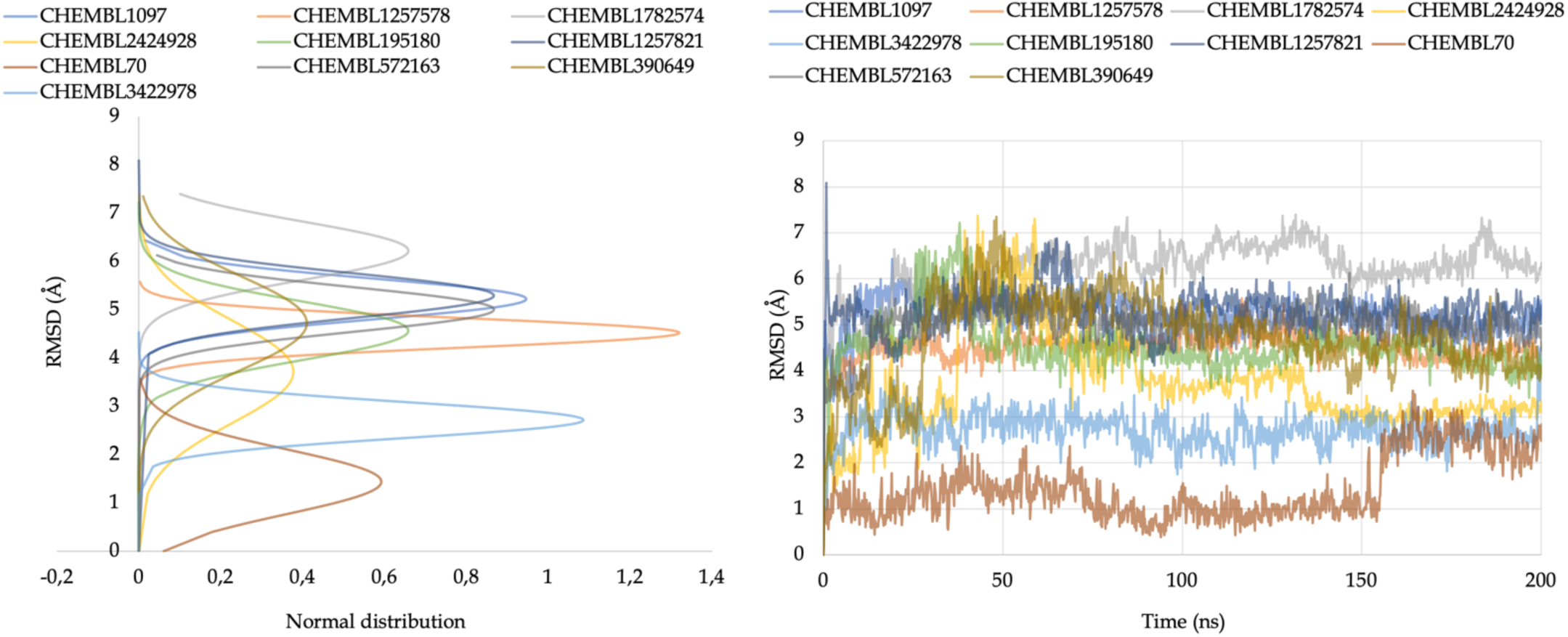
Ligand RMSD analyses. (A) Normal distribution of ligand RMSDs, (B) change of RMSD value of heavy atoms of ligands based on their initial positions at the binding pocket of the channel (LigFitProt).

The MM/GBSA results were thoroughly examined alongside the experimental pIC_50_ data for each pair of molecules. Table 4 presents valuable insights into the average MM/GBSA binding free energy between the ligand and the receptor. As anticipated, the MM/GBSA outcomes revealed a clear correlation between binding energy and activity. Molecules with higher pIC_50_ values exhibited more negative binding energy values, indicating stronger interactions and a better fit with the pore domain of the ion channel. Conversely, molecules with lower pIC_50_ values showed less negative binding energy values, suggesting weaker binding and less favorable interactions, categorizing them as weak inhibitors. The overall agreement between the MM/GBSA results and the experimental data substantiated the predictive capability of the MM/GBSA method. However, it is crucial to acknowledge that the MM/GBSA results for the pair of molecules CHEMBL70 and CHEMBL1257821 did not align well with the experimental data. This discrepancy signals the necessity for further investigation to comprehend the underlying factors contributing to the observed deviation in this specific pair of molecules. These findings underscore the importance of incorporating both experimental data and computational approaches like MM/GBSA in evaluating ligand-ion channel interactions and predicting binding affinities. By combining these approaches, a more comprehensive understanding of molecular interactions and potential inhibitory activity can be achieved, aiding in the design and optimization of effective inhibitors.

**Table 4.** Average MM/GBSA values for the selected compounds.

| Pair Number | ChEMBL ID | Average MM/GBSA (kcal/mol) | Std. Dev. (kcal/mol) | pIC<sub>50</sub> |
| --- | --- | --- | --- | --- |
| Pair 1 | CHEMBL70 | -53.20 | 4.63 | 3.00 |
|  | CHEMBL1257821 | -53.09 | 6.80 | 9.85 |
| Pair 2 | CHEMBL572163 | -64.71 | 6.70 | 7.42 |
|  | CHEMBL1097 | -44.65 | 6.46 | 4.00 |
| Pair 3 | CHEMBL1257578 | -70.80 | 8.00 | 9.59 |
|  | CHEMBL1097 | -44.65 | 6.46 | 4.00 |
| Pair 4 | CHEMBL1782574 | -58.90 | 7.23 | 2.41 |
|  | CHEMBL3422978 | -70.99 | 5.84 | 9.41 |
| Pair 5 | CHEMBL2424928 | -62.32 | 7.16 | 3.51 |
|  | CHEMBL390649 | -63.61 | 10.96 | 7.5 |
| Pair 6 | CHEMBL1097 | -44.65 | 6.47 | 4.00 |
|  | CHEMBL195180 | -70.07 | 6.29 | 7.52 |

In general, the results are in line with our expectations, except for two particular compounds: CHEMBL70 and CHEMBL1257821. While CHEMBL70, a kappa opioid receptor antagonist, was anticipated to have a smaller RMSD and a more negative average MM/GBSA score compared to CHEMBL1257821, the latter displayed a strong inhibitory effect at the hERG channel with a pIC_50_ value of 9.85. However, the mismatched results observed for this pair may be attributed to the state of the hERG channel. The hERG channel can exist in either an open state or an open-inactivated state for hosting the blockers, and different drugs may have varying affinities depending on the specific state. It is plausible that CHEMBL1257821 and CHEMBL70 interact differently with the hERG channel due to variations in the channel’s conformational state. Further investigations and studies are warranted to explore the specific binding mechanisms and interactions of these compounds with the hERG channel in different states. Understanding these nuances can provide crucial insights into the design and development of more effective hERG inhibitors.

In this study, we also conducted a comprehensive MM/GBSA analysis to identify the crucial residues within the hERG channel that significantly impact the inhibition activity. The goal of this analysis was to pinpoint the residues that establish strong interactions with the ligands compared to others. The results of the MM/GBSA per-residue study, depicted in Figures 9, S11-S19, offer valuable insights into the specific residues involved in the binding processes of the ligands at the pore domain. The findings of the per-residue analysis shed light on the essential residues responsible for governing the inhibitory function of the hERG channel. As expected, residues like Phe656 and Tyr652 were identified as key contributors, forming strong contacts with the ligands (Figure 9). Interestingly, each ligand at the pore domain also exhibited additional specific residues that played a role in binding to the hERG channel. For example, compound CHEMBL1257578 showed that Val625 from Chain-A, Leu650 from Chain-B, Thr623, Val625, Met645, and Val659 from Chain-C, and Met645 from Chain-D were significant contributors to the binding energy, in addition to Tyr652 and Phe656 (Figure 9).

**Figure 9.**
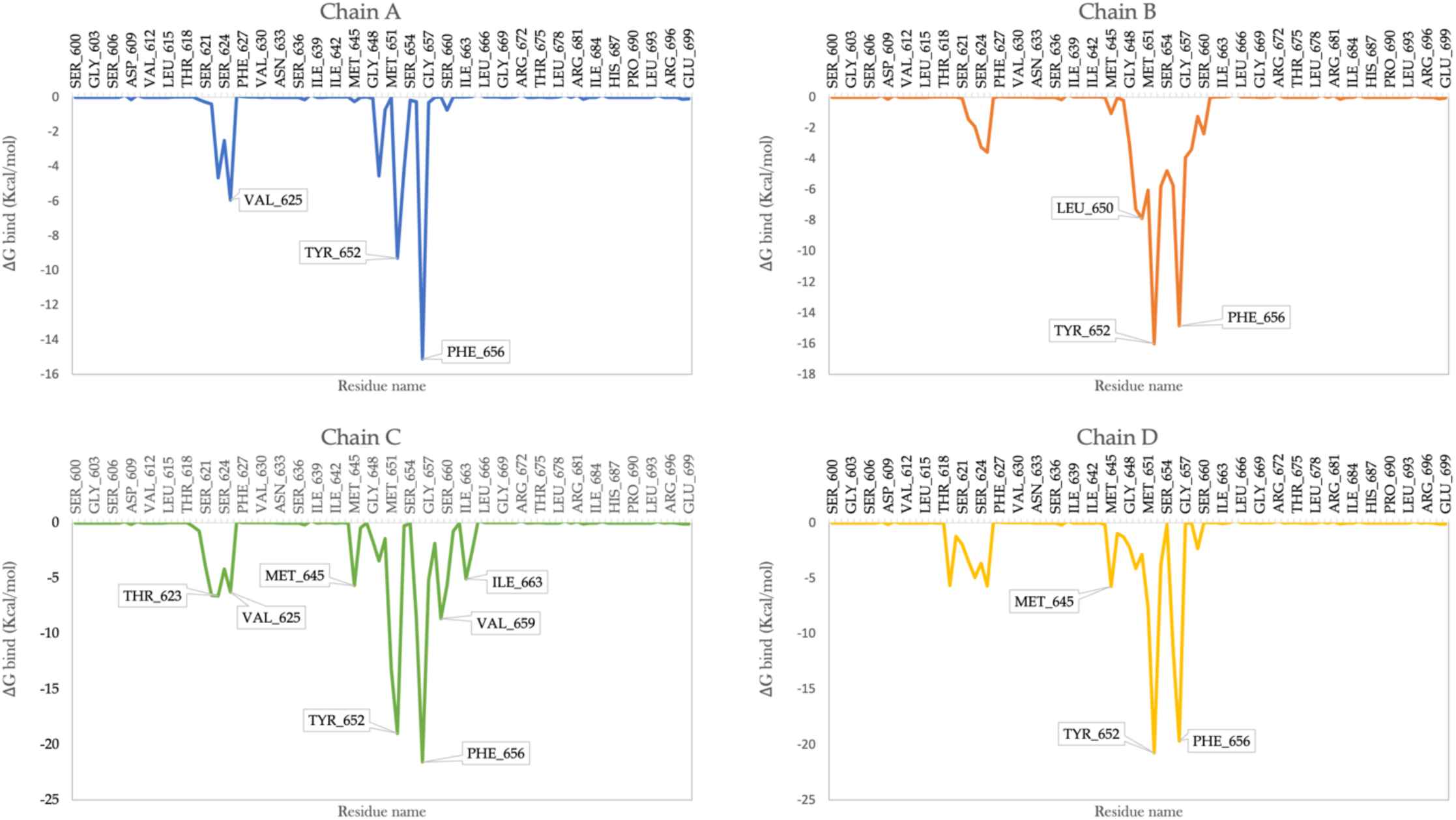
Per-residue MM/GBSA analyses for CHEMBL1257578 compound.

## 4. Conclusions

In conclusion, this study has provided valuable insights into the structure-activity relationships of hERG blockers and the substructures that influence their inhibition activity. The development of ligand-based QSAR models based on heteroatom numbers from extensive ligand libraries proved to be effective, with CANVAS-created QSAR models showing high R² and Q² values. Among the various QSAR modeling techniques employed, the KPLS method demonstrated superior performance, enabling the development of QSAR models using multiple fingerprints. The investigation of heteroatoms revealed significant correlations between the number of oxygen, nitrogen, chlorine, and sulfur atoms in the chemical structure of the inhibitors and their pIC50 values. The increase in the quantity of oxygen and chlorine atoms demonstrated an inverse correlation with inhibition activity, whereas the increased number of sulfur atoms displayed a parallel relationship. However, the impact of nitrogen atoms on hERG channel blocking activity proved to be more complex. It was observed that when the number of nitrogen atoms increased from 1 to 3 within a compound, pIC50 values showed a decreasing trend. However, when the number of nitrogen atoms exceeded 3, a reverse trend emerged, with pIC50 values increasing. Furthermore, having more than 5 nitrogen atoms in a compound did not show a stable pIC50 trend. These findings underscore the importance of heteroatoms in influencing the activity of hERG blockers. The study highlights the significant role that specific heteroatom types and their respective quantities play in modulating the inhibitory effects on the hERG channel. Understanding these relationships can contribute to the design and optimization of potent and selective hERG inhibitors.

The utilization of MD simulations and MM/GBSA calculations enabled a comprehensive analysis of compound pairs, facilitating a comparison between strong and weak inhibitors. The results aligned perfectly with expectations, as strong inhibitors displayed lower RMSD values and more negative average MM/GBSA binding energy values, indicating stronger binding to the hERG channel. On the other hand, weak inhibitors exhibited higher RMSD values and less negative average MM/GBSA values, indicating a weaker binding affinity. Moreover, the MM/GBSA per-residue analysis identified critical residues in the four chains of the hERG channel that play a significant role in blockage activity. Notably, Tyr652 emerged as a key residue, demonstrating consistent and diverse interactions with all of the studied inhibitor ligands. Tyr652 engaged in the highest number of interactions, accounting for over 50% of the simulation time, primarily driven by hydrophobic forces. Additionally, other crucial residues such as Phe656, Ser660, Thr623, and Ser624 were found to actively participate in ligand interactions throughout the simulation period. These findings highlight the pivotal role of Tyr652 and other residues in establishing and modulating inhibitor-ligand interactions, providing valuable insights for the design and optimization of targeted therapeutics.

In summary, this study enhances our comprehension of hERG channel blockage and offers valuable insights that can aid in the design of more effective hERG blockers. The presented findings play a crucial role in advancing drug discovery and provide significant considerations for the development of targeted therapeutics.

