## Supporting Tables and Figures for "Utilizing Heteroatom Types and Numbers from Extensive Ligand Libraries to Develop Novel hERG Blocker QSAR Models Using Machine Learning-based Classifiers"

### Supporting Information

**Table S1.** Results of the top Auto-QSAR traditional method models.

| Numeric model | KPLS_radial_19 | KPLS_radial_49 | KPLS_radial_45 |
|---|---|---|---|
| Ranking score | 0.41 | 0.50 | 0.48 |
| Training set / Test set ratio | 75%-25% | 85%-15% | 85%-15% |
| $R^2$ | 0.79 | 0.78 | 0.78 |
| $Q^2$ | 0.54 | 0.60 | 0.57 |

**Table S2.** Results of the top Auto-QSAR DeepChem models.

| QSAR Model | Model code | Training set / Test set ratio | Score | $R^2$ | $Q^2$ |
|---|---|---|---|---|---|
| qsar_build_15 | dl_1 | 80%-20% | 0.28 | 0.96 | 0.51 |
| qsar_build_35 | dl_1 | 75%-25% | 0.33 | 0.97 | 0.56 |
| qsar_build_34 | dl_1 | 75%-25% | 0.37 | 0.94 | 0.58 |

**Table S3.** Results of the MLR and PLS method models from CANVAS.

| approach | $Q^2$ | $R^2$ | Training set/Test set ratio |
|---|---|---|---|
| MLR | -0.01 | 0.08 | 80%-20% |
| | 0.03 | 0.06 | 85%-15% |
| | 0.03 | 0.07 | 75%-25% |
| | 0.008 | 0.08 | 70%-30% |
| PLS | 1.15 | 0.24 | 80%-20% |
| | 0.24 | 0.20 | 85%-15% |
| | 0.21 | 0.19 | 75%-25% |
| | 0.21 | 0.21 | 70%-30% |

**Table S4.** Investigation of the influence of Chlorine atom on the $pIC_{50}$ value.

| Name | Molecular formula | pIC50 | Cl | Hetatoms | Median Value |
|---|---|---|---|---|---|
| CHEMBL75880 | C19H35N | 5.11 | 0 | 1 | 5.315 |
| CHEMBL629 | C20H23N | 5.52 | 0 | 1 | |
| CHEMBL284348 | C5H6N2 | 2.36 | 0 | 2 | 5.11 |
| CHEMBL3 | C10H14N2 | 3.61 | 0 | 2 | |
| CHEMBL1086033 | C17H22N2 | 3.67 | 0 | 2 | |
| CHEMBL87045 | C18H18N2 | 5.06 | 0 | 2 | |
| CHEMBL459176 | C20H22N2 | 5.16 | 0 | 2 | |
| CHEMBL11 | C19H24N2 | 5.47 | 0 | 2 | |
| CHEMBL83 | C26H29NO | 6.01 | 0 | 2 | |
| CHEMBL1671894 | C27H26N2 | 7.75 | 0 | 2 | |
| CHEMBL1087 | C15H22N2O | 3.81 | 0 | 3 | 5.825 |
| CHEMBL1083707 | C20H24N2O | 4.67 | 0 | 3 | |
| CHEMBL513258 | C19H19FN2 | 5.05 | 0 | 3 | |
| CHEMBL2325209 | $C26H28NO2^{+}$ | 5.54 | 0 | 3 | |
| CHEMBL497171 | C15H17N3 | 5.8 | 0 | 3 | |
| CHEMBL457930 | C20H19N3 | 5.85 | 0 | 3 | |
| CHEMBL2324248 | $C20H20NO2^{+}$ | 6.12 | 0 | 3 | |
| CHEMBL1090528 | C19H24N2S | 6.21 | 0 | 3 | |
| CHEMBL299499 | C17H22IN3 | 6.31 | 0 | 3 | |
| CHEMBL607 | C15H21NO2 | 6.49 | 0 | 3 | |
| CHEMBL70 | C17H19NO3 | 3 | 0 | 4 | 5.01 |
| CHEMBL1671896 | C11H14F3N | 3.27 | 0 | 4 | |
| CHEMBL640 | C13H21N3O | 3.86 | 0 | 4 | |
| CHEMBL219803 | C17H25N3O | 3.94 | 0 | 4 | |
| CHEMBL16 | C15H12N2O2 | 4 | 0 | 4 | |
| CHEMBL3356251 | C24H36N2O2 | 4.01 | 0 | 4 | |
| CHEMBL219128 | C17H33N3O | 4.17 | 0 | 4 | |
| CHEMBL3786542 | C26H26N2O2 | 4.21 | 0 | 4 | |

**Table S4** (cont.d)

| | | | | | |
|---|---|---|---|---|---|
| CHEMBL3356249 | C25H38N2O2 | 4.3 | 0 | 4 | |
| CHEMBL219074 | C20H31N3O | 4.42 | 0 | 4 | |
| CHEMBL3314426 | C15H14F2N2 | 4.66 | 0 | 4 | |
| CHEMBL1086273 | C19H23FN2O | 4.68 | 0 | 4 | |
| CHEMBL1089159 | C22H23N3O | 4.7 | 0 | 4 | |
| CHEMBL2333646 | C19H23BrN2O | 4.89 | 0 | 4 | |
| CHEMBL3220615 | C19H21NO3 | 4.89 | 0 | 4 | |
| CHEMBL3185736 | C22H20N2O2 | 4.91 | 0 | 4 | |
| CHEMBL517 | C21H29N3O | 4.92 | 0 | 4 | |
| CHEMBL4162513 | C35H34N2O2 | 5.1 | 0 | 4 | |
| CHEMBL562948 | C22H33N3O | 5.4 | 0 | 4 | |
| CHEMBL1090175 | C20H25N3O | 5.47 | 0 | 4 | |
| CHEMBL1935443 | C20H23N3O | 5.68 | 0 | 4 | |
| CHEMBL1093058 | C17H21N3S | 5.7 | 0 | 4 | |
| CHEMBL4450665 | C23H28N2O2 | 5.92 | 0 | 4 | |
| CHEMBL3323181 | C22H24N2OS | 6 | 0 | 4 | |
| CHEMBL460893 | C23H25N3O | 6.13 | 0 | 4 | |
| CHEMBL1092377 | C19H21FN2S | 6.48 | 0 | 4 | |
| CHEMBL1346 | C28H30N2O2 | 7.1 | 0 | 4 | |
| CHEMBL2046892 | C22H22F2N2 | 7.4 | 0 | 4 | |
| CHEMBL96153 | C24H31NO3 | 7.52 | 0 | 4 | |
| CHEMBL1223951 | C21H24N2O2 | 9.21 | 0 | 4 | |
| CHEMBL1257938 | C22H26N2O2 | 9.21 | 0 | 4 | |
| CHEMBL1257937 | C22H26N2O2 | **9.28** | **0** | **4** | |
| CHEMBL1257820 | C23H28N2O2 | 9.6 | 0 | 4 | |
| CHEMBL1257821 | C22H26N2O2 | 9.85 | 0 | 4 | |
| CHEMBL1097 | C15H23N3O2 | 4 | 0 | 5 | 5.22 |
| CHEMBL4640366 | C33H34N2O3 | 4.02 | 0 | 5 | |
| CHEMBL4209354 | C15H20N2O2S | 4.07 | 0 | 5 | |
| CHEMBL1091879 | C19H25N3O2 | 4.12 | 0 | 5 | |

**Table S4** (cont.d)

| | | | | |
|---|---|---|---|---|
| CHEMBL561138 | C23H27N3O2 | 4.14 | 0 | 5 |
| CHEMBL1258006 | C22H23N3O2 | 4.27 | 0 | 5 |
| CHEMBL3787069 | C26H26N2O3 | 4.35 | 0 | 5 |
| CHEMBL1094041 | C29H29N3O2 | 4.55 | 0 | 5 |
| CHEMBL4213196 | C25H34N4O | 4.62 | 0 | 5 |
| CHEMBL914 | C32H39NO4 | 4.67 | 0 | 5 |
| CHEMBL217133 | C18H22N4O | 4.68 | 0 | 5 |
| CHEMBL429115 | C20H24N4O | 4.71 | 0 | 5 |
| CHEMBL2441636 | C22H33N3O2 | 4.72 | 0 | 5 |
| CHEMBL1125 | C16H18N4S | 4.90 | 0 | 5 |
| CHEMBL4096683 | C20H22N4O | 4.94 | 0 | 5 |
| CHEMBL237191 | C25H31N5 | 4.98 | 0 | 5 |
| CHEMBL565829 | C15H14N4O | 5.01 | 0 | 5 |
| CHEMBL2177304 | C27H24FN3O | 5.07 | 0 | 5 |
| CHEMBL487063 | C23H27N3O2 | 5.10 | 0 | 5 |
| CHEMBL378547 | C21H21F2NO2 | 5.11 | 0 | 5 |
| CHEMBL485242 | C25H29N3O2 | 5.14 | 0 | 5 |
| CHEMBL478615 | C23H27N3O2 | 5.17 | 0 | 5 |
| CHEMBL2326478 | C18H18FN3S | 5.17 | 0 | 5 |
| CHEMBL476579 | C24H29N3O2 | 5.18 | 0 | 5 |
| CHEMBL479242 | C23H27N3O2 | 5.18 | 0 | 5 |
| CHEMBL3121096 | C20H25N3O2 | 5.22 | 0 | 5 |
| CHEMBL498572 | C19H22N2O3 | 5.23 | 0 | 5 |
| CHEMBL1819137 | C22H25N3O2 | 5.23 | 0 | 5 |
| CHEMBL488249 | C24H27N3O2 | 5.25 | 0 | 5 |
| CHEMBL4100591 | C27H34F2N2O | **5.29** | **0** | **5** |
| CHEMBL1956112 | C21H28N4S | 5.33 | 0 | 5 |
| CHEMBL487064 | C23H27N3O2 | 5.55 | 0 | 5 |
| CHEMBL1110 | C17H18N4O | 5.64 | 0 | 5 |
| CHEMBL478616 | C23H27N3O2 | 5.8 | 0 | 5 |

**Table S4** (cont.d)

| | | | | | |
|---|---|---|---|---|---|
| CHEMBL1224697 | C22H28N2O2S | 5.9 | 0 | 5 | |
| CHEMBL196983 | C21H27NO4 | 6.1 | 0 | 5 | |
| CHEMBL58387 | C31H38N4O | 6.24 | 0 | 5 | |
| CHEMBL3775729 | C26H28N4O | 6.28 | 0 | 5 | |
| CHEMBL376488 | C32H31BrN2O2 | 6.43 | 0 | 5 | |
| CHEMBL1671893 | C23H30N2O3 | 6.6 | 0 | 5 | |
| CHEMBL1087493 | C17H26N4O | 6.63 | 0 | 5 | |
| CHEMBL715 | C17H20N4S | 6.74 | 0 | 5 | |
| CHEMBL2324243 | $C26H32NO4^{+}$ | 6.8 | 0 | 5 | |
| CHEMBL572163 | C26H31NO4 | 7.42 | 0 | 5 | |
| CHEMBL61301 | C19H21FN4 | 7.56 | 0 | 5 | |
| CHEMBL1642487 | C28H28N2O3 | 9.09 | 0 | 5 | |
| CHEMBL1642479 | C28H28N2O3 | 9.17 | 0 | 5 | |
| CHEMBL259732 | C20H27N3O2 | 9.19 | 0 | 5 | |
| CHEMBL1257577 | C19H22N2O2S | 9.37 | 0 | 5 | |
| CHEMBL410832 | C23H25N3O2 | 9.39 | 0 | 5 | |
| CHEMBL1257578 | C20H24N2O2S | 9.59 | 0 | 5 | |
| CHEMBL270190 | C25H32N2O4 | 3.10 | 0 | 6 | 5.92 |
| CHEMBL519266 | C22H28N2O4 | 3.11 | 0 | 6 | |
| CHEMBL4212656 | C24H31N5O | 4.64 | 0 | 6 | |
| CHEMBL398612 | C26H35FN4O | 4.88 | 0 | 6 | |
| CHEMBL460491 | C21H34N4S2 | 4.88 | 0 | 6 | |
| CHEMBL1823042 | C20H28FN3O2 | 5 | 0 | 6 | |
| CHEMBL4067770 | C20H21FN4O | 5.11 | 0 | 6 | |
| CHEMBL4100776 | C20H22N4OS | 5.23 | 0 | 6 | |
| CHEMBL225036 | C24H26N4OS | 5.29 | 0 | 6 | |
| CHEMBL1938433 | C23H24N2O4 | 5.42 | 0 | 6 | |
| CHEMBL4089699 | C19H19N3O2S | 5.42 | 0 | 6 | |
| CHEMBL2177904 | C23H16FN5 | 5.51 | 0 | 6 | |
| CHEMBL1823047 | C22H30N4O2 | 5.54 | 0 | 6 | |

**Table S4** (cont.d)

| | | | | | |
|---|---|---|---|---|---|
| CHEMBL4065208 | C23H22N4O2 | 5.56 | 0 | 6 | |
| CHEMBL4104102 | C20H22N4OS | 5.61 | 0 | 6 | |
| CHEMBL2177905 | C24H16F2N4 | 5.8 | 0 | 6 | |
| CHEMBL3219616 | C24H30N2O3S | 5.8 | 0 | 6 | |
| CHEMBL1224699 | C21H26N2O3S | 5.9 | 0 | 6 | |
| CHEMBL561279 | C25H34N2O3S | 5.93 | 0 | 6 | |
| CHEMBL2147024 | C22H18N6 | 5.96 | 0 | 6 | |
| CHEMBL4081790 | C25H19N5O | 6.3 | 0 | 6 | |
| CHEMBL556100 | C22H25NO4S | 6.33 | 0 | 6 | |
| CHEMBL723 | C24H26N2O4 | 6.46 | 0 | 6 | |
| CHEMBL4096145 | C22H27N5O | 6.52 | 0 | 6 | |
| CHEMBL3774817 | C25H25FN4O | 6.59 | 0 | 6 | |
| CHEMBL522456 | CHEMBL522456 | 7 | 0 | 6 | |
| CHEMBL4173253 | C26H34N2O3S | 7.1 | 0 | 6 | |
| CHEMBL3775282 | C26H28N4O2 | 7.16 | 0 | 6 | |
| CHEMBL1108 | C22H22FN3O2 | 7.49 | 0 | 6 | |
| CHEMBL329067 | C32H33N5O | 7.62 | 0 | 6 | |
| CHEMBL533 | C20H36N2O3S | 8 | 0 | 6 | |
| CHEMBL12780 | C24H27FN4O | 8.16 | 0 | 6 | |
| CHEMBL1423 | C28H29F2N3O | 8.52 | 0 | 6 | |
| CHEMBL296419 | C28H31FN4O | 9.05 | 0 | 6 | |
| CHEMBL1258503 | C22H25N3O3 | 9.14 | 0 | 6 | |
| CHEMBL1258280 | C22H24N2O4 | 9.21 | 0 | 6 | |
| CHEMBL374731 | C10H14N2O5 | 2.3 | 0 | 7 | 5.13 |
| CHEMBL8 | C17H18FN3O3 | 3.02 | 0 | 7 | |
| CHEMBL4855895 | C21H24FN3O3 | 3.53 | 0 | 7 | |
| CHEMBL22 | C14H18N4O3 | 3.62 | 0 | 7 | |
| CHEMBL578834 | C21H25N5OS | 3.72 | 0 | 7 | |
| CHEMBL3582294 | C25H18F2N4O | 4.03 | 0 | 7 | |
| CHEMBL4224807 | C14H16N6O | 4.03 | 0 | 7 | |

**Table S4** (cont.d)

| | | | | |
|---|---|---|---|---|
| CHEMBL388978 | C28H26N4O3 | 4.04 | 0 | 7 |
| CHEMBL4847738 | C20H23N3O4 | 4.05 | 0 | 7 |
| CHEMBL3329814 | C21H28N2O5 | 4.09 | 0 | 7 |
| CHEMBL474484 | C13H21N7 | 4.28 | 0 | 7 |
| CHEMBL583 | C19H22FN3O3 | 4.3 | 0 | 7 |
| CHEMBL218836 | C18H24F3N3O | 4.4 | 0 | 7 |
| CHEMBL2069925 | C21H21N5OS | 4.52 | 0 | 7 |
| CHEMBL1957012 | C26H29N5O2 | 4.6 | 0 | 7 |
| CHEMBL575241 | C21H26N6S | 4.71 | 0 | 7 |
| CHEMBL2333636 | C24H28F4N2O | 4.75 | 0 | 7 |
| CHEMBL2180842 | C17H22FN5O | 4.8 | 0 | 7 |
| CHEMBL2314058 | C25H34F3N3O | 4.83 | 0 | 7 |
| CHEMBL3973288 | C20H22FN3O2S | 4.84 | 0 | 7 |
| CHEMBL523374 | C22H23N5O2 | 4.89 | 0 | 7 |
| CHEMBL3115194 | C26H32BrN3O3 | 4.9 | 0 | 7 |
| CHEMBL3218884 | C28H33N5O2 | 4.96 | 0 | 7 |
| CHEMBL2333607 | C25H30F4N2O | 5.05 | 0 | 7 |
| CHEMBL2333615 | C23H26F4N2O | 5.05 | 0 | 7 |
| CHEMBL498042 | C24H23N5O2 | 5.08 | 0 | 7 |
| CHEMBL3703271 | C22H23N3O4 | 5.08 | 0 | 7 |
| CHEMBL3262625 | C23H26N6O | 5.09 | 0 | 7 |
| CHEMBL246815 | C22H26FN3O3 | 5.13 | 0 | 7 |
| CHEMBL523821 | C21H15F2N5 | 5.16 | 0 | 7 |
| CHEMBL398478 | C25H28FN3O3 | 5.21 | 0 | 7 |
| CHEMBL2178522 | C23H19FN6 | 5.28 | 0 | 7 |
| CHEMBL560386 | C23H29NO5S | 5.31 | 0 | 7 |
| CHEMBL497048 | C27H23N5O2 | 5.36 | 0 | 7 |
| CHEMBL2147022 | C20H17N7 | 5.37 | 0 | 7 |
| CHEMBL2336331 | C24H29N5O2 | 5.4 | 0 | 7 |
| CHEMBL1813015 | C20H20N6O | 5.42 | 0 | 7 |

**Table S4** (cont.d)

| | | | | | |
|---|---|---|---|---|---|
| CHEMBL1939739 | C32H35FN4O2 | 5.44 | 0 | 7 | |
| CHEMBL2147021 | C20H15FN6 | 5.57 | 0 | 7 | |
| CHEMBL246634 | C21H23FN2O4 | 5.58 | 0 | 7 | |
| CHEMBL2151218 | C25H22FN5O | 5.7 | 0 | 7 | |
| CHEMBL3764774 | C28H28N4O3 | 5.7 | 0 | 7 | |
| CHEMBL243901 | C23H31N5OS | 5.9 | 0 | 7 | |
| CHEMBL17423 | C22H25N3O4 | 5.96 | 0 | 7 | |
| CHEMBL3775387 | C25H27N5O2 | 5.99 | 0 | 7 | |
| CHEMBL2387265 | C31H29FN2O4 | 6.09 | 0 | 7 | |
| CHEMBL1079578 | C28H31N5OS | 6.2 | 0 | 7 | |
| CHEMBL3775050 | C25H24F2N4O | 6.2 | 0 | 7 | |
| CHEMBL45816 | C29H38FN3O3 | 6.24 | 0 | 7 | |
| CHEMBL501480 | C29H25N5O2 | 6.5 | 0 | 7 | |
| CHEMBL1084400 | C20H21F3N4 | 6.8 | 0 | 7 | |
| CHEMBL584766 | C23H24FN5S | 7 | 0 | 7 | |
| CHEMBL526466 | C25H23N5O2 | 7.1 | 0 | 7 | |
| CHEMBL2146854 | C23H26N4O3 | 7.33 | 0 | 7 | |
| CHEMBL384487 | C24H23FN2O4 | 7.69 | 0 | 7 | |
| CHEMBL4279819 | C16H18N6O | 8.07 | 0 | 7 | |
| CHEMBL217593 | C23H23FN2O4 | 8.27 | 0 | 7 | |
| CHEMBL33 | C18H20FN3O4 | 3.04 | 0 | 8 | 5.03 |
| CHEMBL399726 | C20H24N6O2 | 3.78 | 0 | 8 | |
| CHEMBL31 | C19H22FN3O4 | 3.89 | 0 | 8 | |
| CHEMBL3804950 | C20H21F2N5O | 4.01 | 0 | 8 | |
| CHEMBL4858551 | C19H19FN6O | 4.01 | 0 | 8 | |
| CHEMBL520463 | C28H24N4O4 | 4.05 | 0 | 8 | |
| CHEMBL485123 | C20H23F3N4O | 4.07 | 0 | 8 | |
| CHEMBL32 | C21H24FN3O4 | 4.1 | 0 | 8 | |
| CHEMBL2333619 | C22H27F4N3O | 4.1 | 0 | 8 | |
| CHEMBL2180070 | C19H23N7O | 4.27 | 0 | 8 | |

**Table S4** (cont.d)

| | | | | |
|---|---|---|---|---|
| CHEMBL193 | C17H18N2O6 | 4.3 | 0 | 8 |
| CHEMBL256653 | C23H26N6OS | 4.3 | 0 | 8 |
| CHEMBL2382343 | C29H32F2N2O4 | 4.32 | 0 | 8 |
| CHEMBL2314064 | C24H32F3N3O2 | 4.41 | 0 | 8 |
| CHEMBL3577935 | C23H34F3N3O2 | 4.5 | 0 | 8 |
| CHEMBL3787345 | C25H28N4O4 | 4.51 | 0 | 8 |
| CHEMBL2164565 | C19H18F3N3O2 | 4.63 | 0 | 8 |
| CHEMBL602875 | C26H34N4O4 | 4.72 | 0 | 8 |
| CHEMBL1939742 | C30H37N5O2S | 4.75 | 0 | 8 |
| CHEMBL3612814 | C24H28N4O4 | 4.85 | 0 | 8 |
| CHEMBL3593771 | C19H19N7O | 4.87 | 0 | 8 |
| CHEMBL257901 | C28H30N6OS | 4.9 | 0 | 8 |
| CHEMBL270239 | C30H34N6OS | 4.9 | 0 | 8 |
| CHEMBL515001 | C21H25FN6O | 4.96 | 0 | 8 |
| CHEMBL1084617 | C24H28N4O4 | 4.96 | 0 | 8 |
| CHEMBL2440407 | C23H26N2O5S | 5.02 | 0 | 8 |
| CHEMBL465417 | C25H23N5O3 | 5.04 | 0 | 8 |
| CHEMBL3786346 | C25H26N4O4 | 5.05 | 0 | 8 |
| CHEMBL3287218 | C29H32N6OS | 5.16 | 0 | 8 |
| CHEMBL262341 | C33H29N7O | 5.25 | 0 | 8 |
| CHEMBL207220 | C24H40N4O3S | 5.4 | 0 | 8 |
| CHEMBL270852 | C32H38N6OS | 5.4 | 0 | 8 |
| CHEMBL271909 | C28H30N6S2 | 5.5 | 0 | 8 |
| CHEMBL2181489 | C23H20F3N3O2 | 5.52 | 0 | 8 |
| CHEMBL272086 | C32H38N6OS | 5.6 | 0 | 8 |
| CHEMBL2207738 | C23H26N4O3S | 5.6 | 0 | 8 |
| CHEMBL399525 | C19H23N7O | 5.75 | 0 | 8 |
| CHEMBL2178521 | C24H21FN6O | 5.75 | 0 | 8 |
| CHEMBL3415593 | C19H26N6O2 | 5.77 | 0 | 8 |
| CHEMBL1080489 | C22H27N5O2S | 5.8 | 0 | 8 |

**Table S4** (cont.d)

| | | | | | |
|---|---|---|---|---|---|
| CHEMBL478462 | C25H38N6O2 | 5.84 | 0 | 8 | |
| CHEMBL2441431 | C22H23N3O5 | 5.89 | 0 | 8 | |
| CHEMBL1621 | C23H27FN4O3 | 6 | 0 | 8 | |
| CHEMBL515025 | C24H36N6O2 | 6.15 | 0 | 8 | |
| CHEMBL478011 | C25H38N6O2 | 6.38 | 0 | 8 | |
| CHEMBL408169 | C28H32N6OS | 6.4 | 0 | 8 | |
| CHEMBL94454 | C24H25FN6O | 6.46 | 0 | 8 | |
| CHEMBL453894 | C25H26F3N3O2 | 6.46 | 0 | 8 | |
| CHEMBL244280 | C26H28FN5OS | 6.6 | 0 | 8 | |
| CHEMBL256442 | C29H32N6S2 | 6.8 | 0 | 8 | |
| CHEMBL568571 | C27H26F3N3O2 | 6.85 | 0 | 8 | |
| CHEMBL397429 | C28H30N6OS | 7.3 | 0 | 8 | |
| CHEMBL2146853 | C20H23N5O3 | 7.6 | 0 | 8 | |
| CHEMBL214021 | C23H21FN2O5 | 7.82 | 0 | 8 | |
| CHEMBL2324519 | C20H21FN6O2 | 3.67 | 0 | 9 | 5.11 |
| CHEMBL3799831 | C25H26FN3O5 | 4.03 | 0 | 9 | |
| CHEMBL3290351 | C24H27FN6O2 | 4.05 | 0 | 9 | |
| CHEMBL216323 | C23H19FN2O6 | 4.07 | 0 | 9 | |
| CHEMBL4092041 | C29H31F3N2O4 | 4.07 | 0 | 9 | |
| CHEMBL3290344 | C24H26N6O3 | 4.13 | 0 | 9 | |
| CHEMBL248296 | C27H31F2N3O4 | 4.19 | 0 | 9 | |
| CHEMBL2165068 | C25H26FN5O3 | 4.24 | 0 | 9 | |
| CHEMBL484816 | C22H21N7O2 | 4.3 | 0 | 9 | |
| CHEMBL3218891 | C27H32N6O3 | 4.37 | 0 | 9 | |
| CHEMBL2158050 | C25H29FN2O4S2 | 4.63 | 0 | 9 | |
| CHEMBL3650850 | C21H23N5O4 | 4.68 | 0 | 9 | |
| CHEMBL4084170 | C15H18F5N3S | 4.69 | 0 | 9 | |
| CHEMBL3612928 | C24H28N4O5 | 4.7 | 0 | 9 | |
| CHEMBL3422244 | C18H18F3N3O2S | 4.71 | 0 | 9 | |
| CHEMBL3786311 | C26H30N4O5 | 4.8 | 0 | 9 | |

**Table S4** (cont.d)

| | | | | | |
|---|---|---|---|---|---|
| CHEMBL596700 | C20H23F3N4O2 | 4.88 | 0 | 9 | |
| CHEMBL1083118 | C17H19F2N3O3S | 4.89 | 0 | 9 | |
| CHEMBL2441417 | C24H26N6O3 | 4.89 | 0 | 9 | |
| CHEMBL3416021 | C20H24N8O | 5.07 | 0 | 9 | |
| CHEMBL502288 | C23H22N6O2S | 5.11 | 0 | 9 | |
| CHEMBL3612926 | C25H30N4O5 | 5.11 | 0 | 9 | |
| CHEMBL2164393 | C26H24F3N3O3 | 5.14 | 0 | 9 | |
| CHEMBL2170611 | C17H21F3N4O2 | 5.16 | 0 | 9 | |
| CHEMBL402015 | C27H27BrN6OS | 5.2 | 0 | 9 | |
| CHEMBL429458 | C25H30FN3O5 | 5.2 | 0 | 9 | |
| CHEMBL4075071 | C19H16F3N5O | 5.25 | 0 | 9 | |
| CHEMBL4243499 | C19H16F3N5O | 5.25 | 0 | 9 | |
| CHEMBL2164389 | C24H20F3N3O3 | 5.34 | 0 | 9 | |
| CHEMBL1927161 | C24H29N5O4 | 5.4 | 0 | 9 | |
| CHEMBL244083 | C29H36N4O4S | 5.6 | 0 | 9 | |
| CHEMBL245119 | C25H31N7OS | 5.6 | 0 | 9 | |
| CHEMBL4213486 | C19H15F2N5O2 | 5.7 | 0 | 9 | |
| CHEMBL429761 | C23H31N7O2 | 5.73 | 0 | 9 | |
| CHEMBL3318999 | C28H25N7O2 | 5.85 | 0 | 9 | |
| CHEMBL446966 | C34H31N7O2 | 5.91 | 0 | 9 | |
| CHEMBL550410 | C25H26F4N4O | 5.92 | 0 | 9 | |
| CHEMBL514042 | C25H38N6O3 | 6.05 | 0 | 9 | |
| CHEMBL549635 | C29H33F4N3O2 | 6.2 | 0 | 9 | |
| CHEMBL217442 | C24H23FN2O6 | 7.8 | 0 | 9 | |
| CHEMBL387178 | C23H20F2N2O5 | 8.21 | 0 | 9 | |
| CHEMBL3422973 | C27H28N6O3 | 8.68 | 0 | 9 | |
| CHEMBL3422970 | C27H28N6O3 | 9.06 | 0 | 9 | |
| CHEMBL3921669 | C23H26N4O5S | 2.22 | 0 | 10 | 5.27 |
| CHEMBL2424928 | C24H26FN5O4 | 3.51 | 0 | 10 | |
| CHEMBL1916543 | C23H28N6O4 | 3.69 | 0 | 10 | |

**Table S4** (cont.d)

| | | | | |
|---|---|---|---|---|
| CHEMBL605785 | C27H27FN4O5 | 4.02 | 0 | 10 |
| CHEMBL4112037 | C16H14F2N6OS | 4.16 | 0 | 10 |
| CHEMBL2151322 | C19H20FN7O2 | 4.28 | 0 | 10 |
| CHEMBL3794265 | C23H25FN6O3 | 4.38 | 0 | 10 |
| CHEMBL247690 | C29H33F2N3O5 | 4.47 | 0 | 10 |
| CHEMBL3425929 | C28H33FN6O3 | 4.51 | 0 | 10 |
| CHEMBL2041188 | C32H30N6O4 | 4.64 | 0 | 10 |
| CHEMBL2324520 | C20H20F2N6O2 | 4.68 | 0 | 10 |
| CHEMBL2315921 | C26H28F3N5O2 | 4.77 | 0 | 10 |
| CHEMBL563791 | C28H29F3N6O | 4.9 | 0 | 10 |
| CHEMBL3400819 | C26H26FN5O4 | 4.91 | 0 | 10 |
| CHEMBL595944 | C27H26N8O2 | 4.98 | 0 | 10 |
| CHEMBL2164365 | C23H19F3N4O3 | 5.04 | 0 | 10 |
| CHEMBL2177736 | C19H23N9O | 5.07 | 0 | 10 |
| CHEMBL3681314 | C26H25F2N7O | 5.07 | 0 | 10 |
| CHEMBL513921 | C23H34N6O4 | 5.24 | 0 | 10 |
| CHEMBL3408394 | C37H24N6O2S2 | 5.24 | 0 | 10 |
| CHEMBL562285 | C24H27N3O2 | 5.27 | 0 | 10 |
| CHEMBL2164047 | C24H19F4N3O3 | 5.28 | 0 | 10 |
| CHEMBL245642 | C25H25F2N3O5 | 5.5 | 0 | 10 |
| CHEMBL551281 | C26H34F3N5O2 | 5.52 | 0 | 10 |
| CHEMBL1236904 | C26H31BrN8O | 5.52 | 0 | 10 |
| CHEMBL3604800 | C28H32FN5O4 | 5.52 | 0 | 10 |
| CHEMBL452823 | C37H38N10 | 5.67 | 0 | 10 |
| CHEMBL553196 | C32H32F4N4O2 | 5.76 | 0 | 10 |
| CHEMBL3323074 | C32H28FN7O2 | 5.85 | 0 | 10 |
| CHEMBL3323073 | C32H28FN7O2 | 6.2 | 0 | 10 |
| CHEMBL3605145 | C29H32FN5O4 | 6.3 | 0 | 10 |
| CHEMBL1081867 | C22H23F3N6S | 6.33 | 0 | 10 |
| CHEMBL3605140 | C27H30FN5O4 | 6.4 | 0 | 10 |

**Table S4** (cont.d)

| | | | | | |
|---|---|---|---|---|---|
| CHEMBL1081746 | C23H26F3N5S2 | 6.46 | 0 | 10 | |
| CHEMBL401576 | C25H26F3N5OS | 6.9 | 0 | 10 | |
| CHEMBL4081080 | C25H17F6N3O | 7.17 | 0 | 10 | |
| CHEMBL390649 | C28H31F3N6S | 7.5 | 0 | 10 | |
| CHEMBL428594 | C29H33F7N2O | 7.8 | 0 | 10 | |
| CHEMBL217707 | C23H19F3N2O5 | 8.25 | 0 | 10 | |
| CHEMBL473 | C19H27N3O5S2 | 8.39 | 0 | 10 | |
| CHEMBL213715 | C24H22F2N2O6 | 8.44 | 0 | 10 | |
| CHEMBL3985847 | C21H26N6O4S | 2.40 | 0 | 11 | 5 |
| CHEMBL1782574 | C23H35F3N4O3S | 2.41 | 0 | 11 | |
| CHEMBL3964789 | C21H22F4N6O | 3.17 | 0 | 11 | |
| CHEMBL3400817 | C26H26FN5O5 | 3.76 | 0 | 11 | |
| CHEMBL2165057 | C25H30N6O5 | 4.06 | 0 | 11 | |
| CHEMBL1916544 | C22H27N7O4 | 4.09 | 0 | 11 | |
| CHEMBL3703025 | C22H25F3N6O2 | 4.5 | 0 | 11 | |
| CHEMBL1091605 | C29H36F3N3O4S | 4.52 | 0 | 11 | |
| CHEMBL1091218 | C29H25F3N2O6 | 4.58 | 0 | 11 | |
| CHEMBL2069410 | C23H19F4N3O4 | 4.6 | 0 | 11 | |
| CHEMBL2204260 | C22H25F3N4O3S | 4.62 | 0 | 11 | |
| CHEMBL4209441 | C26H23FN8O2 | 4.62 | 0 | 11 | |
| CHEMBL4211893 | C30H29FN8O2 | 4.87 | 0 | 11 | |
| CHEMBL2164375 | C25H26F3N3O5 | 4.96 | 0 | 11 | |
| CHEMBL2204270 | C24H26F4N4O3 | 5 | 0 | 11 | |
| CHEMBL3318984 | C23H23F3N6O2 | 5 | 0 | 11 | |
| CHEMBL508098 | C36H37N11 | 5.08 | 0 | 11 | |
| CHEMBL3746204 | C22H17F4N5O2 | 5.24 | 0 | 11 | |
| CHEMBL2331648 | C20H22F3N5O3 | 5.34 | 0 | 11 | |
| CHEMBL4075908 | C20H21BrFN5O3S | 5.46 | 0 | 11 | |
| CHEMBL192 | C22H30N6O4S | 5.48 | 0 | 11 | |
| CHEMBL3422758 | C27H29F2N5O4 | 5.76 | 0 | 11 | |

**Table S4** (cont.d)

| | | | | | |
|---|---|---|---|---|---|
| CHEMBL2041175 | C31H27N7O3S | 5.82 | 0 | 11 | |
| CHEMBL256154 | C27H32F6N4O | 5.85 | 0 | 11 | |
| CHEMBL1082111 | C22H23F4N5OS | 5.9 | 0 | 11 | |
| CHEMBL402016 | C28H27F3N6OS | 6 | 0 | 11 | |
| CHEMBL3605123 | C30H36FN5O5 | 6.16 | 0 | 11 | |
| CHEMBL411293 | C28H28F3N7S | 6.6 | 0 | 11 | |
| CHEMBL3422952 | C26H27F2N5O4 | 9.14 | 0 | 11 | |
| CHEMBL3422978 | C26H26F3N5O3 | 9.41 | 0 | 11 | |
| CHEMBL491571 | C15H24N4O6S2 | 2.1 | 0 | 12 | 4.59 |
| CHEMBL254316 | C20H21FN6O5 | 2.5 | 0 | 12 | |
| CHEMBL3425799 | C26H25F2N5O5 | 3.48 | 0 | 12 | |
| CHEMBL1784523 | C36H40F3N5O3S | 4 | 0 | 12 | |
| CHEMBL3417745 | C23H25F3N2O6S | 4 | 0 | 12 | |
| CHEMBL451887 | C40H57N5O7 | 4.04 | 0 | 12 | |
| CHEMBL3342693 | C23H22F5N5O2 | 4.08 | 0 | 12 | |
| CHEMBL3425807 | C25H25FN6O5 | 4.52 | 0 | 12 | |
| CHEMBL2206791 | C23H21F3N6O3 | 4.58 | 0 | 12 | |
| CHEMBL2147316 | C22H19F5N4O3 | 4.6 | 0 | 12 | |
| CHEMBL2336323 | C26H30F2N6O4 | 4.62 | 0 | 12 | |
| CHEMBL2147303 | C22H19F5N4O3 | 4.7 | 0 | 12 | |
| CHEMBL402624 | C30H34F6N2O4 | 4.71 | 0 | 12 | |
| CHEMBL3605131 | C29H32FN5O6 | 4.91 | 0 | 12 | |
| CHEMBL3288030 | C44H53FN6O5 | 5.2 | 0 | 12 | |
| CHEMBL2441412 | C24H23F3N6O3 | 5.29 | 0 | 12 | |
| CHEMBL550535 | C27H33F6N5O | 5.41 | 0 | 12 | |
| CHEMBL3605006 | C31H34FN5O6 | 5.8 | 0 | 12 | |
| CHEMBL195378 | C23H24F4N6O3 | 4.23 | 0 | 13 | 5.29 |
| CHEMBL3605022 | C29H32FN7O5 | 4.54 | 0 | 13 | |
| CHEMBL370072 | C24H24F4N4O5 | 4.55 | 0 | 13 | |
| CHEMBL3218816 | C19H17F3N8O2 | 4.75 | 0 | 13 | |

**Table S4** (cont.d)

| | | | | | |
|---|---|---|---|---|---|
| CHEMBL1099069 | C19H24F3N9O | 5.29 | 0 | 13 | |
| CHEMBL4079820 | C26H23F4N7O2 | 5.62 | 0 | 13 | |
| CHEMBL490026 | C24H24F6N2O4S | 6.05 | 0 | 13 | |
| CHEMBL4077588 | C29H29F4N7O2 | 7 | 0 | 13 | |
| CHEMBL3422957 | C27H27F4N5O4 | 8.77 | 0 | 13 | |
| CHEMBL231396 | C20H17F6N5O3 | 4 | 0 | 14 | |
| CHEMBL4097247 | C27H26F3N7O4 | 5.24 | 0 | 14 | |
| CHEMBL435882 | C25H26F7N3O4 | 6.06 | 0 | 14 | |
| CHEMBL487708 | C25H22F6N4O3S | 6.3 | 0 | 14 | |
| CHEMBL1908360 | C53H83NO14 | 3.3 | 0 | 15 | |
| CHEMBL3393730 | C42H52N8O6S2 | 4.6 | 0 | 16 | |
| CHEMBL1684059 | C20H28ClN | 5.37 | 1 | 2 | |
| CHEMBL3314420 | C15H13ClFN | 4.3 | 1 | 3 | 5.6 |
| CHEMBL1945694 | C16H15ClN2 | 5.16 | 1 | 3 | |
| CHEMBL191054 | C13H17ClN2 | 5.6 | 1 | 3 | |
| CHEMBL299390 | C19H19ClFN | 5.83 | 1 | 3 | |
| CHEMBL505 | C16H19ClN2 | 5.96 | 1 | 3 | |
| CHEMBL1083407 | C21H22ClN3 | 5.44 | 1 | 4 | |
| CHEMBL53866 | C17H15ClFNO | 5.66 | 1 | 4 | |
| CHEMBL3696120 | C22H26ClN3 | 5.96 | 1 | 4 | |
| CHEMBL384400 | C27H36ClN3O | 5.48 | 1 | 5 | 6.72 |
| CHEMBL562833 | C22H30ClN3O | 6.1 | 1 | 5 | |
| CHEMBL42 | C18H19ClN4 | 6.72 | 1 | 5 | |
| CHEMBL998 | C22H23ClN2O2 | 6.77 | 1 | 5 | |
| CHEMBL195180 | C24H27ClN2OS | 7.52 | 1 | 5 | |
| CHEMBL4787638 | C21H18ClN3O2 | 2.08 | 1 | 6 | 5.05 |
| CHEMBL1688 | C18H19ClN4O | 3.85 | 1 | 6 | |
| CHEMBL1000 | C21H25ClN2O3 | 4.52 | 1 | 6 | |
| CHEMBL494406 | C21H24ClN5 | 5.05 | 1 | 6 | |
| CHEMBL2146805 | C20H21ClN4O | 5.6 | 1 | 6 | |

**Table S4** (cont.d)

| | | | | | |
|---|---|---|---|---|---|
| CHEMBL563926 | C21H26ClN3O2 | 6.2 | 1 | 6 | |
| CHEMBL2146810 | C23H27ClN4O | 8.4 | 1 | 6 | |
| CHEMBL4781551 | C24H21ClN4O2 | 4.08 | 1 | 7 | 5.45 |
| CHEMBL4127458 | C22H22ClN5O | 4.22 | 1 | 7 | |
| CHEMBL512975 | C21H23ClFN5 | 4.64 | 1 | 7 | |
| CHEMBL2333616 | C20H22ClF3N2O | 4.92 | 1 | 7 | |
| CHEMBL1957792 | C27H30ClN3O3 | 4.99 | 1 | 7 | |
| CHEMBL1092106 | C20H24ClFN4O | 5.04 | 1 | 7 | |
| CHEMBL1091834 | C25H26ClFN4O | 5.42 | 1 | 7 | |
| CHEMBL250533 | C25H28ClN3O3 | 5.48 | 1 | 7 | |
| CHEMBL4102707 | C20H20ClF4NO | 5.54 | 1 | 7 | |
| CHEMBL3086038 | C30H33BrClN3O2 | 5.82 | 1 | 7 | |
| CHEMBL2146812 | C19H20ClN5O | 5.9 | 1 | 7 | |
| CHEMBL3806153 | C18H21ClFN3O2 | 6.01 | 1 | 7 | |
| CHEMBL708 | C21H21ClN4OS | 6.92 | 1 | 7 | |
| CHEMBL12713 | C24H26ClFN4O | 8.57 | 1 | 7 | |
| CHEMBL2178221 | C28H29ClN4O3 | 4.82 | 1 | 8 | 5.8 |
| CHEMBL1093696 | C24H25ClFN5O | 4.92 | 1 | 8 | |
| CHEMBL2325997 | C20H23ClN6O | 5.14 | 1 | 8 | |
| CHEMBL506163 | C22H19ClFN5O | 5.2 | 1 | 8 | |
| CHEMBL3218650 | C27H30ClN5O2 | 5.6 | 1 | 8 | |
| CHEMBL3703237 | C19H17ClN4O3 | 5.66 | 1 | 8 | |
| CHEMBL2387229 | C26H26ClFN2O4 | 5.8 | 1 | 8 | |
| CHEMBL1956991 | C23H26ClFN4O2 | 5.89 | 1 | 8 | |
| CHEMBL3318989 | C23H24ClN5O2 | 6 | 1 | 8 | |
| CHEMBL2146813 | C38H48ClN5O2 | 6.8 | 1 | 8 | |
| CHEMBL525413 | C29H24ClN5O2 | 7.09 | 1 | 8 | |
| CHEMBL244278 | C26H28ClN5OS | 7.1 | 1 | 8 | |
| CHEMBL217789 | C23H21ClN2O5 | 8.44 | 1 | 8 | |
| CHEMBL3608687 | C16H15ClN6OS | 4.18 | 1 | 9 | 5.68 |

**Table S4** (cont.d)

| | | | | | |
|---|---|---|---|---|---|
| CHEMBL4094682 | C22H27ClN6O2 | 4.34 | 1 | 9 | |
| CHEMBL2325729 | C26H34ClN7O | 4.53 | 1 | 9 | |
| CHEMBL3262358 | C21H22ClN7O | 4.7 | 1 | 9 | |
| CHEMBL255968 | C25H24ClN5O3 | 5 | 1 | 9 | |
| CHEMBL1091664 | C28H25ClN2O5S | 5 | 1 | 9 | |
| CHEMBL489324 | C28H42ClF2N5O | 5.68 | 1 | 9 | |
| CHEMBL3237444 | C23H25ClN8 | 5.7 | 1 | 9 | |
| CHEMBL2177305 | C28H23ClFN5O2 | 5.89 | 1 | 9 | |
| CHEMBL557306 | C21H24ClF3N2O3 | 5.99 | 1 | 9 | |
| CHEMBL1081747 | C21H23ClFN5OS | 6.4 | 1 | 9 | |
| CHEMBL3983509 | C20H20ClFN4O2S | 6.7 | 1 | 9 | |
| CHEMBL1729 | C23H29ClFN3O4 | 8.19 | 1 | 9 | |
| CHEMBL2440387 | C25H23ClN4O4S | 3.38 | 1 | 10 | 4.73 |
| CHEMBL3221500 | C22H21ClN6O3 | 4.2 | 1 | 10 | |
| CHEMBL3221488 | C23H23ClN6O3 | 4.5 | 1 | 10 | |
| CHEMBL256060 | C26H26ClN5O4 | 4.96 | 1 | 10 | |
| CHEMBL2403108 | C28H36ClN5O3S | 5.9 | 1 | 10 | |
| CHEMBL375330 | C34H38ClF3N4O2 | 6.77 | 1 | 10 | |
| CHEMBL3221490 | C23H22ClN7O3 | 4.4 | 1 | 11 | 4.95 |
| CHEMBL3407784 | C20H19ClN6O3S | 4.89 | 1 | 11 | |
| CHEMBL3221503 | C23H20ClN7O3 | 4.9 | 1 | 11 | |
| CHEMBL4076061 | C22H20ClF2N3O3S2 | 4.99 | 1 | 11 | |
| CHEMBL3353404 | C26H29ClN8O2 | 5.17 | 1 | 11 | |
| CHEMBL2147223 | C22H20ClF3N4O3 | 5.4 | 1 | 11 | |
| CHEMBL3608744 | C28H31ClN6O4S | 4.09 | 1 | 12 | |
| CHEMBL3221487 | C20H19ClN8O3 | 4.8 | 1 | 12 | |
| CHEMBL2064657 | C28H23ClF2N6O3 | 4.91 | 1 | 12 | |
| CHEMBL401608 | C27H31ClF6N4O2 | 6.73 | 1 | 13 | |
| CHEMBL1684047 | C16H23Cl2N | 4.4 | 2 | 3 | |
| CHEMBL2324239 | C24H18Cl2NO2$^{+}$ | 4.51 | 2 | 5 | |

**Table S4** (cont.d)

| | | | | | |
|---|---|---|---|---|---|
| CHEMBL1080790 | C16H22Cl2N2O | 4.78 | 2 | 5 | |
| CHEMBL1080969 | C16H22Cl2N2O | 5.11 | 2 | 5 | |
| CHEMBL2324245 | $C24H26Cl2NO2^{+}$ | 7.22 | 2 | 5 | |
| CHEMBL3741589 | C18H17Cl2NO3 | 4.11 | 2 | 6 | |
| CHEMBL1083463 | C26H26Cl2N4 | 6.22 | 2 | 6 | |
| CHEMBL2158823 | C24H28Cl2N2O3 | 4.5 | 2 | 7 | |
| CHEMBL2158815 | C24H28Cl2N2O3 | 5.3 | 2 | 7 | |
| CHEMBL3806193 | C19H23Cl2N3O2 | 5.84 | 2 | 7 | |
| CHEMBL1107 | C26H30Cl2F3NO | 7.4 | 2 | 7 | |
| CHEMBL540634 | C26H31Cl2N3O3 | 4.5 | 2 | 8 | 5.15 |
| CHEMBL3216126 | C25H35Cl2N5O | 4.65 | 2 | 8 | |
| CHEMBL3964723 | C23H27Cl2N3O3 | 5 | 2 | 8 | |
| CHEMBL2158771 | C27H31Cl2N3O3 | 5.3 | 2 | 8 | |
| CHEMBL233047 | C20H32Cl2N6 | 6.52 | 2 | 8 | |
| CHEMBL128007 | C28H28Cl2N4O2 | 7 | 2 | 8 | |
| CHEMBL589833 | C24H25Cl2FN4O2 | 5.1 | 2 | 9 | 5.36 |
| CHEMBL4170966 | C24H21Cl2N3O4 | 5.33 | 2 | 9 | |
| CHEMBL255645 | C28H31Cl2N5O2 | 5.36 | 2 | 9 | |
| CHEMBL2158625 | C20H19Cl2N5OS | 5.44 | 2 | 9 | |
| CHEMBL2158629 | C20H21Cl2N5OS | 5.85 | 2 | 9 | |
| CHEMBL560063 | C24H23Cl2F3N4O | 5.46 | 2 | 10 | |
| CHEMBL537847 | C23H29Cl2N5O3 | 6.25 | 2 | 10 | |
| CHEMBL3764895 | C29H31Cl2F3N4O3 | 5.13 | 2 | 12 | |
| CHEMBL3764357 | C29H31Cl2F3N4O3 | 5.2 | 2 | 12 | |
| CHEMBL2207664 | C22H24Cl2F3N3O4 | 5.3 | 2 | 12 | |
| CHEMBL3765012 | C30H31Cl2F3N4O3 | 5.51 | 2 | 12 | |
| CHEMBL1098847 | C24H21Cl3N2O2 | 5.58 | 3 | 7 | |
| CHEMBL1095476 | C28H25Cl3N4O2 | 5.06 | 3 | 9 | |
| CHEMBL498411 | C22H19Cl3N6 | 6.46 | 3 | 9 | |
| CHEMBL1097456 | C26H22Cl3F3N2O3 | 5.84 | 3 | 11 | |

**Table S5.** Investigation of the influence of oxygen atom on the $pIC_{50}$ value.

| Name | Molecular formula | pIC50 | O | Hetatoms | Median |
|---|---|---|---|---|---|
| CHEMBL284348 | C5H6N2 | 2.36 | 0 | 2 | 5.11 |
| CHEMBL3 | C10H14N2 | 3.61 | 0 | 2 | |
| CHEMBL1086033 | C17H22N2 | 3.67 | 0 | 2 | |
| CHEMBL87045 | C18H18N2 | 5.06 | 0 | 2 | |
| CHEMBL459176 | C20H22N2 | 5.16 | 0 | 2 | |
| CHEMBL1684059 | C20H28ClN | 5.37 | 0 | 2 | |
| CHEMBL11 | C19H24N2 | 5.47 | 0 | 2 | |
| CHEMBL1671894 | C27H26N2 | 7.75 | 0 | 2 | |
| CHEMBL3314420 | C15H13ClFN | 4.30 | 0 | 3 | 5.80 |
| CHEMBL1684047 | C16H23Cl2N | 4.40 | 0 | 3 | |
| CHEMBL513258 | C19H19FN2 | 5.05 | 0 | 3 | |
| CHEMBL1945694 | C16H15ClN2 | 5.16 | 0 | 3 | |
| CHEMBL191054 | C13H17ClN2 | 5.60 | 0 | 3 | |
| CHEMBL497171 | C15H17N3 | 5.80 | 0 | 3 | |
| CHEMBL299390 | C19H19ClFN | 5.83 | 0 | 3 | |
| CHEMBL457930 | C20H19N3 | 5.85 | 0 | 3 | |
| CHEMBL505 | C16H19ClN2 | 5.96 | 0 | 3 | |
| CHEMBL1090528 | C19H24N2S | 6.21 | 0 | 3 | |
| CHEMBL299499 | C17H22IN3 | 6.31 | 0 | 3 | |
| CHEMBL1671896 | C11H14F3N | 3.27 | 0 | 4 | 5.70 |
| CHEMBL3314426 | C15H14F2N2 | 4.66 | 0 | 4 | |
| CHEMBL1083407 | C21H22ClN3 | 5.44 | 0 | 4 | |
| CHEMBL1093058 | C17H21N3S | 5.70 | 0 | 4 | |
| CHEMBL3696120 | C22H26ClN3 | 5.96 | 0 | 4 | |
| CHEMBL1092377 | C19H21FN2S | 6.48 | 0 | 4 | |
| CHEMBL2046892 | C22H22F2N2 | 7.40 | 0 | 4 | |
| CHEMBL1125 | C16H18N4S | 4.90 | 0 | 5 | 5.33 |
| CHEMBL237191 | C25H31N5 | 4.98 | 0 | 5 | |
| CHEMBL2326478 | C18H18FN3S | 5.17 | 0 | 5 | |
| CHEMBL1956112 | C21H28N4S | 5.33 | 0 | 5 | |

**Table S5** (Cont.d)

| | | | | | |
|---|---|---|---|---|---|
| CHEMBL42 | C18H19ClN4 | 6.72 | 0 | 5 | |
| CHEMBL715 | C17H20N4S | 6.74 | 0 | 5 | |
| CHEMBL61301 | C19H21FN4 | 7.56 | 0 | 5 | |
| CHEMBL460491 | C21H34N4S2 | 4.88 | 0 | 6 | 5.80 |
| CHEMBL494406 | C21H24ClN5 | 5.05 | 0 | 6 | |
| CHEMBL2177904 | C23H16FN5 | 5.51 | 0 | 6 | |
| CHEMBL2177905 | C24H16F2N4 | 5.80 | 0 | 6 | |
| CHEMBL2147024 | C22H18N6 | 5.96 | 0 | 6 | |
| CHEMBL1083463 | C26H26Cl2N4 | 6.22 | 0 | 6 | |
| CHEMBL522456 | CHEMBL522456 | 7.00 | 0 | 6 | |
| CHEMBL474484 | C13H21N7 | 4.28 | 0 | 7 | 5.28 |
| CHEMBL512975 | C21H23ClFN5 | 4.64 | 0 | 7 | |
| CHEMBL575241 | C21H26N6S | 4.71 | 0 | 7 | |
| CHEMBL523821 | C21H15F2N5 | 5.16 | 0 | 7 | |
| CHEMBL2178522 | C23H19FN6 | 5.28 | 0 | 7 | |
| CHEMBL2147022 | C20H17N7 | 5.37 | 0 | 7 | |
| CHEMBL2147021 | C20H15FN6 | 5.57 | 0 | 7 | |
| CHEMBL1084400 | C20H21F3N4 | 6.80 | 0 | 7 | |
| CHEMBL584766 | C23H24FN5S | 7.00 | 0 | 7 | |
| CHEMBL640 | C13H21N3O | 3.86 | 1 | 4 | 4.91 |
| CHEMBL219803 | C17H25N3O | 3.94 | 1 | 4 | |
| CHEMBL219128 | C17H33N3O | 4.17 | 1 | 4 | |
| CHEMBL219074 | C20H31N3O | 4.42 | 1 | 4 | |
| CHEMBL1086273 | C19H23FN2O | 4.68 | 1 | 4 | |
| CHEMBL1089159 | C22H23N3O | 4.70 | 1 | 4 | |
| CHEMBL2333646 | C19H23BrN2O | 4.89 | 1 | 4 | |
| CHEMBL517 | C21H29N3O | 4.92 | 1 | 4 | |
| CHEMBL562948 | C22H33N3O | 5.40 | 1 | 4 | |
| CHEMBL1090175 | C20H25N3O | 5.47 | 1 | 4 | |
| CHEMBL53866 | C17H15ClFNO | 5.66 | 1 | 4 | |

**Table S5** (cont.d)

| | | | | | |
|---|---|---|---|---|---|
| CHEMBL1935443 | C20H23N3O | 5.68 | 1 | 4 | |
| CHEMBL3323181 | C22H24N2OS | 6.00 | 1 | 4 | |
| CHEMBL460893 | C23H25N3O | 6.13 | 1 | 4 | |
| CHEMBL4213196 | C25H34N4O | 4.62 | 1 | 5 | 5.20 |
| CHEMBL217133 | C18H22N4O | 4.68 | 1 | 5 | |
| CHEMBL429115 | C20H24N4O | 4.71 | 1 | 5 | |
| CHEMBL1080790 | C16H22Cl2N2O | 4.78 | 1 | 5 | |
| CHEMBL4096683 | C20H22N4O | 4.94 | 1 | 5 | |
| CHEMBL565829 | C15H14N4O | 5.01 | 1 | 5 | |
| CHEMBL2177304 | C27H24FN3O | 5.07 | 1 | 5 | |
| CHEMBL1080969 | C16H22Cl2N2O | 5.11 | 1 | 5 | |
| CHEMBL4100591 | C27H34F2N2O | 5.29 | 1 | 5 | |
| CHEMBL384400 | C27H36ClN3O | 5.48 | 1 | 5 | |
| CHEMBL1110 | C17H18N4O | 5.64 | 1 | 5 | |
| CHEMBL562833 | C22H30ClN3O | 6.10 | 1 | 5 | |
| CHEMBL58387 | C31H38N4O | 6.24 | 1 | 5 | |
| CHEMBL3775729 | C26H28N4O | 6.28 | 1 | 5 | |
| CHEMBL1087493 | C17H26N4O | 6.63 | 1 | 5 | |
| CHEMBL195180 | C24H27ClN2OS | 7.52 | 1 | 5 | |
| CHEMBL1688 | C18H19ClN4O | 3.85 | 1 | 6 | 5.96 |
| CHEMBL4212656 | C24H31N5O | 4.64 | 1 | 6 | |
| CHEMBL398612 | C26H35FN4O | 4.88 | 1 | 6 | |
| CHEMBL4067770 | C20H21FN4O | 5.11 | 1 | 6 | |
| CHEMBL4100776 | C20H22N4OS | 5.23 | 1 | 6 | |
| CHEMBL225036 | C24H26N4OS | 5.29 | 1 | 6 | |
| CHEMBL2146805 | C20H21ClN4O | 5.60 | 1 | 6 | |
| CHEMBL4104102 | C20H22N4OS | 5.61 | 1 | 6 | |
| CHEMBL4081790 | C25H19N5O | 6.30 | 1 | 6 | |
| CHEMBL4096145 | C22H27N5O | 6.52 | 1 | 6 | |

**Table S5** (cont.d)

| | | | | | |
|---|---|---|---|---|---|
| CHEMBL3774817 | C25H25FN4O | 6.59 | 1 | 6 | |
| CHEMBL329067 | C32H33N5O | 7.62 | 1 | 6 | |
| CHEMBL12780 | C24H27FN4O | 8.16 | 1 | 6 | |
| CHEMBL2146810 | C23H27ClN4O | 8.40 | 1 | 6 | |
| CHEMBL1423 | C28H29F2N3O | 8.52 | 1 | 6 | |
| CHEMBL296419 | C28H31FN4O | 9.05 | 1 | 6 | |
| CHEMBL578834 | C21H25N5OS | 3.72 | 1 | 7 | 5.07 |
| CHEMBL3582294 | C25H18F2N4O | 4.03 | 1 | 7 | |
| CHEMBL4224807 | C14H16N6O | 4.03 | 1 | 7 | |
| CHEMBL4127458 | C22H22ClN5O | 4.22 | 1 | 7 | |
| CHEMBL218836 | C18H24F3N3O | 4.40 | 1 | 7 | |
| CHEMBL2069925 | C21H21N5OS | 4.52 | 1 | 7 | |
| CHEMBL2333636 | C24H28F4N2O | 4.75 | 1 | 7 | |
| CHEMBL2180842 | C17H22FN5O | 4.80 | 1 | 7 | |
| CHEMBL2314058 | C25H34F3N3O | 4.83 | 1 | 7 | |
| CHEMBL2333616 | C20H22ClF3N2O | 4.92 | 1 | 7 | |
| CHEMBL1092106 | C20H24ClFN4O | 5.04 | 1 | 7 | |
| CHEMBL2333607 | C25H30F4N2O | 5.05 | 1 | 7 | |
| CHEMBL2333615 | C23H26F4N2O | 5.05 | 1 | 7 | |
| CHEMBL3262625 | C23H26N6O | 5.09 | 1 | 7 | |
| CHEMBL1091834 | C25H26ClFN4O | 5.42 | 1 | 7 | |
| CHEMBL1813015 | C20H20N6O | 5.42 | 1 | 7 | |
| CHEMBL4102707 | C20H20ClF4NO | 5.54 | 1 | 7 | |
| CHEMBL2151218 | C25H22FN5O | 5.70 | 1 | 7 | |
| CHEMBL243901 | C23H31N5OS | 5.90 | 1 | 7 | |
| CHEMBL2146812 | C19H20ClN5O | 5.90 | 1 | 7 | |
| CHEMBL1079578 | C28H31N5OS | 6.20 | 1 | 7 | |
| CHEMBL3775050 | C25H24F2N4O | 6.20 | 1 | 7 | |
| CHEMBL708 | C21H21ClN4OS | 6.92 | 1 | 7 | |
| CHEMBL1107 | C26H30Cl2F3NO | 7.40 | 1 | 7 | |
| CHEMBL4279819 | C16H18N6O | 8.07 | 1 | 7 | |

**Table S5** (cont.d)

| | | | | | |
|---|---|---|---|---|---|
| CHEMBL12713 | C24H26ClFN4O | 8.57 | 1 | 7 | |
| CHEMBL3804950 | C20H21F2N5O | 4.01 | 1 | 8 | 5.14 |
| CHEMBL4858551 | C19H19FN6O | 4.01 | 1 | 8 | |
| CHEMBL485123 | C20H23F3N4O | 4.07 | 1 | 8 | |
| CHEMBL2333619 | C22H27F4N3O | 4.10 | 1 | 8 | |
| CHEMBL2180070 | C19H23N7O | 4.27 | 1 | 8 | |
| CHEMBL256653 | C23H26N6OS | 4.30 | 1 | 8 | |
| CHEMBL3216126 | C25H35Cl2N5O | 4.65 | 1 | 8 | |
| CHEMBL3593771 | C19H19N7O | 4.87 | 1 | 8 | |
| CHEMBL257901 | C28H30N6OS | 4.90 | 1 | 8 | |
| CHEMBL270239 | C30H34N6OS | 4.90 | 1 | 8 | |
| CHEMBL1093696 | C24H25ClFN5O | 4.92 | 1 | 8 | |
| CHEMBL515001 | C21H25FN6O | 4.96 | 1 | 8 | |
| CHEMBL2325997 | C20H23ClN6O | 5.14 | 1 | 8 | |
| CHEMBL3287218 | C29H32N6OS | 5.16 | 1 | 8 | |
| CHEMBL506163 | C22H19ClFN5O | 5.20 | 1 | 8 | |
| CHEMBL262341 | C33H29N7O | 5.25 | 1 | 8 | |
| CHEMBL270852 | C32H38N6OS | 5.40 | 1 | 8 | |
| CHEMBL272086 | C32H38N6OS | 5.60 | 1 | 8 | |
| CHEMBL399525 | C19H23N7O | 5.75 | 1 | 8 | |
| CHEMBL2178521 | C24H21FN6O | 5.75 | 1 | 8 | |
| CHEMBL408169 | C28H32N6OS | 6.40 | 1 | 8 | |
| CHEMBL94454 | C24H25FN6O | 6.46 | 1 | 8 | |
| CHEMBL244280 | C26H28FN5OS | 6.60 | 1 | 8 | |
| CHEMBL244278 | C26H28ClN5OS | 7.10 | 1 | 8 | |
| CHEMBL397429 | C28H30N6OS | 7.30 | 1 | 8 | |
| CHEMBL3608687 | C16H15ClN6OS | 4.18 | 1 | 9 | 5.35 |
| CHEMBL2325729 | C26H34ClN7O | 4.53 | 1 | 9 | |
| CHEMBL3262358 | C21H22ClN7O | 4.70 | 1 | 9 | |
| CHEMBL3416021 | C20H24N8O | 5.07 | 1 | 9 | |
| CHEMBL402015 | C27H27BrN6OS | 5.20 | 1 | 9 | |

**Table S5** (cont.d)

| | | | | | |
|---|---|---|---|---|---|
| CHEMBL4075071 | C19H16F3N5O | 5.25 | 1 | 9 | |
| CHEMBL2158625 | C20H19Cl2N5OS | 5.44 | 1 | 9 | |
| CHEMBL245119 | C25H31N7OS | 5.60 | 1 | 9 | |
| CHEMBL489324 | C28H42ClF2N5O | 5.68 | 1 | 9 | |
| CHEMBL2158629 | C20H21Cl2N5OS | 5.85 | 1 | 9 | |
| CHEMBL550410 | C25H26F4N4O | 5.92 | 1 | 9 | |
| CHEMBL1081747 | C21H23ClFN5OS | 6.40 | 1 | 9 | |
| CHEMBL4112037 | C16H14F2N6OS | 4.16 | 1 | 10 | 5.37 |
| CHEMBL563791 | C28H29F3N6O | 4.90 | 1 | 10 | |
| CHEMBL2177736 | C19H23N9O | 5.07 | 1 | 10 | |
| CHEMBL3681314 | C26H25F2N7O | 5.07 | 1 | 10 | |
| CHEMBL562285 | C24H27N3O2 | 5.27 | 1 | 10 | |
| CHEMBL560063 | C24H23Cl2F3N4O | 5.46 | 1 | 10 | |
| CHEMBL1236904 | C26H31BrN8O | 5.52 | 1 | 10 | |
| CHEMBL401576 | C25H26F3N5OS | 6.90 | 1 | 10 | |
| CHEMBL4081080 | C25H17F6N3O | 7.17 | 1 | 10 | |
| CHEMBL428594 | C29H33F7N2O | 7.80 | 1 | 10 | |
| CHEMBL16 | C15H12N2O2 | 4.00 | 2 | 4 | 5.92 |
| CHEMBL3356251 | C24H36N2O2 | 4.01 | 2 | 4 | |
| CHEMBL3786542 | C26H26N2O2 | 4.21 | 2 | 4 | |
| CHEMBL3356249 | C25H38N2O2 | 4.30 | 2 | 4 | |
| CHEMBL3185736 | C22H20N2O2 | 4.91 | 2 | 4 | |
| CHEMBL4162513 | C35H34N2O2 | 5.10 | 2 | 4 | |
| CHEMBL4450665 | C23H28N2O2 | 5.92 | 2 | 4 | |
| CHEMBL1346 | C28H30N2O2 | 7.10 | 2 | 4 | |
| CHEMBL1223951 | C21H24N2O2 | 9.21 | 2 | 4 | |
| CHEMBL1257938 | C22H26N2O2 | 9.21 | 2 | 4 | |
| CHEMBL1257937 | C22H26N2O2 | 9.28 | 2 | 4 | |
| CHEMBL1257820 | C23H28N2O2 | 9.60 | 2 | 4 | |
| CHEMBL1257821 | C22H26N2O2 | 9.85 | 2 | 4 | |
| CHEMBL1097 | C15H23N3O2 | 4.00 | 2 | 5 | 5.18 |

**Table S5** (cont.d)

| | | | | | |
|---|---|---|---|---|---|
| CHEMBL4209354 | C15H20N2O2S | 4.07 | 2 | 5 | |
| CHEMBL1091879 | C19H25N3O2 | 4.12 | 2 | 5 | |
| CHEMBL561138 | C23H27N3O2 | 4.14 | 2 | 5 | |
| CHEMBL1258006 | C22H23N3O2 | 4.27 | 2 | 5 | |
| CHEMBL2324239 | $C24H18Cl2NO2^{+}$ | 4.51 | 2 | 5 | |
| CHEMBL1094041 | C29H29N3O2 | 4.55 | 2 | 5 | |
| CHEMBL2441636 | C22H33N3O2 | 4.72 | 2 | 5 | |
| CHEMBL487063 | C23H27N3O2 | 5.10 | 2 | 5 | |
| CHEMBL378547 | C21H21F2NO2 | 5.11 | 2 | 5 | |
| CHEMBL485242 | C25H29N3O2 | 5.14 | 2 | 5 | |
| CHEMBL478615 | C23H27N3O2 | 5.17 | 2 | 5 | |
| CHEMBL476579 | C24H29N3O2 | 5.18 | 2 | 5 | |
| CHEMBL479242 | C23H27N3O2 | 5.18 | 2 | 5 | |
| CHEMBL3121096 | C20H25N3O2 | 5.22 | 2 | 5 | |
| CHEMBL1819137 | C22H25N3O2 | 5.23 | 2 | 5 | |
| CHEMBL488249 | C24H27N3O2 | 5.25 | 2 | 5 | |
| CHEMBL487064 | C23H27N3O2 | 5.55 | 2 | 5 | |
| CHEMBL478616 | C23H27N3O2 | 5.80 | 2 | 5 | |
| CHEMBL1224697 | C22H28N2O2S | 5.90 | 2 | 5 | |
| CHEMBL376488 | C32H31BrN2O2 | 6.43 | 2 | 5 | |
| CHEMBL998 | C22H23ClN2O2 | 6.77 | 2 | 5 | |
| CHEMBL2324245 | $C24H26Cl2NO2^{+}$ | 7.22 | 2 | 5 | |
| CHEMBL259732 | C20H27N3O2 | 9.19 | 2 | 5 | |
| CHEMBL1257577 | C19H22N2O2S | 9.37 | 2 | 5 | |
| CHEMBL410832 | C23H25N3O2 | 9.39 | 2 | 5 | |
| CHEMBL1257578 | C20H24N2O2S | 9.59 | 2 | 5 | |
| CHEMBL4787638 | C21H18ClN3O2 | 2.08 | 2 | 6 | 5.55 |
| CHEMBL1823042 | C20H28FN3O2 | 5.00 | 2 | 6 | |
| CHEMBL4089699 | C19H19N3O2S | 5.42 | 2 | 6 | |
| CHEMBL1823047 | C22H30N4O2 | 5.54 | 2 | 6 | |
| CHEMBL4065208 | C23H22N4O2 | 5.56 | 2 | 6 | |

**Table S5** (cont.d)

| | | | | | |
|---|---|---|---|---|---|
| CHEMBL563926 | C21H26ClN3O2 | 6.20 | 2 | 6 | |
| CHEMBL3775282 | C26H28N4O2 | 7.16 | 2 | 6 | |
| CHEMBL1108 | C22H22FN3O2 | 7.49 | 2 | 6 | |
| CHEMBL4781551 | C24H21ClN4O2 | 4.08 | 2 | 7 | 5.42 |
| CHEMBL1957012 | C26H29N5O2 | 4.60 | 2 | 7 | |
| CHEMBL3973288 | C20H22FN3O2S | 4.84 | 2 | 7 | |
| CHEMBL523374 | C22H23N5O2 | 4.89 | 2 | 7 | |
| CHEMBL3218884 | C28H33N5O2 | 4.96 | 2 | 7 | |
| CHEMBL498042 | C24H23N5O2 | 5.08 | 2 | 7 | |
| CHEMBL497048 | C27H23N5O2 | 5.36 | 2 | 7 | |
| CHEMBL2336331 | C24H29N5O2 | 5.40 | 2 | 7 | |
| CHEMBL1939739 | C32H35FN4O2 | 5.44 | 2 | 7 | |
| CHEMBL1098847 | C24H21Cl3N2O2 | 5.58 | 2 | 7 | |
| CHEMBL3086038 | C30H33BrClN3O2 | 5.82 | 2 | 7 | |
| CHEMBL3806193 | C19H23Cl2N3O2 | 5.84 | 2 | 7 | |
| CHEMBL3775387 | C25H27N5O2 | 5.99 | 2 | 7 | |
| CHEMBL3806153 | C18H21ClFN3O2 | 6.01 | 2 | 7 | |
| CHEMBL501480 | C29H25N5O2 | 6.50 | 2 | 7 | |
| CHEMBL526466 | C25H23N5O2 | 7.10 | 2 | 7 | |
| CHEMBL399726 | C20H24N6O2 | 3.78 | 2 | 8 | 5.84 |
| CHEMBL2314064 | C24H32F3N3O2 | 4.41 | 2 | 8 | |
| CHEMBL3577935 | C23H34F3N3O2 | 4.50 | 2 | 8 | |
| CHEMBL2164565 | C19H18F3N3O2 | 4.63 | 2 | 8 | |
| CHEMBL1939742 | C30H37N5O2S | 4.75 | 2 | 8 | |
| CHEMBL2181489 | C23H20F3N3O2 | 5.52 | 2 | 8 | |
| CHEMBL3218650 | C27H30ClN5O2 | 5.60 | 2 | 8 | |
| CHEMBL3415593 | C19H26N6O2 | 5.77 | 2 | 8 | |
| CHEMBL1080489 | C22H27N5O2S | 5.80 | 2 | 8 | |
| CHEMBL478462 | C25H38N6O2 | 5.84 | 2 | 8 | |
| CHEMBL1956991 | C23H26ClFN4O2 | 5.89 | 2 | 8 | |
| CHEMBL3318989 | C23H24ClN5O2 | 6.00 | 2 | 8 | |

**Table S5** (cont.d)

| | | | | | |
|---|---|---|---|---|---|
| CHEMBL515025 | C24H36N6O2 | 6.15 | 2 | 8 | |
| CHEMBL478011 | C25H38N6O2 | 6.38 | 2 | 8 | |
| CHEMBL453894 | C25H26F3N3O2 | 6.46 | 2 | 8 | |
| CHEMBL2146813 | C38H48ClN5O2 | 6.80 | 2 | 8 | |
| CHEMBL568571 | C27H26F3N3O2 | 6.85 | 2 | 8 | |
| CHEMBL128007 | C28H28Cl2N4O2 | 7.00 | 2 | 8 | |
| CHEMBL525413 | C29H24ClN5O2 | 7.09 | 2 | 8 | |
| CHEMBL2324519 | C20H21FN6O2 | 3.67 | 2 | 9 | 5.14 |
| CHEMBL3290351 | C24H27FN6O2 | 4.05 | 2 | 9 | |
| CHEMBL484816 | C22H21N7O2 | 4.30 | 2 | 9 | |
| CHEMBL4094682 | C22H27ClN6O2 | 4.34 | 2 | 9 | |
| CHEMBL3422244 | C18H18F3N3O2S | 4.71 | 2 | 9 | |
| CHEMBL596700 | C20H23F3N4O2 | 4.88 | 2 | 9 | |
| CHEMBL1095476 | C28H25Cl3N4O2 | 5.06 | 2 | 9 | |
| CHEMBL589833 | C24H25Cl2FN4O2 | 5.10 | 2 | 9 | |
| CHEMBL502288 | C23H22N6O2S | 5.11 | 2 | 9 | |
| CHEMBL2170611 | C17H21F3N4O2 | 5.16 | 2 | 9 | |
| CHEMBL255645 | C28H31Cl2N5O2 | 5.36 | 2 | 9 | |
| CHEMBL4213486 | C19H15F2N5O2 | 5.70 | 2 | 9 | |
| CHEMBL429761 | C23H31N7O2 | 5.73 | 2 | 9 | |
| CHEMBL3318999 | C28H25N7O2 | 5.85 | 2 | 9 | |
| CHEMBL2177305 | C28H23ClFN5O2 | 5.89 | 2 | 9 | |
| CHEMBL446966 | C34H31N7O2 | 5.91 | 2 | 9 | |
| CHEMBL549635 | C29H33F4N3O2 | 6.20 | 2 | 9 | |
| CHEMBL3983509 | C20H20ClFN4O2S | 6.70 | 2 | 9 | |
| CHEMBL2151322 | C19H20FN7O2 | 4.28 | 2 | 10 | 5.38 |
| CHEMBL2324520 | C20H20F2N6O2 | 4.68 | 2 | 10 | |
| CHEMBL2315921 | C26H28F3N5O2 | 4.77 | 2 | 10 | |
| CHEMBL595944 | C27H26N8O2 | 4.98 | 2 | 10 | |
| CHEMBL3408394 | C37H24N6O2S2 | 5.24 | 2 | 10 | |
| CHEMBL551281 | C26H34F3N5O2 | 5.52 | 2 | 10 | |

**Table S5** (cont.d)

| | | | | | |
|---|---|---|---|---|---|
| CHEMBL553196 | C32H32F4N4O2 | 5.76 | 2 | 10 | |
| CHEMBL3323074 | C32H28FN7O2 | 5.85 | 2 | 10 | |
| CHEMBL3323073 | C32H28FN7O2 | 6.20 | 2 | 10 | |
| CHEMBL375330 | C34H38ClF3N4O2 | 6.77 | 2 | 10 | |
| CHEMBL3703025 | C22H25F3N6O2 | 4.50 | 2 | 11 | 4.94 |
| CHEMBL4209441 | C26H23FN8O2 | 4.62 | 2 | 11 | |
| CHEMBL4211893 | C30H29FN8O2 | 4.87 | 2 | 11 | |
| CHEMBL3318984 | C23H23F3N6O2 | 5.00 | 2 | 11 | |
| CHEMBL3353404 | C26H29ClN8O2 | 5.17 | 2 | 11 | |
| CHEMBL3746204 | C22H17F4N5O2 | 5.24 | 2 | 11 | |
| CHEMBL4640366 | C33H34N2O3 | 4.02 | 3 | 5 | 5.92 |
| CHEMBL3787069 | C26H26N2O3 | 4.35 | 3 | 5 | |
| CHEMBL498572 | C19H22N2O3 | 5.23 | 3 | 5 | |
| CHEMBL1671893 | C23H30N2O3 | 6.60 | 3 | 5 | |
| CHEMBL1642487 | C28H28N2O3 | 9.09 | 3 | 5 | |
| CHEMBL1642479 | C28H28N2O3 | 9.17 | 3 | 5 | |
| CHEMBL3741589 | C18H17Cl2NO3 | 4.11 | 3 | 6 | 5.92 |
| CHEMBL1000 | C21H25ClN2O3 | 4.52 | 3 | 6 | |
| CHEMBL3219616 | C24H30N2O3S | 5.80 | 3 | 6 | |
| CHEMBL1224699 | C21H26N2O3S | 5.90 | 3 | 6 | |
| CHEMBL561279 | C25H34N2O3S | 5.93 | 3 | 6 | |
| CHEMBL4173253 | C26H34N2O3S | 7.10 | 3 | 6 | |
| CHEMBL533 | C20H36N2O3S | 8.00 | 3 | 6 | |
| CHEMBL1258503 | C22H25N3O3 | 9.14 | 3 | 6 | |
| CHEMBL8 | C17H18FN3O3 | 3.02 | 3 | 7 | 4.99 |
| CHEMBL4855895 | C21H24FN3O3 | 3.53 | 3 | 7 | |
| CHEMBL22 | C14H18N4O3 | 3.62 | 3 | 7 | |
| CHEMBL388978 | C28H26N4O3 | 4.04 | 3 | 7 | |
| CHEMBL583 | C19H22FN3O3 | 4.30 | 3 | 7 | |
| CHEMBL2158823 | C24H28Cl2N2O3 | 4.50 | 3 | 7 | |
| CHEMBL3115194 | C26H32BrN3O3 | 4.90 | 3 | 7 | |

**Table S5** (cont.d)

| | | | | | |
|---|---|---|---|---|---|
| CHEMBL1957792 | C27H30ClN3O3 | 4.99 | 3 | 7 | |
| CHEMBL246815 | C22H26FN3O3 | 5.13 | 3 | 7 | |
| CHEMBL398478 | C25H28FN3O3 | 5.21 | 3 | 7 | |
| CHEMBL2158815 | C24H28Cl2N2O3 | 5.30 | 3 | 7 | |
| CHEMBL250533 | C25H28ClN3O3 | 5.48 | 3 | 7 | |
| CHEMBL3764774 | C28H28N4O3 | 5.70 | 3 | 7 | |
| CHEMBL45816 | C29H38FN3O3 | 6.24 | 3 | 7 | |
| CHEMBL2146854 | C23H26N4O3 | 7.33 | 3 | 7 | |
| CHEMBL540634 | C26H31Cl2N3O3 | 4.50 | 3 | 8 | 5.35 |
| CHEMBL2178221 | C28H29ClN4O3 | 4.82 | 3 | 8 | |
| CHEMBL3964723 | C23H27Cl2N3O3 | 5.00 | 3 | 8 | |
| CHEMBL465417 | C25H23N5O3 | 5.04 | 3 | 8 | |
| CHEMBL2158771 | C27H31Cl2N3O3 | 5.30 | 3 | 8 | |
| CHEMBL207220 | C24H40N4O3S | 5.40 | 3 | 8 | |
| CHEMBL2207738 | C23H26N4O3S | 5.60 | 3 | 8 | |
| CHEMBL3703237 | C19H17ClN4O3 | 5.66 | 3 | 8 | |
| CHEMBL1621 | C23H27FN4O3 | 6.00 | 3 | 8 | |
| CHEMBL2146853 | C20H23N5O3 | 7.60 | 3 | 8 | |
| CHEMBL3290344 | C24H26N6O3 | 4.13 | 3 | 9 | 5.07 |
| CHEMBL2165068 | C25H26FN5O3 | 4.24 | 3 | 9 | |
| CHEMBL3218891 | C27H32N6O3 | 4.37 | 3 | 9 | |
| CHEMBL1083118 | C17H19F2N3O3S | 4.89 | 3 | 9 | |
| CHEMBL2441417 | C24H26N6O3 | 4.89 | 3 | 9 | |
| CHEMBL255968 | C25H24ClN5O3 | 5.00 | 3 | 9 | |
| CHEMBL2164393 | C26H24F3N3O3 | 5.14 | 3 | 9 | |
| CHEMBL2164389 | C24H20F3N3O3 | 5.34 | 3 | 9 | |
| CHEMBL557306 | C21H24ClF3N2O3 | 5.99 | 3 | 9 | |
| CHEMBL514042 | C25H38N6O3 | 6.05 | 3 | 9 | |
| CHEMBL3422973 | C27H28N6O3 | 8.68 | 3 | 9 | |
| CHEMBL3422970 | C27H28N6O3 | 9.06 | 3 | 9 | |
| CHEMBL3221500 | C22H21ClN6O3 | 4.20 | 3 | 10 | 4.78 |

**Table S5** (cont.d)

| | | | | | |
|---|---|---|---|---|---|
| CHEMBL3794265 | C23H25FN6O3 | 4.38 | 3 | 10 | |
| CHEMBL3221488 | C23H23ClN6O3 | 4.50 | 3 | 10 | |
| CHEMBL3425929 | C28H33FN6O3 | 4.51 | 3 | 10 | |
| CHEMBL2164365 | C23H19F3N4O3 | 5.04 | 3 | 10 | |
| CHEMBL2164047 | C24H19F4N3O3 | 5.28 | 3 | 10 | |
| CHEMBL2403108 | C28H36ClN5O3S | 5.90 | 3 | 10 | |
| CHEMBL537847 | C23H29Cl2N5O3 | 6.25 | 3 | 10 | |
| CHEMBL1782574 | C23H35F3N4O3S | 2.41 | 3 | 11 | 5.00 |
| CHEMBL3221490 | C23H22ClN7O3 | 4.40 | 3 | 11 | |
| CHEMBL2204260 | C22H25F3N4O3S | 4.62 | 3 | 11 | |
| CHEMBL3407784 | C20H19ClN6O3S | 4.89 | 3 | 11 | |
| CHEMBL3221503 | C23H20ClN7O3 | 4.90 | 3 | 11 | |
| CHEMBL4076061 | C22H20ClF2N3O3S2 | 4.99 | 3 | 11 | |
| CHEMBL2204270 | C24H26F4N4O3 | 5.00 | 3 | 11 | |
| CHEMBL2331648 | C20H22F3N5O3 | 5.34 | 3 | 11 | |
| CHEMBL2147223 | C22H20ClF3N4O3 | 5.40 | 3 | 11 | |
| CHEMBL4075908 | C20H21BrFN5O3S | 5.46 | 3 | 11 | |
| CHEMBL2041175 | C31H27N7O3S | 5.82 | 3 | 11 | |
| CHEMBL1097456 | C26H22Cl3F3N2O3 | 5.84 | 3 | 11 | |
| CHEMBL3422978 | C26H26F3N5O3 | 9.41 | 3 | 11 | |
| CHEMBL1784523 | C36H40F3N5O3S | 4.00 | 3 | 12 | 4.86 |
| CHEMBL2206791 | C23H21F3N6O3 | 4.58 | 3 | 12 | |
| CHEMBL2147316 | C22H19F5N4O3 | 4.60 | 3 | 12 | |
| CHEMBL2147303 | C22H19F5N4O3 | 4.70 | 3 | 12 | |
| CHEMBL3221487 | C20H19ClN8O3 | 4.80 | 3 | 12 | |
| CHEMBL2064657 | C28H23ClF2N6O3 | 4.91 | 3 | 12 | |
| CHEMBL3764895 | C29H31Cl2F3N4O3 | 5.13 | 3 | 12 | |
| CHEMBL3764357 | C29H31Cl2F3N4O3 | 5.20 | 3 | 12 | |
| CHEMBL2441412 | C24H23F3N6O3 | 5.29 | 3 | 12 | |
| CHEMBL3765012 | C30H31Cl2F3N4O3 | 5.51 | 3 | 12 | |
| CHEMBL270190 | C25H32N2O4 | 3.10 | 4 | 6 | 5.88 |

**Table S5** (cont.d)

| | | | | | |
|---|---|---|---|---|---|
| CHEMBL519266 | C22H28N2O4 | 3.11 | 4 | 6 | |
| CHEMBL1938433 | C23H24N2O4 | 5.42 | 4 | 6 | |
| CHEMBL556100 | C22H25NO4S | 6.33 | 4 | 6 | |
| CHEMBL723 | C24H26N2O4 | 6.46 | 4 | 6 | |
| CHEMBL1258280 | C22H24N2O4 | 9.21 | 4 | 6 | |
| CHEMBL4847738 | C20H23N3O4 | 4.05 | 4 | 7 | 5.96 |
| CHEMBL3703271 | C22H23N3O4 | 5.08 | 4 | 7 | |
| CHEMBL246634 | C21H23FN2O4 | 5.58 | 4 | 7 | |
| CHEMBL17423 | C22H25N3O4 | 5.96 | 4 | 7 | |
| CHEMBL2387265 | C31H29FN2O4 | 6.09 | 4 | 7 | |
| CHEMBL384487 | C24H23FN2O4 | 7.69 | 4 | 7 | |
| CHEMBL217593 | C23H23FN2O4 | 8.27 | 4 | 7 | |
| CHEMBL33 | C18H20FN3O4 | 3.04 | 4 | 8 | 4.51 |
| CHEMBL31 | C19H22FN3O4 | 3.89 | 4 | 8 | |
| CHEMBL520463 | C28H24N4O4 | 4.05 | 4 | 8 | |
| CHEMBL32 | C21H24FN3O4 | 4.10 | 4 | 8 | |
| CHEMBL2382343 | C29H32F2N2O4 | 4.32 | 4 | 8 | |
| CHEMBL3787345 | C25H28N4O4 | 4.51 | 4 | 8 | |
| CHEMBL602875 | C26H34N4O4 | 4.72 | 4 | 8 | |
| CHEMBL3612814 | C24H28N4O4 | 4.85 | 4 | 8 | |
| CHEMBL1084617 | C24H28N4O4 | 4.96 | 4 | 8 | |
| CHEMBL3786346 | C25H26N4O4 | 5.05 | 4 | 8 | |
| CHEMBL2387229 | C26H26ClFN2O4 | 5.80 | 4 | 8 | |
| CHEMBL4092041 | C29H31F3N2O4 | 4.07 | 4 | 9 | 5.25 |
| CHEMBL248296 | C27H31F2N3O4 | 4.19 | 4 | 9 | |
| CHEMBL2158050 | C25H29FN2O4S2 | 4.63 | 4 | 9 | |
| CHEMBL3650850 | C21H23N5O4 | 4.68 | 4 | 9 | |
| CHEMBL4243499 | C19H16F3N5O | 5.25 | 4 | 9 | |
| CHEMBL4170966 | C24H21Cl2N3O4 | 5.33 | 4 | 9 | |
| CHEMBL1927161 | C24H29N5O4 | 5.40 | 4 | 9 | |
| CHEMBL244083 | C29H36N4O4S | 5.60 | 4 | 9 | |

**Table S5** (cont.d)

| | | | | | |
|---|---|---|---|---|---|
| CHEMBL1729 | C23H29ClFN3O4 | 8.19 | 4 | 9 | |
| CHEMBL2440387 | C25H23ClN4O4S | 3.38 | 4 | 10 | 4.94 |
| CHEMBL2424928 | C24H26FN5O4 | 3.51 | 4 | 10 | |
| CHEMBL1916543 | C23H28N6O4 | 3.69 | 4 | 10 | |
| CHEMBL2041188 | C32H30N6O4 | 4.64 | 4 | 10 | |
| CHEMBL3400819 | C26H26FN5O4 | 4.91 | 4 | 10 | |
| CHEMBL256060 | C26H26ClN5O4 | 4.96 | 4 | 10 | |
| CHEMBL513921 | C23H34N6O4 | 5.24 | 4 | 10 | |
| CHEMBL3604800 | C28H32FN5O4 | 5.52 | 4 | 10 | |
| CHEMBL3605145 | C29H32FN5O4 | 6.30 | 4 | 10 | |
| CHEMBL3605140 | C27H30FN5O4 | 6.40 | 4 | 10 | |
| CHEMBL3985847 | C21H26N6O4S | 2.40 | 4 | 11 | 4.60 |
| CHEMBL1916544 | C22H27N7O4 | 4.09 | 4 | 11 | |
| CHEMBL1091605 | C29H36F3N3O4S | 4.52 | 4 | 11 | |
| CHEMBL2069410 | C23H19F4N3O4 | 4.60 | 4 | 11 | |
| CHEMBL192 | C22H30N6O4S | 5.48 | 4 | 11 | |
| CHEMBL3422758 | C27H29F2N5O4 | 5.76 | 4 | 11 | |
| CHEMBL3422952 | C26H27F2N5O4 | 9.14 | 4 | 11 | |
| CHEMBL3799831 | C25H26FN3O5 | 4.03 | 5 | 9 | 5.00 |
| CHEMBL3612928 | C24H28N4O5 | 4.70 | 5 | 9 | |
| CHEMBL3786311 | C26H30N4O5 | 4.80 | 5 | 9 | |
| CHEMBL1091664 | C28H25ClN2O5S | 5.00 | 5 | 9 | |
| CHEMBL3612926 | C25H30N4O5 | 5.11 | 5 | 9 | |
| CHEMBL429458 | C25H30FN3O5 | 5.20 | 5 | 9 | |
| CHEMBL387178 | C23H20F2N2O5 | 8.21 | 5 | 9 | |
| CHEMBL3921669 | C23H26N4O5S | 2.22 | 5 | 10 | 4.99 |
| CHEMBL605785 | C27H27FN4O5 | 4.02 | 5 | 10 | |
| CHEMBL247690 | C29H33F2N3O5 | 4.47 | 5 | 10 | |
| CHEMBL245642 | C25H25F2N3O5 | 5.50 | 5 | 10 | |
| CHEMBL217707 | C23H19F3N2O5 | 8.25 | 5 | 10 | |
| CHEMBL473 | C19H27N3O5S2 | 8.39 | 5 | 10 | |

**Table S6.** Investigation of the influence of nitrogen atom on the $pIC_{50}$ value.

| Name | Molecular formula | pIC50 | N | Hetatoms | median value |
|---|---|---|---|---|---|
| CHEMBL3314420 | C15H13ClFN | 4.30 | 1 | 3 | 5.69 |
| CHEMBL1684047 | C16H23Cl2N | 4.40 | 1 | 3 | |
| CHEMBL2325209 | C26H28NO2$^{+}$ | 5.54 | 1 | 3 | |
| CHEMBL299390 | C19H19ClFN | 5.83 | 1 | 3 | |
| CHEMBL2324248 | C20H20NO2$^{+}$ | 6.12 | 1 | 3 | |
| CHEMBL607 | C15H21NO2 | 6.49 | 1 | 3 | |
| CHEMBL70 | C17H19NO3 | 3.00 | 1 | 4 | 4.89 |
| CHEMBL1671896 | C11H14F3N | 3.27 | 1 | 4 | |
| CHEMBL3220615 | C19H21NO3 | 4.89 | 1 | 4 | |
| CHEMBL53866 | C17H15ClFNO | 5.66 | 1 | 4 | |
| CHEMBL96153 | C24H31NO3 | 7.52 | 1 | 4 | |
| CHEMBL2324239 | C24H18Cl2NO2$^{+}$ | 4.51 | 1 | 5 | 6.10 |
| CHEMBL914 | C32H39NO4 | 4.67 | 1 | 5 | |
| CHEMBL378547 | C21H21F2NO2 | 5.11 | 1 | 5 | |
| CHEMBL196983 | C21H27NO4 | 6.10 | 1 | 5 | |
| CHEMBL2324243 | C26H32NO4$^{+}$ | 6.80 | 1 | 5 | |
| CHEMBL2324245 | C24H26Cl2NO2$^{+}$ | 7.22 | 1 | 5 | |
| CHEMBL572163 | C26H31NO4 | 7.42 | 1 | 5 | |
| CHEMBL284348 | C5H6N2 | 2.36 | 2 | 2 | 5.06 |
| CHEMBL3 | C10H14N2 | 3.61 | 2 | 2 | |
| CHEMBL1086033 | C17H22N2 | 3.67 | 2 | 2 | |
| CHEMBL87045 | C18H18N2 | 5.06 | 2 | 2 | |
| CHEMBL459176 | C20H22N2 | 5.16 | 2 | 2 | |
| CHEMBL11 | C19H24N2 | 5.47 | 2 | 2 | |
| CHEMBL1671894 | C27H26N2 | 7.75 | 2 | 2 | |
| CHEMBL1087 | C15H22N2O | 3.81 | 2 | 3 | 5.16 |
| CHEMBL1083707 | C20H24N2O | 4.67 | 2 | 3 | |
| CHEMBL513258 | C19H19FN2 | 5.05 | 2 | 3 | |
| CHEMBL1945694 | C16H15ClN2 | 5.16 | 2 | 3 | |

**Table S6** (Cont.d)

| | | | | | |
|---|---|---|---|---|---|
| CHEMBL191054 | C13H17ClN2 | 5.60 | 2 | 3 | |
| CHEMBL505 | C16H19ClN2 | 5.96 | 2 | 3 | |
| CHEMBL1090528 | C19H24N2S | 6.21 | 2 | 3 | |
| CHEMBL16 | C15H12N2O2 | 4.00 | 2 | 4 | 5.92 |
| CHEMBL3356251 | C24H36N2O2 | 4.01 | 2 | 4 | |
| CHEMBL3786542 | C26H26N2O2 | 4.21 | 2 | 4 | |
| CHEMBL3356249 | C25H38N2O2 | 4.30 | 2 | 4 | |
| CHEMBL3314426 | C15H14F2N2 | 4.66 | 2 | 4 | |
| CHEMBL1086273 | C19H23FN2O | 4.68 | 2 | 4 | |
| CHEMBL2333646 | C19H23BrN2O | 4.89 | 2 | 4 | |
| CHEMBL3185736 | C22H20N2O2 | 4.91 | 2 | 4 | |
| CHEMBL4162513 | C35H34N2O2 | 5.10 | 2 | 4 | |
| CHEMBL4450665 | C23H28N2O2 | 5.92 | 2 | 4 | |
| CHEMBL3323181 | C22H24N2OS | 6.00 | 2 | 4 | |
| CHEMBL1092377 | C19H21FN2S | 6.48 | 2 | 4 | |
| CHEMBL1346 | C28H30N2O2 | 7.10 | 2 | 4 | |
| CHEMBL2046892 | C22H22F2N2 | 7.40 | 2 | 4 | |
| CHEMBL1223951 | C21H24N2O2 | 9.21 | 2 | 4 | |
| CHEMBL1257938 | C22H26N2O2 | 9.21 | 2 | 4 | |
| CHEMBL1257937 | C22H26N2O2 | 9.28 | 2 | 4 | |
| CHEMBL1257820 | C23H28N2O2 | 9.60 | 2 | 4 | |
| CHEMBL1257821 | C22H26N2O2 | 9.85 | 2 | 4 | |
| CHEMBL4640366 | C33H34N2O3 | 4.02 | 2 | 5 | 6.17 |
| CHEMBL4209354 | C15H20N2O2S | 4.07 | 2 | 5 | |
| CHEMBL3787069 | C26H26N2O3 | 4.35 | 2 | 5 | |
| CHEMBL1080790 | C16H22Cl2N2O | 4.78 | 2 | 5 | |
| CHEMBL1080969 | C16H22Cl2N2O | 5.11 | 2 | 5 | |
| CHEMBL498572 | C19H22N2O3 | 5.23 | 2 | 5 | |
| CHEMBL4100591 | C27H34F2N2O | 5.29 | 2 | 5 | |
| CHEMBL1224697 | C22H28N2O2S | 5.90 | 2 | 5 | |

**Table S6** (Cont.d)

| | | | | | |
|---|---|---|---|---|---|
| CHEMBL376488 | C32H31BrN2O2 | 6.43 | 2 | 5 | |
| CHEMBL1671893 | C23H30N2O3 | 6.60 | 2 | 5 | |
| CHEMBL998 | C22H23ClN2O2 | 6.77 | 2 | 5 | |
| CHEMBL195180 | C24H27ClN2OS | 7.52 | 2 | 5 | |
| CHEMBL1642487 | C28H28N2O3 | 9.09 | 2 | 5 | |
| CHEMBL1642479 | C28H28N2O3 | 9.17 | 2 | 5 | |
| CHEMBL1257577 | C19H22N2O2S | 9.37 | 2 | 5 | |
| CHEMBL1257578 | C20H24N2O2S | 9.59 | 2 | 5 | |
| CHEMBL270190 | C25H32N2O4 | 3.10 | 2 | 6 | 5.90 |
| CHEMBL519266 | C22H28N2O4 | 3.11 | 2 | 6 | |
| CHEMBL1000 | C21H25ClN2O3 | 4.52 | 2 | 6 | |
| CHEMBL1938433 | C23H24N2O4 | 5.42 | 2 | 6 | |
| CHEMBL3219616 | C24H30N2O3S | 5.80 | 2 | 6 | |
| CHEMBL1224699 | C21H26N2O3S | 5.90 | 2 | 6 | |
| CHEMBL561279 | C25H34N2O3S | 5.93 | 2 | 6 | |
| CHEMBL723 | C24H26N2O4 | 6.46 | 2 | 6 | |
| CHEMBL4173253 | C26H34N2O3S | 7.10 | 2 | 6 | |
| CHEMBL533 | C20H36N2O3S | 8.00 | 2 | 6 | |
| CHEMBL1258280 | C22H24N2O4 | 9.21 | 2 | 6 | |
| CHEMBL374731 | C10H14N2O5 | 2.30 | 2 | 7 | 5.05 |
| CHEMBL3329814 | C21H28N2O5 | 4.09 | 2 | 7 | |
| CHEMBL2158823 | C24H28Cl2N2O3 | 4.50 | 2 | 7 | |
| CHEMBL2333636 | C24H28F4N2O | 4.75 | 2 | 7 | |
| CHEMBL2333616 | C20H22ClF3N2O | 4.92 | 2 | 7 | |
| CHEMBL2333607 | C25H30F4N2O | 5.05 | 2 | 7 | |
| CHEMBL2333615 | C23H26F4N2O | 5.05 | 2 | 7 | |
| CHEMBL2158815 | C24H28Cl2N2O3 | 5.30 | 2 | 7 | |
| CHEMBL246634 | C21H23FN2O4 | 5.58 | 2 | 7 | |
| CHEMBL1098847 | C24H21Cl3N2O2 | 5.58 | 2 | 7 | |
| CHEMBL2387265 | C31H29FN2O4 | 6.09 | 2 | 7 | |
| CHEMBL384487 | C24H23FN2O4 | 7.69 | 2 | 7 | |

**Table S6** (Cont.d)

| | | | | | |
|---|---|---|---|---|---|
| CHEMBL217593 | C23H23FN2O4 | 8.27 | 2 | 7 | |
| CHEMBL193 | C17H18N2O6 | 4.30 | 2 | 8 | 5.41 |
| CHEMBL2382343 | C29H32F2N2O4 | 4.32 | 2 | 8 | |
| CHEMBL2440407 | C23H26N2O5S | 5.02 | 2 | 8 | |
| CHEMBL2387229 | C26H26ClFN2O4 | 5.80 | 2 | 8 | |
| CHEMBL214021 | C23H21FN2O5 | 7.82 | 2 | 8 | |
| CHEMBL217789 | C23H21ClN2O5 | 8.44 | 2 | 8 | |
| CHEMBL216323 | C23H19FN2O6 | 4.07 | 2 | 9 | 5.00 |
| CHEMBL4092041 | C29H31F3N2O4 | 4.07 | 2 | 9 | |
| CHEMBL2158050 | C25H29FN2O4S2 | 4.63 | 2 | 9 | |
| CHEMBL1091664 | C28H25ClN2O5S | 5.00 | 2 | 9 | |
| CHEMBL557306 | C21H24ClF3N2O3 | 5.99 | 2 | 9 | |
| CHEMBL217442 | C24H23FN2O6 | 7.80 | 2 | 9 | |
| CHEMBL387178 | C23H20F2N2O5 | 8.21 | 2 | 9 | |
| CHEMBL428594 | C13H21N3O | 3.86 | 3 | 4 | 5.40 |
| CHEMBL217707 | C17H25N3O | 3.94 | 3 | 4 | |
| CHEMBL213715 | C17H33N3O | 4.17 | 3 | 4 | |
| CHEMBL1091218 | C20H31N3O | 4.42 | 3 | 4 | |
| CHEMBL1097456 | C22H23N3O | 4.70 | 3 | 4 | |
| CHEMBL3417745 | C21H29N3O | 4.92 | 3 | 4 | |
| CHEMBL402624 | C22H33N3O | 5.40 | 3 | 4 | |
| CHEMBL490026 | C21H22ClN3 | 5.44 | 3 | 4 | |
| CHEMBL497171 | C20H25N3O | 5.47 | 3 | 4 | |
| CHEMBL457930 | C20H23N3O | 5.68 | 3 | 4 | |
| CHEMBL299499 | C17H21N3S | 5.70 | 3 | 4 | |
| CHEMBL640 | C22H26ClN3 | 5.96 | 3 | 4 | |
| CHEMBL219803 | C23H25N3O | 6.13 | 3 | 4 | |
| CHEMBL219128 | C15H23N3O2 | 4.00 | 3 | 5 | 5.18 |
| CHEMBL219074 | C19H25N3O2 | 4.12 | 3 | 5 | |
| CHEMBL1089159 | C23H27N3O2 | 4.14 | 3 | 5 | |
| CHEMBL517 | C22H23N3O2 | 4.27 | 3 | 5 | |

**Table S6** (Cont.d)

| | | | | | |
|---|---|---|---|---|---|
| CHEMBL1094041 | C29H29N3O2 | 4.55 | 3 | 5 | |
| CHEMBL2441636 | C22H33N3O2 | 4.72 | 3 | 5 | |
| CHEMBL2177304 | C27H24FN3O | 5.07 | 3 | 5 | |
| CHEMBL487063 | C23H27N3O2 | 5.10 | 3 | 5 | |
| CHEMBL485242 | C25H29N3O2 | 5.14 | 3 | 5 | |
| CHEMBL478615 | C23H27N3O2 | 5.17 | 3 | 5 | |
| CHEMBL2326478 | C18H18FN3S | 5.17 | 3 | 5 | |
| CHEMBL476579 | C24H29N3O2 | 5.18 | 3 | 5 | |
| CHEMBL479242 | C23H27N3O2 | 5.18 | 3 | 5 | |
| CHEMBL3121096 | C20H25N3O2 | 5.22 | 3 | 5 | |
| CHEMBL1819137 | C22H25N3O2 | 5.23 | 3 | 5 | |
| CHEMBL488249 | C24H27N3O2 | 5.25 | 3 | 5 | |
| CHEMBL384400 | C27H36ClN3O | 5.48 | 3 | 5 | |
| CHEMBL487064 | C23H27N3O2 | 5.55 | 3 | 5 | |
| CHEMBL478616 | C23H27N3O2 | 5.80 | 3 | 5 | |
| CHEMBL562833 | C22H30ClN3O | 6.10 | 3 | 5 | |
| CHEMBL259732 | C20H27N3O2 | 9.19 | 3 | 5 | |
| CHEMBL410832 | C23H25N3O2 | 9.39 | 3 | 5 | |
| CHEMBL4787638 | C21H18ClN3O2 | 2.08 | 3 | 6 | 6.20 |
| CHEMBL1823042 | C20H28FN3O2 | 5.00 | 3 | 6 | |
| CHEMBL4089699 | C19H19N3O2S | 5.42 | 3 | 6 | |
| CHEMBL563926 | C21H26ClN3O2 | 6.20 | 3 | 6 | |
| CHEMBL1108 | C22H22FN3O2 | 7.49 | 3 | 6 | |
| CHEMBL1423 | C28H29F2N3O | 8.52 | 3 | 6 | |
| CHEMBL1258503 | C22H25N3O3 | 9.14 | 3 | 6 | |
| CHEMBL8 | C17H18FN3O3 | 3.02 | 3 | 7 | 5.04 |
| CHEMBL4855895 | C21H24FN3O3 | 3.53 | 3 | 7 | |
| CHEMBL4847738 | C20H23N3O4 | 4.05 | 3 | 7 | |
| CHEMBL583 | C19H22FN3O3 | 4.30 | 3 | 7 | |
| CHEMBL218836 | C18H24F3N3O | 4.40 | 3 | 7 | |
| CHEMBL2314058 | C25H34F3N3O | 4.83 | 3 | 7 | |

**Table S6** (Cont.d)

| | | | | | |
|---|---|---|---|---|---|
| CHEMBL3973288 | C20H22FN3O2S | 4.84 | 3 | 7 | |
| CHEMBL3115194 | C26H32BrN3O3 | 4.90 | 3 | 7 | |
| CHEMBL1957792 | C27H30ClN3O3 | 4.99 | 3 | 7 | |
| CHEMBL3703271 | C22H23N3O4 | 5.08 | 3 | 7 | |
| CHEMBL246815 | C22H26FN3O3 | 5.13 | 3 | 7 | |
| CHEMBL398478 | C25H28FN3O3 | 5.21 | 3 | 7 | |
| CHEMBL250533 | C25H28ClN3O3 | 5.48 | 3 | 7 | |
| CHEMBL3086038 | C30H33BrClN3O2 | 5.82 | 3 | 7 | |
| CHEMBL3806193 | C19H23Cl2N3O2 | 5.84 | 3 | 7 | |
| CHEMBL17423 | C22H25N3O4 | 5.96 | 3 | 7 | |
| CHEMBL3806153 | C18H21ClFN3O2 | 6.01 | 3 | 7 | |
| CHEMBL45816 | C29H38FN3O3 | 6.24 | 3 | 7 | |
| CHEMBL33 | C18H20FN3O4 | 3.04 | 3 | 8 | 4.57 |
| CHEMBL31 | C19H22FN3O4 | 3.89 | 3 | 8 | |
| CHEMBL32 | C21H24FN3O4 | 4.10 | 3 | 8 | |
| CHEMBL2333619 | C22H27F4N3O | 4.10 | 3 | 8 | |
| CHEMBL2314064 | C24H32F3N3O2 | 4.41 | 3 | 8 | |
| CHEMBL540634 | C26H31Cl2N3O3 | 4.50 | 3 | 8 | |
| CHEMBL3577935 | C23H34F3N3O2 | 4.50 | 3 | 8 | |
| CHEMBL2164565 | C19H18F3N3O2 | 4.63 | 3 | 8 | |
| CHEMBL3964723 | C23H27Cl2N3O3 | 5.00 | 3 | 8 | |
| CHEMBL2158771 | C27H31Cl2N3O3 | 5.30 | 3 | 8 | |
| CHEMBL2181489 | C23H20F3N3O2 | 5.52 | 3 | 8 | |
| CHEMBL2441431 | C22H23N3O5 | 5.89 | 3 | 8 | |
| CHEMBL453894 | C25H26F3N3O2 | 6.46 | 3 | 8 | |
| CHEMBL568571 | C27H26F3N3O2 | 6.85 | 3 | 8 | |
| CHEMBL3799831 | C25H26FN3O5 | 4.03 | 3 | 9 | 5.14 |
| CHEMBL248296 | C27H31F2N3O4 | 4.19 | 3 | 9 | |
| CHEMBL4084170 | C15H18F5N3S | 4.69 | 3 | 9 | |
| CHEMBL3422244 | C18H18F3N3O2S | 4.71 | 3 | 9 | |
| CHEMBL1083118 | C17H19F2N3O3S | 4.89 | 3 | 9 | |

**Table S6** (Cont.d)

| | | | | | |
|---|---|---|---|---|---|
| CHEMBL2164393 | C26H24F3N3O3 | 5.14 | 3 | 9 | |
| CHEMBL429458 | C25H30FN3O5 | 5.20 | 3 | 9 | |
| CHEMBL4170966 | C24H21Cl2N3O4 | 5.33 | 3 | 9 | |
| CHEMBL2164389 | C24H20F3N3O3 | 5.34 | 3 | 9 | |
| CHEMBL549635 | C29H33F4N3O2 | 6.20 | 3 | 9 | |
| CHEMBL1729 | C23H29ClFN3O4 | 8.19 | 3 | 9 | |
| CHEMBL247690 | C29H33F2N3O5 | 4.47 | 3 | 10 | 5.50 |
| CHEMBL2164047 | C24H19F4N3O3 | 5.28 | 3 | 10 | |
| CHEMBL245642 | C25H25F2N3O5 | 5.50 | 3 | 10 | |
| CHEMBL4081080 | C25H17F6N3O | 7.17 | 3 | 10 | |
| CHEMBL473 | C19H27N3O5S2 | 8.39 | 3 | 10 | |
| CHEMBL1091605 | C25H34N4O | 4.62 | 4 | 5 | 5.49 |
| CHEMBL2069410 | C18H22N4O | 4.68 | 4 | 5 | |
| CHEMBL2164375 | C20H24N4O | 4.71 | 4 | 5 | |
| CHEMBL4076061 | C16H18N4S | 4.90 | 4 | 5 | |
| CHEMBL2207664 | C20H22N4O | 4.94 | 4 | 5 | |
| CHEMBL435882 | C15H14N4O | 5.01 | 4 | 5 | |
| CHEMBL4213196 | C21H28N4S | 5.33 | 4 | 5 | |
| CHEMBL217133 | C17H18N4O | 5.64 | 4 | 5 | |
| CHEMBL429115 | C31H38N4O | 6.24 | 4 | 5 | |
| CHEMBL1125 | C26H28N4O | 6.28 | 4 | 5 | |
| CHEMBL4096683 | C17H26N4O | 6.63 | 4 | 5 | |
| CHEMBL565829 | C18H19ClN4 | 6.72 | 4 | 5 | |
| CHEMBL1956112 | C17H20N4S | 6.74 | 4 | 5 | |
| CHEMBL1110 | C19H21FN4 | 7.56 | 4 | 5 | |
| CHEMBL58387 | C18H19ClN4O | 3.85 | 4 | 6 | 5.61 |
| CHEMBL3775729 | C26H35FN4O | 4.88 | 4 | 6 | |
| CHEMBL1087493 | C21H34N4S2 | 4.88 | 4 | 6 | |
| CHEMBL42 | C20H21FN4O | 5.11 | 4 | 6 | |
| CHEMBL715 | C20H22N4OS | 5.23 | 4 | 6 | |
| CHEMBL61301 | C24H26N4OS | 5.29 | 4 | 6 | |

**Table S6** (Cont.d)

| | | | | | |
|---|---|---|---|---|---|
| CHEMBL1823047 | C22H30N4O2 | 5.54 | 4 | 6 | |
| CHEMBL4065208 | C23H22N4O2 | 5.56 | 4 | 6 | |
| CHEMBL2146805 | C20H21ClN4O | 5.60 | 4 | 6 | |
| CHEMBL4104102 | C20H22N4OS | 5.61 | 4 | 6 | |
| CHEMBL2177905 | C24H16F2N4 | 5.80 | 4 | 6 | |
| CHEMBL1083463 | C26H26Cl2N4 | 6.22 | 4 | 6 | |
| CHEMBL3774817 | C25H25FN4O | 6.59 | 4 | 6 | |
| CHEMBL522456 | CHEMBL522456 | 7.00 | 4 | 6 | |
| CHEMBL3775282 | C26H28N4O2 | 7.16 | 4 | 6 | |
| CHEMBL12780 | C24H27FN4O | 8.16 | 4 | 6 | |
| CHEMBL2146810 | C23H27ClN4O | 8.40 | 4 | 6 | |
| CHEMBL296419 | C28H31FN4O | 9.05 | 4 | 6 | |
| CHEMBL22 | C14H18N4O3 | 3.62 | 4 | 7 | 5.44 |
| CHEMBL3582294 | C25H18F2N4O | 4.03 | 4 | 7 | |
| CHEMBL388978 | C28H26N4O3 | 4.04 | 4 | 7 | |
| CHEMBL4781551 | C24H21ClN4O2 | 4.08 | 4 | 7 | |
| CHEMBL1092106 | C20H24ClFN4O | 5.04 | 4 | 7 | |
| CHEMBL1091834 | C25H26ClFN4O | 5.42 | 4 | 7 | |
| CHEMBL1939739 | C32H35FN4O2 | 5.44 | 4 | 7 | |
| CHEMBL3764774 | C28H28N4O3 | 5.70 | 4 | 7 | |
| CHEMBL3775050 | C25H24F2N4O | 6.20 | 4 | 7 | |
| CHEMBL1084400 | C20H21F3N4 | 6.80 | 4 | 7 | |
| CHEMBL708 | C21H21ClN4OS | 6.92 | 4 | 7 | |
| CHEMBL2146854 | C23H26N4O3 | 7.33 | 4 | 7 | |
| CHEMBL12713 | C24H26ClFN4O | 8.57 | 4 | 7 | |
| CHEMBL520463 | C28H24N4O4 | 4.05 | 4 | 8 | 5.01 |
| CHEMBL485123 | C20H23F3N4O | 4.07 | 4 | 8 | |
| CHEMBL3787345 | C25H28N4O4 | 4.51 | 4 | 8 | |
| CHEMBL602875 | C26H34N4O4 | 4.72 | 4 | 8 | |
| CHEMBL2178221 | C28H29ClN4O3 | 4.82 | 4 | 8 | |
| CHEMBL3612814 | C24H28N4O4 | 4.85 | 4 | 8 | |

**Table S6** (Cont.d)

| | | | | | |
|---|---|---|---|---|---|
| CHEMBL1084617 | C24H28N4O4 | 4.96 | 4 | 8 | |
| CHEMBL3786346 | C25H26N4O4 | 5.05 | 4 | 8 | |
| CHEMBL207220 | C24H40N4O3S | 5.40 | 4 | 8 | |
| CHEMBL2207738 | C23H26N4O3S | 5.60 | 4 | 8 | |
| CHEMBL3703237 | C19H17ClN4O3 | 5.66 | 4 | 8 | |
| CHEMBL1956991 | C23H26ClFN4O2 | 5.89 | 4 | 8 | |
| CHEMBL1621 | C23H27FN4O3 | 6.00 | 4 | 8 | |
| CHEMBL128007 | C28H28Cl2N4O2 | 7.00 | 4 | 8 | |
| CHEMBL3612928 | C24H28N4O5 | 4.70 | 4 | 9 | 5.11 |
| CHEMBL3786311 | C26H30N4O5 | 4.80 | 4 | 9 | |
| CHEMBL596700 | C20H23F3N4O2 | 4.88 | 4 | 9 | |
| CHEMBL1095476 | C28H25Cl3N4O2 | 5.06 | 4 | 9 | |
| CHEMBL589833 | C24H25Cl2FN4O2 | 5.10 | 4 | 9 | |
| CHEMBL3612926 | C25H30N4O5 | 5.11 | 4 | 9 | |
| CHEMBL2170611 | C17H21F3N4O2 | 5.16 | 4 | 9 | |
| CHEMBL4243499 | C19H16F3N5O | 5.25 | 4 | 9 | |
| CHEMBL244083 | C29H36N4O4S | 5.60 | 4 | 9 | |
| CHEMBL550410 | C25H26F4N4O | 5.92 | 4 | 9 | |
| CHEMBL3983509 | C20H20ClFN4O2S | 6.70 | 4 | 9 | |
| CHEMBL3921669 | C23H26N4O5S | 2.22 | 4 | 10 | 5.04 |
| CHEMBL2440387 | C25H23ClN4O4S | 3.38 | 4 | 10 | |
| CHEMBL605785 | C27H27FN4O5 | 4.02 | 4 | 10 | |
| CHEMBL2164365 | C23H19F3N4O3 | 5.04 | 4 | 10 | |
| CHEMBL560063 | C24H23Cl2F3N4O | 5.46 | 4 | 10 | |
| CHEMBL553196 | C32H32F4N4O2 | 5.76 | 4 | 10 | |
| CHEMBL375330 | C34H38ClF3N4O2 | 6.77 | 4 | 10 | |
| CHEMBL1782574 | C23H35F3N4O3S | 2.41 | 4 | 11 | 4.81 |
| CHEMBL2204260 | C22H25F3N4O3S | 4.62 | 4 | 11 | |
| CHEMBL2204270 | C24H26F4N4O3 | 5.00 | 4 | 11 | |
| CHEMBL2147223 | C22H20ClF3N4O3 | 5.40 | 4 | 11 | |
| CHEMBL256154 | C27H32F6N4O | 5.85 | 4 | 11 | |

**Table S6** (Cont.d)

| | | | | | |
|---|---|---|---|---|---|
| CHEMBL491571 | C15H24N4O6S2 | 2.10 | 4 | 12 | 4.92 |
| CHEMBL2147316 | C22H19F5N4O3 | 4.60 | 4 | 12 | |
| CHEMBL2147303 | C22H19F5N4O3 | 4.70 | 4 | 12 | |
| CHEMBL3764895 | C29H31Cl2F3N4O3 | 5.13 | 4 | 12 | |
| CHEMBL3764357 | C29H31Cl2F3N4O3 | 5.20 | 4 | 12 | |
| CHEMBL3765012 | C30H31Cl2F3N4O3 | 5.51 | 4 | 12 | |
| CHEMBL370072 | C24H31N5O | 4.64 | 5 | 6 | 5.91 |
| CHEMBL401608 | C21H24ClN5 | 5.05 | 5 | 6 | |
| CHEMBL487708 | C23H16FN5 | 5.51 | 5 | 6 | |
| CHEMBL237191 | C25H19N5O | 6.30 | 5 | 6 | |
| CHEMBL4212656 | C22H27N5O | 6.52 | 5 | 6 | |
| CHEMBL494406 | C32H33N5O | 7.62 | 5 | 6 | |
| CHEMBL2177904 | C21H25N5OS | 3.72 | 5 | 7 | 5.26 |
| CHEMBL4081790 | C22H22ClN5O | 4.22 | 5 | 7 | |
| CHEMBL4096145 | C21H21N5OS | 4.52 | 5 | 7 | |
| CHEMBL329067 | C26H29N5O2 | 4.60 | 5 | 7 | |
| CHEMBL578834 | C21H23ClFN5 | 4.64 | 5 | 7 | |
| CHEMBL4127458 | C17H22FN5O | 4.80 | 5 | 7 | |
| CHEMBL2069925 | C22H23N5O2 | 4.89 | 5 | 7 | |
| CHEMBL1957012 | C28H33N5O2 | 4.96 | 5 | 7 | |
| CHEMBL512975 | C24H23N5O2 | 5.08 | 5 | 7 | |
| CHEMBL2180842 | C21H15F2N5 | 5.16 | 5 | 7 | |
| CHEMBL523374 | C27H23N5O2 | 5.36 | 5 | 7 | |
| CHEMBL3218884 | C24H29N5O2 | 5.40 | 5 | 7 | |
| CHEMBL498042 | C25H22FN5O | 5.70 | 5 | 7 | |
| CHEMBL523821 | C23H31N5OS | 5.90 | 5 | 7 | |
| CHEMBL497048 | C19H20ClN5O | 5.90 | 5 | 7 | |
| CHEMBL2336331 | C25H27N5O2 | 5.99 | 5 | 7 | |
| CHEMBL2151218 | C28H31N5OS | 6.20 | 5 | 7 | |
| CHEMBL243901 | C29H25N5O2 | 6.50 | 5 | 7 | |
| CHEMBL2146812 | C23H24FN5S | 7.00 | 5 | 7 | |

**Table S6** (Cont.d)

| | | | | | |
|---|---|---|---|---|---|
| CHEMBL526466 | C25H23N5O2 | 7.10 | 5 | 7 | |
| CHEMBL3804950 | C20H21F2N5O | 4.01 | 5 | 8 | 5.70 |
| CHEMBL3216126 | C25H35Cl2N5O | 4.65 | 5 | 8 | |
| CHEMBL1939742 | C30H37N5O2S | 4.75 | 5 | 8 | |
| CHEMBL1093696 | C24H25ClFN5O | 4.92 | 5 | 8 | |
| CHEMBL465417 | C25H23N5O3 | 5.04 | 5 | 8 | |
| CHEMBL506163 | C22H19ClFN5O | 5.20 | 5 | 8 | |
| CHEMBL3218650 | C27H30ClN5O2 | 5.60 | 5 | 8 | |
| CHEMBL1080489 | C22H27N5O2S | 5.80 | 5 | 8 | |
| CHEMBL3318989 | C23H24ClN5O2 | 6.00 | 5 | 8 | |
| CHEMBL244280 | C26H28FN5OS | 6.60 | 5 | 8 | |
| CHEMBL2146813 | C38H48ClN5O2 | 6.80 | 5 | 8 | |
| CHEMBL525413 | C29H24ClN5O2 | 7.09 | 5 | 8 | |
| CHEMBL244278 | C26H28ClN5OS | 7.10 | 5 | 8 | |
| CHEMBL2146853 | C20H23N5O3 | 7.60 | 5 | 8 | |
| CHEMBL2165068 | C25H26FN5O3 | 4.24 | 5 | 9 | 5.42 |
| CHEMBL3650850 | C21H23N5O4 | 4.68 | 5 | 9 | |
| CHEMBL255968 | C25H24ClN5O3 | 5.00 | 5 | 9 | |
| CHEMBL4075071 | C19H16F3N5O | 5.25 | 5 | 9 | |
| CHEMBL255645 | C28H31Cl2N5O2 | 5.36 | 5 | 9 | |
| CHEMBL1927161 | C24H29N5O4 | 5.40 | 5 | 9 | |
| CHEMBL2158625 | C20H19Cl2N5OS | 5.44 | 5 | 9 | |
| CHEMBL489324 | C28H42ClF2N5O | 5.68 | 5 | 9 | |
| CHEMBL4213486 | C19H15F2N5O2 | 5.70 | 5 | 9 | |
| CHEMBL2158629 | C20H21Cl2N5OS | 5.85 | 5 | 9 | |
| CHEMBL2177305 | C28H23ClFN5O2 | 5.89 | 5 | 9 | |
| CHEMBL1081747 | C21H23ClFN5OS | 6.40 | 5 | 9 | |
| CHEMBL2424928 | C24H26FN5O4 | 3.51 | 5 | 10 | 5.71 |
| CHEMBL2315921 | C26H28F3N5O2 | 4.77 | 5 | 10 | |
| CHEMBL3400819 | C26H26FN5O4 | 4.91 | 5 | 10 | |

**Table S6** (Cont.d)

| | | | | | |
|---|---|---|---|---|---|
| CHEMBL256060 | C26H26ClN5O4 | 4.96 | 5 | 10 | |
| CHEMBL551281 | C26H34F3N5O2 | 5.52 | 5 | 10 | |
| CHEMBL3604800 | C28H32FN5O4 | 5.52 | 5 | 10 | |
| CHEMBL2403108 | C28H36ClN5O3S | 5.90 | 5 | 10 | |
| CHEMBL537847 | C23H29Cl2N5O3 | 6.25 | 5 | 10 | |
| CHEMBL3605145 | C29H32FN5O4 | 6.30 | 5 | 10 | |
| CHEMBL3605140 | C27H30FN5O4 | 6.40 | 5 | 10 | |
| CHEMBL1081746 | C23H26F3N5S2 | 6.46 | 5 | 10 | |
| CHEMBL401576 | C25H26F3N5OS | 6.90 | 5 | 10 | |
| CHEMBL3400817 | C26H26FN5O5 | 3.76 | 5 | 11 | 5.76 |
| CHEMBL3746204 | C22H17F4N5O2 | 5.24 | 5 | 11 | |
| CHEMBL2331648 | C20H22F3N5O3 | 5.34 | 5 | 11 | |
| CHEMBL4075908 | C20H21BrFN5O3S | 5.46 | 5 | 11 | |
| CHEMBL3422758 | C27H29F2N5O4 | 5.76 | 5 | 11 | |
| CHEMBL1082111 | C22H23F4N5OS | 5.90 | 5 | 11 | |
| CHEMBL3605123 | C30H36FN5O5 | 6.16 | 5 | 11 | |
| CHEMBL3422952 | C26H27F2N5O4 | 9.14 | 5 | 11 | |
| CHEMBL3422978 | C26H26F3N5O3 | 9.41 | 5 | 11 | |
| CHEMBL3425799 | C26H25F2N5O5 | 3.48 | 5 | 12 | 4.08 |
| CHEMBL1784523 | C36H40F3N5O3S | 4.00 | 5 | 12 | |
| CHEMBL451887 | C40H57N5O7 | 4.04 | 5 | 12 | |
| CHEMBL3342693 | C23H22F5N5O2 | 4.08 | 5 | 12 | |
| CHEMBL3605131 | C29H32FN5O6 | 4.91 | 5 | 12 | |
| CHEMBL550535 | C27H33F6N5O | 5.41 | 5 | 12 | |
| CHEMBL3605006 | C31H34FN5O6 | 5.80 | 5 | 12 | |
| CHEMBL3422957 | C14H16N6O | 4.03 | 6 | 7 | 5.28 |
| CHEMBL231396 | C21H26N6S | 4.71 | 6 | 7 | |
| CHEMBL2147024 | C23H26N6O | 5.09 | 6 | 7 | |
| CHEMBL4224807 | C23H19FN6 | 5.28 | 6 | 7 | |
| CHEMBL575241 | C20H20N6O | 5.42 | 6 | 7 | |

**Table S6** (Cont.d)

| | | | | | |
|---|---|---|---|---|---|
| CHEMBL2147021 | C20H15FN6 | 5.57 | 6 | 7 | |
| CHEMBL4279819 | C16H18N6O | 8.07 | 6 | 7 | |
| CHEMBL399726 | C20H24N6O2 | 3.78 | 6 | 8 | 5.60 |
| CHEMBL4858551 | C19H19FN6O | 4.01 | 6 | 8 | |
| CHEMBL256653 | C23H26N6OS | 4.30 | 6 | 8 | |
| CHEMBL257901 | C28H30N6OS | 4.90 | 6 | 8 | |
| CHEMBL270239 | C30H34N6OS | 4.90 | 6 | 8 | |
| CHEMBL515001 | C21H25FN6O | 4.96 | 6 | 8 | |
| CHEMBL2325997 | C20H23ClN6O | 5.14 | 6 | 8 | |
| CHEMBL3287218 | C29H32N6OS | 5.16 | 6 | 8 | |
| CHEMBL270852 | C32H38N6OS | 5.40 | 6 | 8 | |
| CHEMBL271909 | C28H30N6S2 | 5.50 | 6 | 8 | |
| CHEMBL272086 | C32H38N6OS | 5.60 | 6 | 8 | |
| CHEMBL2178521 | C24H21FN6O | 5.75 | 6 | 8 | |
| CHEMBL3415593 | C19H26N6O2 | 5.77 | 6 | 8 | |
| CHEMBL478462 | C25H38N6O2 | 5.84 | 6 | 8 | |
| CHEMBL515025 | C24H36N6O2 | 6.15 | 6 | 8 | |
| CHEMBL478011 | C25H38N6O2 | 6.38 | 6 | 8 | |
| CHEMBL408169 | C28H32N6OS | 6.40 | 6 | 8 | |
| CHEMBL94454 | C24H25FN6O | 6.46 | 6 | 8 | |
| CHEMBL233047 | C20H32Cl2N6 | 6.52 | 6 | 8 | |
| CHEMBL256442 | C29H32N6S2 | 6.80 | 6 | 8 | |
| CHEMBL397429 | C28H30N6OS | 7.30 | 6 | 8 | |
| CHEMBL2324519 | C20H21FN6O2 | 3.67 | 6 | 9 | 4.89 |
| CHEMBL3290351 | C24H27FN6O2 | 4.05 | 6 | 9 | |
| CHEMBL3290344 | C24H26N6O3 | 4.13 | 6 | 9 | |
| CHEMBL3608687 | C16H15ClN6OS | 4.18 | 6 | 9 | |
| CHEMBL4094682 | C22H27ClN6O2 | 4.34 | 6 | 9 | |
| CHEMBL3218891 | C27H32N6O3 | 4.37 | 6 | 9 | |
| CHEMBL2441417 | C24H26N6O3 | 4.89 | 6 | 9 | |

**Table S6** (Cont.d)

| | | | | | |
|---|---|---|---|---|---|
| CHEMBL502288 | C23H22N6O2S | 5.11 | 6 | 9 | |
| CHEMBL402015 | C27H27BrN6OS | 5.20 | 6 | 9 | |
| CHEMBL514042 | C25H38N6O3 | 6.05 | 6 | 9 | |
| CHEMBL498411 | C22H19Cl3N6 | 6.46 | 6 | 9 | |
| CHEMBL3422973 | C27H28N6O3 | 8.68 | 6 | 9 | |
| CHEMBL3422970 | C27H28N6O3 | 9.06 | 6 | 9 | |
| CHEMBL1916543 | C23H28N6O4 | 3.69 | 6 | 10 | 4.66 |
| CHEMBL4112037 | C16H14F2N6OS | 4.16 | 6 | 10 | |
| CHEMBL3221500 | C22H21ClN6O3 | 4.20 | 6 | 10 | |
| CHEMBL3794265 | C23H25FN6O3 | 4.38 | 6 | 10 | |
| CHEMBL3221488 | C23H23ClN6O3 | 4.50 | 6 | 10 | |
| CHEMBL3425929 | C28H33FN6O3 | 4.51 | 6 | 10 | |
| CHEMBL2041188 | C32H30N6O4 | 4.64 | 6 | 10 | |
| CHEMBL2324520 | C20H20F2N6O2 | 4.68 | 6 | 10 | |
| CHEMBL563791 | C28H29F3N6O | 4.90 | 6 | 10 | |
| CHEMBL513921 | C23H34N6O4 | 5.24 | 6 | 10 | |
| CHEMBL3408394 | C37H24N6O2S2 | 5.24 | 6 | 10 | |
| CHEMBL562285 | C24H27N3O2 | 5.27 | 6 | 10 | |
| CHEMBL1081867 | C22H23F3N6S | 6.33 | 6 | 10 | |
| CHEMBL390649 | C28H31F3N6S | 7.50 | 6 | 10 | |
| CHEMBL3985847 | C21H26N6O4S | 2.40 | 6 | 11 | 4.70 |
| CHEMBL3964789 | C21H22F4N6O | 3.17 | 6 | 11 | |
| CHEMBL2165057 | C25H30N6O5 | 4.06 | 6 | 11 | |
| CHEMBL3703025 | C22H25F3N6O2 | 4.50 | 6 | 11 | |
| CHEMBL3407784 | C20H19ClN6O3S | 4.89 | 6 | 11 | |
| CHEMBL3318984 | C23H23F3N6O2 | 5.00 | 6 | 11 | |
| CHEMBL192 | C22H30N6O4S | 5.48 | 6 | 11 | |
| CHEMBL402016 | C28H27F3N6OS | 6.00 | 6 | 11 | |
| CHEMBL254316 | C20H21FN6O5 | 2.50 | 6 | 12 | 4.60 |
| CHEMBL3608744 | C28H31ClN6O4S | 4.09 | 6 | 12 | |

**Table S6** (Cont.d)

| | | | | | |
|---|---|---|---|---|---|
| CHEMBL3425807 | C25H25FN6O5 | 4.52 | 6 | 12 | |
| CHEMBL2206791 | C23H21F3N6O3 | 4.58 | 6 | 12 | |
| CHEMBL2336323 | C26H30F2N6O4 | 4.62 | 6 | 12 | |
| CHEMBL2064657 | C28H23ClF2N6O3 | 4.91 | 6 | 12 | |
| CHEMBL3288030 | C44H53FN6O5 | 5.20 | 6 | 12 | |
| CHEMBL2441412 | C24H23F3N6O3 | 5.29 | 6 | 12 | |
| CHEMBL195378 | C22H21N7O2 | 4.30 | 7 | 9 | 5.60 |
| CHEMBL474484 | C26H34ClN7O | 4.53 | 7 | 9 | |
| CHEMBL2147022 | C21H22ClN7O | 4.70 | 7 | 9 | |
| CHEMBL2180070 | C25H31N7OS | 5.60 | 7 | 9 | |
| CHEMBL3593771 | C23H31N7O2 | 5.73 | 7 | 9 | |
| CHEMBL262341 | C28H25N7O2 | 5.85 | 7 | 9 | |
| CHEMBL399525 | C34H31N7O2 | 5.91 | 7 | 9 | |
| CHEMBL484816 | C22H27N7O4 | 4.09 | 7 | 11 | 4.90 |
| CHEMBL2325729 | C23H22ClN7O3 | 4.40 | 7 | 11 | |
| CHEMBL3262358 | C23H20ClN7O3 | 4.90 | 7 | 11 | |
| CHEMBL245119 | C31H27N7O3S | 5.82 | 7 | 11 | |
| CHEMBL429761 | C28H28F3N7S | 6.60 | 7 | 11 | |

**Table S7.** Investigation of the influence of Sulfur atom on the $pIC_{50}$ values.

| Name | Molecular formula | pIC50 | S | Heteroatoms | median |
|---|---|---|---|---|---|
| CHEMBL284348 | C5H6N2 | 2.36 | 0 | 2 | 5.16 |
| CHEMBL3 | C10H14N2 | 3.61 | 0 | 2 | |
| CHEMBL1086033 | C17H22N2 | 3.67 | 0 | 2 | |
| CHEMBL87045 | C18H18N2 | 5.06 | 0 | 2 | |
| CHEMBL459176 | C20H22N2 | 5.16 | 0 | 2 | |
| CHEMBL1684059 | C20H28ClN | 5.37 | 0 | 2 | |

**Table S7** (cont.d)

| | | | | | |
|---|---|---|---|---|---|
| CHEMBL11 | C19H24N2 | 5.47 | 0 | 2 | |
| CHEMBL83 | C26H29NO | 6.01 | 0 | 2 | |
| CHEMBL1671894 | C27H26N2 | 7.75 | 0 | 2 | |
| CHEMBL1087 | C15H22N2O | 3.81 | 0 | 3 | 5.60 |
| CHEMBL3314420 | C15H13ClFN | 4.30 | 0 | 3 | |
| CHEMBL1684047 | C16H23Cl2N | 4.40 | 0 | 3 | |
| CHEMBL1083707 | C20H24N2O | 4.67 | 0 | 3 | |
| CHEMBL513258 | C19H19FN2 | 5.05 | 0 | 3 | |
| CHEMBL1945694 | C16H15ClN2 | 5.16 | 0 | 3 | |
| CHEMBL2325209 | $C26H28NO2^{+}$ | 5.54 | 0 | 3 | |
| CHEMBL191054 | C13H17ClN2 | 5.60 | 0 | 3 | |
| CHEMBL497171 | C15H17N3 | 5.80 | 0 | 3 | |
| CHEMBL299390 | C19H19ClFN | 5.83 | 0 | 3 | |
| CHEMBL457930 | C20H19N3 | 5.85 | 0 | 3 | |
| CHEMBL505 | C16H19ClN2 | 5.96 | 0 | 3 | |
| CHEMBL2324248 | $C20H20NO2^{+}$ | 6.12 | 0 | 3 | |
| CHEMBL299499 | C17H22IN3 | 6.31 | 0 | 3 | |
| CHEMBL607 | C15H21NO2 | 6.49 | 0 | 3 | |
| CHEMBL70 | C17H19NO3 | 3.00 | 0 | 4 | 5.01 |
| CHEMBL1671896 | C11H14F3N | 3.27 | 0 | 4 | |
| CHEMBL640 | C13H21N3O | 3.86 | 0 | 4 | |
| CHEMBL219803 | C17H25N3O | 3.94 | 0 | 4 | |
| CHEMBL16 | C15H12N2O2 | 4.00 | 0 | 4 | |
| CHEMBL3356251 | C24H36N2O2 | 4.01 | 0 | 4 | |
| CHEMBL219128 | C17H33N3O | 4.17 | 0 | 4 | |
| CHEMBL3786542 | C26H26N2O2 | 4.21 | 0 | 4 | |
| CHEMBL3356249 | C25H38N2O2 | 4.30 | 0 | 4 | |
| CHEMBL219074 | C20H31N3O | 4.42 | 0 | 4 | |
| CHEMBL3314426 | C15H14F2N2 | 4.66 | 0 | 4 | |

**Table S7** (Cont.d)

| | | | | | |
|---|---|---|---|---|---|
| CHEMBL1086273 | C19H23FN2O | 4.68 | 0 | 4 | |
| CHEMBL1089159 | C22H23N3O | 4.70 | 0 | 4 | |
| CHEMBL2333646 | C19H23BrN2O | 4.89 | 0 | 4 | |
| CHEMBL3220615 | C19H21NO3 | 4.89 | 0 | 4 | |
| CHEMBL3185736 | C22H20N2O2 | 4.91 | 0 | 4 | |
| CHEMBL517 | C21H29N3O | 4.92 | 0 | 4 | |
| CHEMBL4162513 | C35H34N2O2 | 5.10 | 0 | 4 | |
| CHEMBL562948 | C22H33N3O | 5.40 | 0 | 4 | |
| CHEMBL1083407 | C21H22ClN3 | 5.44 | 0 | 4 | |
| CHEMBL1090175 | C20H25N3O | 5.47 | 0 | 4 | |
| CHEMBL53866 | C17H15ClFNO | 5.66 | 0 | 4 | |
| CHEMBL1935443 | C20H23N3O | 5.68 | 0 | 4 | |
| CHEMBL4450665 | C23H28N2O2 | 5.92 | 0 | 4 | |
| CHEMBL3696120 | C22H26ClN3 | 5.96 | 0 | 4 | |
| CHEMBL460893 | C23H25N3O | 6.13 | 0 | 4 | |
| CHEMBL1346 | C28H30N2O2 | 7.10 | 0 | 4 | |
| CHEMBL2046892 | C22H22F2N2 | 7.40 | 0 | 4 | |
| CHEMBL96153 | C24H31NO3 | 7.52 | 0 | 4 | |
| CHEMBL1223951 | C21H24N2O2 | 9.21 | 0 | 4 | |
| CHEMBL1257938 | C22H26N2O2 | 9.21 | 0 | 4 | |
| CHEMBL1257937 | C22H26N2O2 | 9.28 | 0 | 4 | |
| CHEMBL1257820 | C23H28N2O2 | 9.60 | 0 | 4 | |
| CHEMBL1257821 | C22H26N2O2 | 9.85 | 0 | 4 | |
| CHEMBL1097 | C15H23N3O2 | 4.00 | 0 | 5 | 5.22 |
| CHEMBL4640366 | C33H34N2O3 | 4.02 | 0 | 5 | |
| CHEMBL1091879 | C19H25N3O2 | 4.12 | 0 | 5 | |
| CHEMBL561138 | C23H27N3O2 | 4.14 | 0 | 5 | |
| CHEMBL1258006 | C22H23N3O2 | 4.27 | 0 | 5 | |
| CHEMBL3787069 | C26H26N2O3 | 4.35 | 0 | 5 | |

**Table S7** (Cont.d)

| | | | | |
|---|---|---|---|---|
| CHEMBL2324239 | $C24H18Cl2NO2^{+}$ | 4.51 | 0 | 5 |
| CHEMBL1094041 | C29H29N3O2 | 4.55 | 0 | 5 |
| CHEMBL4213196 | C25H34N4O | 4.62 | 0 | 5 |
| CHEMBL914 | C32H39NO4 | 4.67 | 0 | 5 |
| CHEMBL217133 | C18H22N4O | 4.68 | 0 | 5 |
| CHEMBL429115 | C20H24N4O | 4.71 | 0 | 5 |
| CHEMBL2441636 | C22H33N3O2 | 4.72 | 0 | 5 |
| CHEMBL1080790 | C16H22Cl2N2O | 4.78 | 0 | 5 |
| CHEMBL4096683 | C20H22N4O | 4.94 | 0 | 5 |
| CHEMBL237191 | C25H31N5 | 4.98 | 0 | 5 |
| CHEMBL565829 | C15H14N4O | 5.01 | 0 | 5 |
| CHEMBL2177304 | C27H24FN3O | 5.07 | 0 | 5 |
| CHEMBL487063 | C23H27N3O2 | 5.10 | 0 | 5 |
| CHEMBL378547 | C21H21F2NO2 | 5.11 | 0 | 5 |
| CHEMBL1080969 | C16H22Cl2N2O | 5.11 | 0 | 5 |
| CHEMBL485242 | C25H29N3O2 | 5.14 | 0 | 5 |
| CHEMBL478615 | C23H27N3O2 | 5.17 | 0 | 5 |
| CHEMBL476579 | C24H29N3O2 | 5.18 | 0 | 5 |
| CHEMBL479242 | C23H27N3O2 | 5.18 | 0 | 5 |
| CHEMBL3121096 | C20H25N3O2 | 5.22 | 0 | 5 |
| CHEMBL498572 | C19H22N2O3 | 5.23 | 0 | 5 |
| CHEMBL1819137 | C22H25N3O2 | 5.23 | 0 | 5 |
| CHEMBL488249 | C24H27N3O2 | 5.25 | 0 | 5 |
| CHEMBL4100591 | C27H34F2N2O | 5.29 | 0 | 5 |
| CHEMBL384400 | C27H36ClN3O | 5.48 | 0 | 5 |
| CHEMBL487064 | C23H27N3O2 | 5.55 | 0 | 5 |
| CHEMBL1110 | C17H18N4O | 5.64 | 0 | 5 |
| CHEMBL478616 | C23H27N3O2 | 5.80 | 0 | 5 |
| CHEMBL196983 | C21H27NO4 | 6.10 | 0 | 5 |

**Table S7** (Cont.d)

| | | | | | |
|---|---|---|---|---|---|
| CHEMBL562833 | C22H30ClN3O | 6.10 | 0 | 5 | |
| CHEMBL58387 | C31H38N4O | 6.24 | 0 | 5 | |
| CHEMBL3775729 | C26H28N4O | 6.28 | 0 | 5 | |
| CHEMBL376488 | C32H31BrN2O2 | 6.43 | 0 | 5 | |
| CHEMBL1671893 | C23H30N2O3 | 6.60 | 0 | 5 | |
| CHEMBL1087493 | C17H26N4O | 6.63 | 0 | 5 | |
| CHEMBL42 | C18H19ClN4 | 6.72 | 0 | 5 | |
| CHEMBL998 | C22H23ClN2O2 | 6.77 | 0 | 5 | |
| CHEMBL2324243 | $C26H32NO4^{+}$ | 6.80 | 0 | 5 | |
| CHEMBL2324245 | $C24H26Cl2NO2^{+}$ | 7.22 | 0 | 5 | |
| CHEMBL572163 | C26H31NO4 | 7.42 | 0 | 5 | |
| CHEMBL61301 | C19H21FN4 | 7.56 | 0 | 5 | |
| CHEMBL1642487 | C28H28N2O3 | 9.09 | 0 | 5 | |
| CHEMBL1642479 | C28H28N2O3 | 9.17 | 0 | 5 | |
| CHEMBL259732 | C20H27N3O2 | 9.19 | 0 | 5 | |
| CHEMBL410832 | C23H25N3O2 | 9.39 | 0 | 5 | |
| CHEMBL4787638 | C21H18ClN3O2 | 2.08 | 0 | 6 | **5.88** |
| CHEMBL270190 | C25H32N2O4 | 3.10 | 0 | 6 | |
| CHEMBL519266 | C22H28N2O4 | 3.11 | 0 | 6 | |
| CHEMBL1688 | C18H19ClN4O | 3.85 | 0 | 6 | |
| CHEMBL3741589 | C18H17Cl2NO3 | 4.11 | 0 | 6 | |
| CHEMBL1000 | C21H25ClN2O3 | 4.52 | 0 | 6 | |
| CHEMBL4212656 | C24H31N5O | 4.64 | 0 | 6 | |
| CHEMBL398612 | C26H35FN4O | 4.88 | 0 | 6 | |
| CHEMBL1823042 | C20H28FN3O2 | 5.00 | 0 | 6 | |
| CHEMBL494406 | C21H24ClN5 | 5.05 | 0 | 6 | |
| CHEMBL4067770 | C20H21FN4O | 5.11 | 0 | 6 | |
| CHEMBL1938433 | C23H24N2O4 | 5.42 | 0 | 6 | |
| CHEMBL2177904 | C23H16FN5 | 5.51 | 0 | 6 | |

**Table S7** (Cont.d)

| | | | | | |
|---|---|---|---|---|---|
| CHEMBL1823047 | C22H30N4O2 | 5.54 | 0 | 6 | |
| CHEMBL4065208 | C23H22N4O2 | 5.56 | 0 | 6 | |
| CHEMBL2146805 | C20H21ClN4O | 5.60 | 0 | 6 | |
| CHEMBL2177905 | C24H16F2N4 | 5.80 | 0 | 6 | |
| CHEMBL2147024 | C22H18N6 | 5.96 | 0 | 6 | |
| CHEMBL563926 | C21H26ClN3O2 | 6.20 | 0 | 6 | |
| CHEMBL1083463 | C26H26Cl2N4 | 6.22 | 0 | 6 | |
| CHEMBL4081790 | C25H19N5O | 6.30 | 0 | 6 | |
| CHEMBL723 | C24H26N2O4 | 6.46 | 0 | 6 | |
| CHEMBL4096145 | C22H27N5O | 6.52 | 0 | 6 | |
| CHEMBL3774817 | C25H25FN4O | 6.59 | 0 | 6 | |
| CHEMBL522456 | CHEMBL522456 | 7.00 | 0 | 6 | |
| CHEMBL3775282 | C26H28N4O2 | 7.16 | 0 | 6 | |
| CHEMBL1108 | C22H22FN3O2 | 7.49 | 0 | 6 | |
| CHEMBL329067 | C32H33N5O | 7.62 | 0 | 6 | |
| CHEMBL12780 | C24H27FN4O | 8.16 | 0 | 6 | |
| CHEMBL2146810 | C23H27ClN4O | 8.40 | 0 | 6 | |
| CHEMBL1423 | C28H29F2N3O | 8.52 | 0 | 6 | |
| CHEMBL296419 | C28H31FN4O | 9.05 | 0 | 6 | |
| CHEMBL1258503 | C22H25N3O3 | 9.14 | 0 | 6 | |
| CHEMBL1258280 | C22H24N2O4 | 9.21 | 0 | 6 | |
| CHEMBL374731 | C10H14N2O5 | 2.30 | 0 | 7 | **5.21** |
| CHEMBL8 | C17H18FN3O3 | 3.02 | 0 | 7 | |
| CHEMBL4855895 | C21H24FN3O3 | 3.53 | 0 | 7 | |
| CHEMBL22 | C14H18N4O3 | 3.62 | 0 | 7 | |
| CHEMBL3582294 | C25H18F2N4O | 4.03 | 0 | 7 | |
| CHEMBL4224807 | C14H16N6O | 4.03 | 0 | 7 | |
| CHEMBL388978 | C28H26N4O3 | 4.04 | 0 | 7 | |
| CHEMBL4847738 | C20H23N3O4 | 4.05 | 0 | 7 | |

**Table S7** (Cont.d)

| | | | | |
|---|---|---|---|---|
| CHEMBL4781551 | C24H21ClN4O2 | 4.08 | 0 | 7 |
| CHEMBL3329814 | C21H28N2O5 | 4.09 | 0 | 7 |
| CHEMBL4127458 | C22H22ClN5O | 4.22 | 0 | 7 |
| CHEMBL474484 | C13H21N7 | 4.28 | 0 | 7 |
| CHEMBL583 | C19H22FN3O3 | 4.30 | 0 | 7 |
| CHEMBL218836 | C18H24F3N3O | 4.40 | 0 | 7 |
| CHEMBL2158823 | C24H28Cl2N2O3 | 4.50 | 0 | 7 |
| CHEMBL1957012 | C26H29N5O2 | 4.60 | 0 | 7 |
| CHEMBL512975 | C21H23ClFN5 | 4.64 | 0 | 7 |
| CHEMBL2333636 | C24H28F4N2O | 4.75 | 0 | 7 |
| CHEMBL2180842 | C17H22FN5O | 4.80 | 0 | 7 |
| CHEMBL2314058 | C25H34F3N3O | 4.83 | 0 | 7 |
| CHEMBL523374 | C22H23N5O2 | 4.89 | 0 | 7 |
| CHEMBL3115194 | C26H32BrN3O3 | 4.90 | 0 | 7 |
| CHEMBL2333616 | C20H22ClF3N2O | 4.92 | 0 | 7 |
| CHEMBL3218884 | C28H33N5O2 | 4.96 | 0 | 7 |
| CHEMBL1957792 | C27H30ClN3O3 | 4.99 | 0 | 7 |
| CHEMBL1092106 | C20H24ClFN4O | 5.04 | 0 | 7 |
| CHEMBL2333607 | C25H30F4N2O | 5.05 | 0 | 7 |
| CHEMBL2333615 | C23H26F4N2O | 5.05 | 0 | 7 |
| CHEMBL498042 | C24H23N5O2 | 5.08 | 0 | 7 |
| CHEMBL3703271 | C22H23N3O4 | 5.08 | 0 | 7 |
| CHEMBL3262625 | C23H26N6O | 5.09 | 0 | 7 |
| CHEMBL246815 | C22H26FN3O3 | 5.13 | 0 | 7 |
| CHEMBL523821 | C21H15F2N5 | 5.16 | 0 | 7 |
| CHEMBL398478 | C25H28FN3O3 | 5.21 | 0 | 7 |
| CHEMBL2178522 | C23H19FN6 | 5.28 | 0 | 7 |
| CHEMBL2158815 | C24H28Cl2N2O3 | 5.30 | 0 | 7 |
| CHEMBL497048 | C27H23N5O2 | 5.36 | 0 | 7 |

Table S6 (Cont.d)

| | | | | |
|---|---|---|---|---|
| CHEMBL2147022 | C20H17N7 | 5.37 | 0 | 7 |
| CHEMBL2336331 | C24H29N5O2 | 5.40 | 0 | 7 |
| CHEMBL1091834 | C25H26ClFN4O | 5.42 | 0 | 7 |
| CHEMBL1813015 | C20H20N6O | 5.42 | 0 | 7 |
| CHEMBL1939739 | C32H35FN4O2 | 5.44 | 0 | 7 |
| CHEMBL250533 | C25H28ClN3O3 | 5.48 | 0 | 7 |
| CHEMBL4102707 | C20H20ClF4NO | 5.54 | 0 | 7 |
| CHEMBL2147021 | C20H15FN6 | 5.57 | 0 | 7 |
| CHEMBL246634 | C21H23FN2O4 | 5.58 | 0 | 7 |
| CHEMBL1098847 | C24H21Cl3N2O2 | 5.58 | 0 | 7 |
| CHEMBL2151218 | C25H22FN5O | 5.70 | 0 | 7 |
| CHEMBL3764774 | C28H28N4O3 | 5.70 | 0 | 7 |
| CHEMBL3086038 | C30H33BrClN3O2 | 5.82 | 0 | 7 |
| CHEMBL3806193 | C19H23Cl2N3O2 | 5.84 | 0 | 7 |
| CHEMBL2146812 | C19H20ClN5O | 5.90 | 0 | 7 |
| CHEMBL17423 | C22H25N3O4 | 5.96 | 0 | 7 |
| CHEMBL3775387 | C25H27N5O2 | 5.99 | 0 | 7 |
| CHEMBL3806153 | C18H21ClFN3O2 | 6.01 | 0 | 7 |
| CHEMBL2387265 | C31H29FN2O4 | 6.09 | 0 | 7 |
| CHEMBL3775050 | C25H24F2N4O | 6.20 | 0 | 7 |
| CHEMBL45816 | C29H38FN3O3 | 6.24 | 0 | 7 |
| CHEMBL501480 | C29H25N5O2 | 6.50 | 0 | 7 |
| CHEMBL1084400 | C20H21F3N4 | 6.80 | 0 | 7 |
| CHEMBL526466 | C25H23N5O2 | 7.10 | 0 | 7 |
| CHEMBL2146854 | C23H26N4O3 | 7.33 | 0 | 7 |
| CHEMBL1107 | C26H30Cl2F3NO | 7.40 | 0 | 7 |
| CHEMBL384487 | C24H23FN2O4 | 7.69 | 0 | 7 |
| CHEMBL4279819 | C16H18N6O | 8.07 | 0 | 7 |
| CHEMBL217593 | C23H23FN2O4 | 8.27 | 0 | 7 |

Table S6 (Cont.d)

| | | | | | |
|---|---|---|---|---|---|
| CHEMBL12713 | C24H26ClFN4O | 8.57 | 0 | 7 | |
| CHEMBL33 | C18H20FN3O4 | 3.04 | 0 | 8 | **5.10** |
| CHEMBL399726 | C20H24N6O2 | 3.78 | 0 | 8 | |
| CHEMBL31 | C19H22FN3O4 | 3.89 | 0 | 8 | |
| CHEMBL3804950 | C20H21F2N5O | 4.01 | 0 | 8 | |
| CHEMBL4858551 | C19H19FN6O | 4.01 | 0 | 8 | |
| CHEMBL520463 | C28H24N4O4 | 4.05 | 0 | 8 | |
| CHEMBL485123 | C20H23F3N4O | 4.07 | 0 | 8 | |
| CHEMBL32 | C21H24FN3O4 | 4.10 | 0 | 8 | |
| CHEMBL2333619 | C22H27F4N3O | 4.10 | 0 | 8 | |
| CHEMBL2180070 | C19H23N7O | 4.27 | 0 | 8 | |
| CHEMBL193 | C17H18N2O6 | 4.30 | 0 | 8 | |
| CHEMBL2382343 | C29H32F2N2O4 | 4.32 | 0 | 8 | |
| CHEMBL2314064 | C24H32F3N3O2 | 4.41 | 0 | 8 | |
| CHEMBL540634 | C26H31Cl2N3O3 | 4.50 | 0 | 8 | |
| CHEMBL3577935 | C23H34F3N3O2 | 4.50 | 0 | 8 | |
| CHEMBL3787345 | C25H28N4O4 | 4.51 | 0 | 8 | |
| CHEMBL2164565 | C19H18F3N3O2 | 4.63 | 0 | 8 | |
| CHEMBL3216126 | C25H35Cl2N5O | 4.65 | 0 | 8 | |
| CHEMBL602875 | C26H34N4O4 | 4.72 | 0 | 8 | |
| CHEMBL2178221 | C28H29ClN4O3 | 4.82 | 0 | 8 | |
| CHEMBL3612814 | C24H28N4O4 | 4.85 | 0 | 8 | |
| CHEMBL3593771 | C19H19N7O | 4.87 | 0 | 8 | |
| CHEMBL1093696 | C24H25ClFN5O | 4.92 | 0 | 8 | |
| CHEMBL515001 | C21H25FN6O | 4.96 | 0 | 8 | |
| CHEMBL1084617 | C24H28N4O4 | 4.96 | 0 | 8 | |
| CHEMBL3964723 | C23H27Cl2N3O3 | 5.00 | 0 | 8 | |
| CHEMBL465417 | C25H23N5O3 | 5.04 | 0 | 8 | |
| CHEMBL3786346 | C25H26N4O4 | 5.05 | 0 | 8 | |

Table S6 (Cont.d)

| | | | | | |
|---|---|---|---|---|---|
| CHEMBL2325997 | C20H23ClN6O | 5.14 | 0 | 8 | |
| CHEMBL506163 | C22H19ClFN5O | 5.20 | 0 | 8 | |
| CHEMBL262341 | C33H29N7O | 5.25 | 0 | 8 | |
| CHEMBL2158771 | C27H31Cl2N3O3 | 5.30 | 0 | 8 | |
| CHEMBL2181489 | C23H20F3N3O2 | 5.52 | 0 | 8 | |
| CHEMBL3218650 | C27H30ClN5O2 | 5.60 | 0 | 8 | |
| CHEMBL3703237 | C19H17ClN4O3 | 5.66 | 0 | 8 | |
| CHEMBL399525 | C19H23N7O | 5.75 | 0 | 8 | |
| CHEMBL2178521 | C24H21FN6O | 5.75 | 0 | 8 | |
| CHEMBL3415593 | C19H26N6O2 | 5.77 | 0 | 8 | |
| CHEMBL2387229 | C26H26ClFN2O4 | 5.80 | 0 | 8 | |
| CHEMBL478462 | C25H38N6O2 | 5.84 | 0 | 8 | |
| CHEMBL1956991 | C23H26ClFN4O2 | 5.89 | 0 | 8 | |
| CHEMBL2441431 | C22H23N3O5 | 5.89 | 0 | 8 | |
| CHEMBL1621 | C23H27FN4O3 | 6.00 | 0 | 8 | |
| CHEMBL3318989 | C23H24ClN5O2 | 6.00 | 0 | 8 | |
| CHEMBL515025 | C24H36N6O2 | 6.15 | 0 | 8 | |
| CHEMBL478011 | C25H38N6O2 | 6.38 | 0 | 8 | |
| CHEMBL94454 | C24H25FN6O | 6.46 | 0 | 8 | |
| CHEMBL453894 | C25H26F3N3O2 | 6.46 | 0 | 8 | |
| CHEMBL233047 | C20H32Cl2N6 | 6.52 | 0 | 8 | |
| CHEMBL2146813 | C38H48ClN5O2 | 6.80 | 0 | 8 | |
| CHEMBL568571 | C27H26F3N3O2 | 6.85 | 0 | 8 | |
| CHEMBL128007 | C28H28Cl2N4O2 | 7.00 | 0 | 8 | |
| CHEMBL525413 | C29H24ClN5O2 | 7.09 | 0 | 8 | |
| CHEMBL2146853 | C20H23N5O3 | 7.60 | 0 | 8 | |
| CHEMBL214021 | C23H21FN2O5 | 7.82 | 0 | 8 | |
| CHEMBL217789 | C23H21ClN2O5 | 8.44 | 0 | 8 | |
| CHEMBL2324519 | C20H21FN6O2 | 3.67 | 0 | 9 | **5.15** |

**Table S6** (Cont.d)

| | | | | |
|---|---|---|---|---|
| CHEMBL3799831 | C25H26FN3O5 | 4.03 | 0 | 9 |
| CHEMBL3290351 | C24H27FN6O2 | 4.05 | 0 | 9 |
| CHEMBL216323 | C23H19FN2O6 | 4.07 | 0 | 9 |
| CHEMBL4092041 | C29H31F3N2O4 | 4.07 | 0 | 9 |
| CHEMBL3290344 | C24H26N6O3 | 4.13 | 0 | 9 |
| CHEMBL248296 | C27H31F2N3O4 | 4.19 | 0 | 9 |
| CHEMBL2165068 | C25H26FN5O3 | 4.24 | 0 | 9 |
| CHEMBL484816 | C22H21N7O2 | 4.30 | 0 | 9 |
| CHEMBL4094682 | C22H27ClN6O2 | 4.34 | 0 | 9 |
| CHEMBL3218891 | C27H32N6O3 | 4.37 | 0 | 9 |
| CHEMBL2325729 | C26H34ClN7O | 4.53 | 0 | 9 |
| CHEMBL3650850 | C21H23N5O4 | 4.68 | 0 | 9 |
| CHEMBL3262358 | C21H22ClN7O | 4.70 | 0 | 9 |
| CHEMBL3612928 | C24H28N4O5 | 4.70 | 0 | 9 |
| CHEMBL3786311 | C26H30N4O5 | 4.80 | 0 | 9 |
| CHEMBL596700 | C20H23F3N4O2 | 4.88 | 0 | 9 |
| CHEMBL2441417 | C24H26N6O3 | 4.89 | 0 | 9 |
| CHEMBL255968 | C25H24ClN5O3 | 5.00 | 0 | 9 |
| CHEMBL1095476 | C28H25Cl3N4O2 | 5.06 | 0 | 9 |
| CHEMBL3416021 | C20H24N8O | 5.07 | 0 | 9 |
| CHEMBL589833 | C24H25Cl2FN4O2 | 5.10 | 0 | 9 |
| CHEMBL3612926 | C25H30N4O5 | 5.11 | 0 | 9 |
| CHEMBL2164393 | C26H24F3N3O3 | 5.14 | 0 | 9 |
| CHEMBL2170611 | C17H21F3N4O2 | 5.16 | 0 | 9 |
| CHEMBL429458 | C25H30FN3O5 | 5.20 | 0 | 9 |
| CHEMBL4075071 | C19H16F3N5O | 5.25 | 0 | 9 |
| CHEMBL4170966 | C24H21Cl2N3O4 | 5.33 | 0 | 9 |
| CHEMBL2164389 | C24H20F3N3O3 | 5.34 | 0 | 9 |
| CHEMBL255645 | C28H31Cl2N5O2 | 5.36 | 0 | 9 |

**Table S6** (Cont.d)

| | | | | | |
|---|---|---|---|---|---|
| CHEMBL1927161 | C24H29N5O4 | 5.40 | 0 | 9 | |
| CHEMBL489324 | C28H42ClF2N5O | 5.68 | 0 | 9 | |
| CHEMBL3237444 | C23H25ClN8 | 5.70 | 0 | 9 | |
| CHEMBL4213486 | C19H15F2N5O2 | 5.70 | 0 | 9 | |
| CHEMBL429761 | C23H31N7O2 | 5.73 | 0 | 9 | |
| CHEMBL3318999 | C28H25N7O2 | 5.85 | 0 | 9 | |
| CHEMBL2177305 | C28H23ClFN5O2 | 5.89 | 0 | 9 | |
| CHEMBL446966 | C34H31N7O2 | 5.91 | 0 | 9 | |
| CHEMBL550410 | C25H26F4N4O | 5.92 | 0 | 9 | |
| CHEMBL557306 | C21H24ClF3N2O3 | 5.99 | 0 | 9 | |
| CHEMBL514042 | C25H38N6O3 | 6.05 | 0 | 9 | |
| CHEMBL549635 | C29H33F4N3O2 | 6.20 | 0 | 9 | |
| CHEMBL498411 | C22H19Cl3N6 | 6.46 | 0 | 9 | |
| CHEMBL217442 | C24H23FN2O6 | 7.80 | 0 | 9 | |
| CHEMBL1729 | C23H29ClFN3O4 | 8.19 | 0 | 9 | |
| CHEMBL387178 | C23H20F2N2O5 | 8.21 | 0 | 9 | |
| CHEMBL3422973 | C27H28N6O3 | 8.68 | 0 | 9 | |
| CHEMBL3422970 | C27H28N6O3 | 9.06 | 0 | 9 | |
| CHEMBL2424928 | C24H26FN5O4 | 3.51 | 0 | 10 | **5.24** |
| CHEMBL1916543 | C23H28N6O4 | 3.69 | 0 | 10 | |
| CHEMBL605785 | C27H27FN4O5 | 4.02 | 0 | 10 | |
| CHEMBL3221500 | C22H21ClN6O3 | 4.20 | 0 | 10 | |
| CHEMBL2151322 | C19H20FN7O2 | 4.28 | 0 | 10 | |
| CHEMBL3794265 | C23H25FN6O3 | 4.38 | 0 | 10 | |
| CHEMBL247690 | C29H33F2N3O5 | 4.47 | 0 | 10 | |
| CHEMBL3221488 | C23H23ClN6O3 | 4.50 | 0 | 10 | |
| CHEMBL3425929 | C28H33FN6O3 | 4.51 | 0 | 10 | |
| CHEMBL2041188 | C32H30N6O4 | 4.64 | 0 | 10 | |
| CHEMBL2324520 | C20H20F2N6O2 | 4.68 | 0 | 10 | |

**Table S6** (Cont.d)

| | | | | | |
|---|---|---|---|---|---|
| CHEMBL2315921 | C26H28F3N5O2 | 4.77 | 0 | 10 | |
| CHEMBL563791 | C28H29F3N6O | 4.90 | 0 | 10 | |
| CHEMBL3400819 | C26H26FN5O4 | 4.91 | 0 | 10 | |
| CHEMBL256060 | C26H26ClN5O4 | 4.96 | 0 | 10 | |
| CHEMBL595944 | C27H26N8O2 | 4.98 | 0 | 10 | |
| CHEMBL2164365 | C23H19F3N4O3 | 5.04 | 0 | 10 | |
| CHEMBL2177736 | C19H23N9O | 5.07 | 0 | 10 | |
| CHEMBL3681314 | C26H25F2N7O | 5.07 | 0 | 10 | |
| CHEMBL513921 | C23H34N6O4 | 5.24 | 0 | 10 | |
| CHEMBL562285 | C24H27N3O2 | 5.27 | 0 | 10 | |
| CHEMBL2164047 | C24H19F4N3O3 | 5.28 | 0 | 10 | |
| CHEMBL560063 | C24H23Cl2F3N4O | 5.46 | 0 | 10 | |
| CHEMBL245642 | C25H25F2N3O5 | 5.50 | 0 | 10 | |
| CHEMBL551281 | C26H34F3N5O2 | 5.52 | 0 | 10 | |
| CHEMBL1236904 | C26H31BrN8O | 5.52 | 0 | 10 | |
| CHEMBL3604800 | C28H32FN5O4 | 5.52 | 0 | 10 | |
| CHEMBL452823 | C37H38N10 | 5.67 | 0 | 10 | |
| CHEMBL553196 | C32H32F4N4O2 | 5.76 | 0 | 10 | |
| CHEMBL3323074 | C32H28FN7O2 | 5.85 | 0 | 10 | |
| CHEMBL3323073 | C32H28FN7O2 | 6.20 | 0 | 10 | |
| CHEMBL537847 | C23H29Cl2N5O3 | 6.25 | 0 | 10 | |
| CHEMBL3605145 | C29H32FN5O4 | 6.30 | 0 | 10 | |
| CHEMBL3605140 | C27H30FN5O4 | 6.40 | 0 | 10 | |
| CHEMBL375330 | C34H38ClF3N4O2 | 6.77 | 0 | 10 | |
| CHEMBL4081080 | C25H17F6N3O | 7.17 | 0 | 10 | |
| CHEMBL428594 | C29H33F7N2O | 7.80 | 0 | 10 | |
| CHEMBL217707 | C23H19F3N2O5 | 8.25 | 0 | 10 | |
| CHEMBL213715 | C24H22F2N2O6 | 8.44 | 0 | 10 | |
| CHEMBL3964789 | C21H22F4N6O | 3.17 | 0 | 11 | **5.00** |

Table S6 (Cont.d)

| | | | | | |
|---|---|---|---|---|---|
| CHEMBL3400817 | C26H26FN5O5 | 3.76 | 0 | 11 | |
| CHEMBL2165057 | C25H30N6O5 | 4.06 | 0 | 11 | |
| CHEMBL1916544 | C22H27N7O4 | 4.09 | 0 | 11 | |
| CHEMBL3221490 | C23H22ClN7O3 | 4.40 | 0 | 11 | |
| CHEMBL3703025 | C22H25F3N6O2 | 4.50 | 0 | 11 | |
| CHEMBL1091218 | C29H25F3N2O6 | 4.58 | 0 | 11 | |
| CHEMBL2069410 | C23H19F4N3O4 | 4.60 | 0 | 11 | |
| CHEMBL4209441 | C26H23FN8O2 | 4.62 | 0 | 11 | |
| CHEMBL4211893 | C30H29FN8O2 | 4.87 | 0 | 11 | |
| CHEMBL3221503 | C23H20ClN7O3 | 4.90 | 0 | 11 | |
| CHEMBL2164375 | C25H26F3N3O5 | 4.96 | 0 | 11 | |
| CHEMBL2204270 | C24H26F4N4O3 | 5.00 | 0 | 11 | |
| CHEMBL3318984 | C23H23F3N6O2 | 5.00 | 0 | 11 | |
| CHEMBL508098 | C36H37N11 | 5.08 | 0 | 11 | |
| CHEMBL3353404 | C26H29ClN8O2 | 5.17 | 0 | 11 | |
| CHEMBL3746204 | C22H17F4N5O2 | 5.24 | 0 | 11 | |
| CHEMBL2331648 | C20H22F3N5O3 | 5.34 | 0 | 11 | |
| CHEMBL2147223 | C22H20ClF3N4O3 | 5.40 | 0 | 11 | |
| CHEMBL3422758 | C27H29F2N5O4 | 5.76 | 0 | 11 | |
| CHEMBL1097456 | C26H22Cl3F3N2O3 | 5.84 | 0 | 11 | |
| CHEMBL256154 | C27H32F6N4O | 5.85 | 0 | 11 | |
| CHEMBL3605123 | C30H36FN5O5 | 6.16 | 0 | 11 | |
| CHEMBL3422952 | C26H27F2N5O4 | 9.14 | 0 | 11 | |
| CHEMBL3422978 | C26H26F3N5O3 | 9.41 | 0 | 11 | |
| CHEMBL254316 | C20H21FN6O5 | 2.50 | 0 | 12 | **4.80** |
| CHEMBL3425799 | C26H25F2N5O5 | 3.48 | 0 | 12 | |
| CHEMBL451887 | C40H57N5O7 | 4.04 | 0 | 12 | |
| CHEMBL3342693 | C23H22F5N5O2 | 4.08 | 0 | 12 | |
| CHEMBL3425807 | C25H25FN6O5 | 4.52 | 0 | 12 | |

**Table S6** (Cont.d)

| | | | | | |
|---|---|---|---|---|---|
| CHEMBL2206791 | C23H21F3N6O3 | 4.58 | 0 | 12 | |
| CHEMBL2147316 | C22H19F5N4O3 | 4.60 | 0 | 12 | |
| CHEMBL2336323 | C26H30F2N6O4 | 4.62 | 0 | 12 | |
| CHEMBL2147303 | C22H19F5N4O3 | 4.70 | 0 | 12 | |
| CHEMBL402624 | C30H34F6N2O4 | 4.71 | 0 | 12 | |
| CHEMBL3221487 | C20H19ClN8O3 | 4.80 | 0 | 12 | |
| CHEMBL2064657 | C28H23ClF2N6O3 | 4.91 | 0 | 12 | |
| CHEMBL3605131 | C29H32FN5O6 | 4.91 | 0 | 12 | |
| CHEMBL3764895 | C29H31Cl2F3N4O3 | 5.13 | 0 | 12 | |
| CHEMBL3288030 | C44H53FN6O5 | 5.20 | 0 | 12 | |
| CHEMBL3764357 | C29H31Cl2F3N4O3 | 5.20 | 0 | 12 | |
| CHEMBL2441412 | C24H23F3N6O3 | 5.29 | 0 | 12 | |
| CHEMBL2207664 | C22H24Cl2F3N3O4 | 5.30 | 0 | 12 | |
| CHEMBL550535 | C27H33F6N5O | 5.41 | 0 | 12 | |
| CHEMBL3765012 | C30H31Cl2F3N4O3 | 5.51 | 0 | 12 | |
| CHEMBL3605006 | C31H34FN5O6 | 5.80 | 0 | 12 | |
| CHEMBL195378 | C23H24F4N6O3 | 4.23 | 0 | 13 | **5.29** |
| CHEMBL3605022 | C29H32FN7O5 | 4.54 | 0 | 13 | |
| CHEMBL370072 | C24H24F4N4O5 | 4.55 | 0 | 13 | |
| CHEMBL3218816 | C19H17F3N8O2 | 4.75 | 0 | 13 | |
| CHEMBL1099069 | C19H24F3N9O | 5.29 | 0 | 13 | |
| CHEMBL4079820 | C26H23F4N7O2 | 5.62 | 0 | 13 | |
| CHEMBL401608 | C27H31ClF6N4O2 | 6.73 | 0 | 13 | |
| CHEMBL4077588 | C29H29F4N7O2 | 7.00 | 0 | 13 | |
| CHEMBL3422957 | C27H27F4N5O4 | 8.77 | 0 | 13 | |
| CHEMBL4209354 | C15H20N2O2S | 4.07 | 1 | 5 | **5.90** |
| CHEMBL1125 | C16H18N4S | 4.90 | 1 | 5 | |
| CHEMBL2326478 | C18H18FN3S | 5.17 | 1 | 5 | |
| CHEMBL1956112 | C21H28N4S | 5.33 | 1 | 5 | |

**Table S6** (Cont.d)

| | | | | | |
|---|---|---|---|---|---|
| CHEMBL1224697 | C22H28N2O2S | 5.90 | 1 | 5 | |
| CHEMBL715 | C17H20N4S | 6.74 | 1 | 5 | |
| CHEMBL195180 | C24H27ClN2OS | 7.52 | 1 | 5 | |
| CHEMBL1257577 | C19H22N2O2S | 9.37 | 1 | 5 | |
| CHEMBL1257578 | C20H24N2O2S | 9.59 | 1 | 5 | |
| CHEMBL4100776 | C20H22N4OS | 5.23 | 1 | 6 | **5.85** |
| CHEMBL225036 | C24H26N4OS | 5.29 | 1 | 6 | |
| CHEMBL4089699 | C19H19N3O2S | 5.42 | 1 | 6 | |
| CHEMBL4104102 | C20H22N4OS | 5.61 | 1 | 6 | |
| CHEMBL3219616 | C24H30N2O3S | 5.80 | 1 | 6 | |
| CHEMBL1224699 | C21H26N2O3S | 5.90 | 1 | 6 | |
| CHEMBL561279 | C25H34N2O3S | 5.93 | 1 | 6 | |
| CHEMBL556100 | C22H25NO4S | 6.33 | 1 | 6 | |
| CHEMBL4173253 | C26H34N2O3S | 7.10 | 1 | 6 | |
| CHEMBL533 | C20H36N2O3S | 8.00 | 1 | 6 | |
| CHEMBL578834 | C21H25N5OS | 3.72 | 1 | 7 | **5.31** |
| CHEMBL2069925 | C21H21N5OS | 4.52 | 1 | 7 | |
| CHEMBL575241 | C21H26N6S | 4.71 | 1 | 7 | |
| CHEMBL3973288 | C20H22FN3O2S | 4.84 | 1 | 7 | |
| CHEMBL560386 | C23H29NO5S | 5.31 | 1 | 7 | |
| CHEMBL243901 | C23H31N5OS | 5.90 | 1 | 7 | |
| CHEMBL1079578 | C28H31N5OS | 6.20 | 1 | 7 | |
| CHEMBL708 | C21H21ClN4OS | 6.92 | 1 | 7 | |
| CHEMBL584766 | C23H24FN5S | 7.00 | 1 | 7 | |
| CHEMBL256653 | C23H26N6OS | 4.30 | 1 | 8 | **5.40** |
| CHEMBL1939742 | C30H37N5O2S | 4.75 | 1 | 8 | |
| CHEMBL257901 | C28H30N6OS | 4.90 | 1 | 8 | |
| CHEMBL270239 | C30H34N6OS | 4.90 | 1 | 8 | |
| CHEMBL2440407 | C23H26N2O5S | 5.02 | 1 | 8 | |

Table S6 (Cont.d)

| | | | | | |
|---|---|---|---|---|---|
| CHEMBL3287218 | C29H32N6OS | 5.16 | 1 | 8 | |
| CHEMBL207220 | C24H40N4O3S | 5.40 | 1 | 8 | |
| CHEMBL270852 | C32H38N6OS | 5.40 | 1 | 8 | |
| CHEMBL272086 | C32H38N6OS | 5.60 | 1 | 8 | |
| CHEMBL2207738 | C23H26N4O3S | 5.60 | 1 | 8 | |
| CHEMBL1080489 | C22H27N5O2S | 5.80 | 1 | 8 | |
| CHEMBL408169 | C28H32N6OS | 6.40 | 1 | 8 | |
| CHEMBL244280 | C26H28FN5OS | 6.60 | 1 | 8 | |
| CHEMBL244278 | C26H28ClN5OS | 7.10 | 1 | 8 | |
| CHEMBL397429 | C28H30N6OS | 7.30 | 1 | 8 | |
| CHEMBL3608687 | C16H15ClN6OS | 4.18 | 1 | 9 | **5.23** |
| CHEMBL4084170 | C15H18F5N3S | 4.69 | 1 | 9 | |
| CHEMBL3422244 | C18H18F3N3O2S | 4.71 | 1 | 9 | |
| CHEMBL1083118 | C17H19F2N3O3S | 4.89 | 1 | 9 | |
| CHEMBL1091664 | C28H25ClN2O5S | 5.00 | 1 | 9 | |
| CHEMBL502288 | C23H22N6O2S | 5.11 | 1 | 9 | |
| CHEMBL402015 | C27H27BrN6OS | 5.20 | 1 | 9 | |
| CHEMBL4243499 | C19H16F3N5O | 5.25 | 1 | 9 | |
| CHEMBL2158625 | C20H19Cl2N5OS | 5.44 | 1 | 9 | |
| CHEMBL244083 | C29H36N4O4S | 5.60 | 1 | 9 | |
| CHEMBL245119 | C25H31N7OS | 5.60 | 1 | 9 | |
| CHEMBL2158629 | C20H21Cl2N5OS | 5.85 | 1 | 9 | |
| CHEMBL1081747 | C21H23ClFN5OS | 6.40 | 1 | 9 | |
| CHEMBL3983509 | C20H20ClFN4O2S | 6.70 | 1 | 9 | |
| CHEMBL3921669 | C23H26N4O5S | 2.22 | 1 | 10 | **5.90** |
| CHEMBL2440387 | C25H23ClN4O4S | 3.38 | 1 | 10 | |
| CHEMBL4112037 | C16H14F2N6OS | 4.16 | 1 | 10 | |
| CHEMBL2403108 | C28H36ClN5O3S | 5.90 | 1 | 10 | |
| CHEMBL1081867 | C22H23F3N6S | 6.33 | 1 | 10 | |

**Table S6** (Cont.d)

| | | | | | |
|---|---|---|---|---|---|
| CHEMBL401576 | C25H26F3N5OS | 6.90 | 1 | 10 | |
| CHEMBL390649 | C28H31F3N6S | 7.50 | 1 | 10 | |
| CHEMBL3985847 | C21H26N6O4S | 2.40 | 1 | 11 | **5.46** |
| CHEMBL1782574 | C23H35F3N4O3S | 2.41 | 1 | 11 | |
| CHEMBL1091605 | C29H36F3N3O4S | 4.52 | 1 | 11 | |
| CHEMBL2204260 | C22H25F3N4O3S | 4.62 | 1 | 11 | |
| CHEMBL3407784 | C20H19ClN6O3S | 4.89 | 1 | 11 | |
| CHEMBL4075908 | C20H21BrFN5O3S | 5.46 | 1 | 11 | |
| CHEMBL192 | C22H30N6O4S | 5.48 | 1 | 11 | |
| CHEMBL2041175 | C31H27N7O3S | 5.82 | 1 | 11 | |
| CHEMBL1082111 | C22H23F4N5OS | 5.90 | 1 | 11 | |
| CHEMBL402016 | C28H27F3N6OS | 6.00 | 1 | 11 | |
| CHEMBL411293 | C28H28F3N7S | 6.60 | 1 | 11 | |

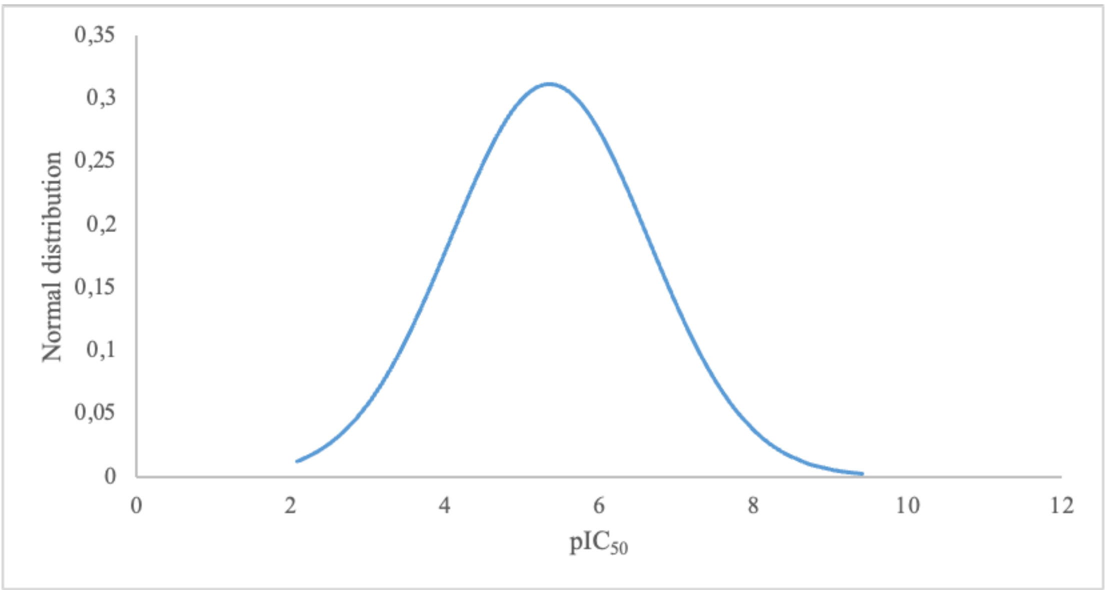

**Figure S1.** Normal distribution analysis of hERG blockers in a subset 508 compounds.

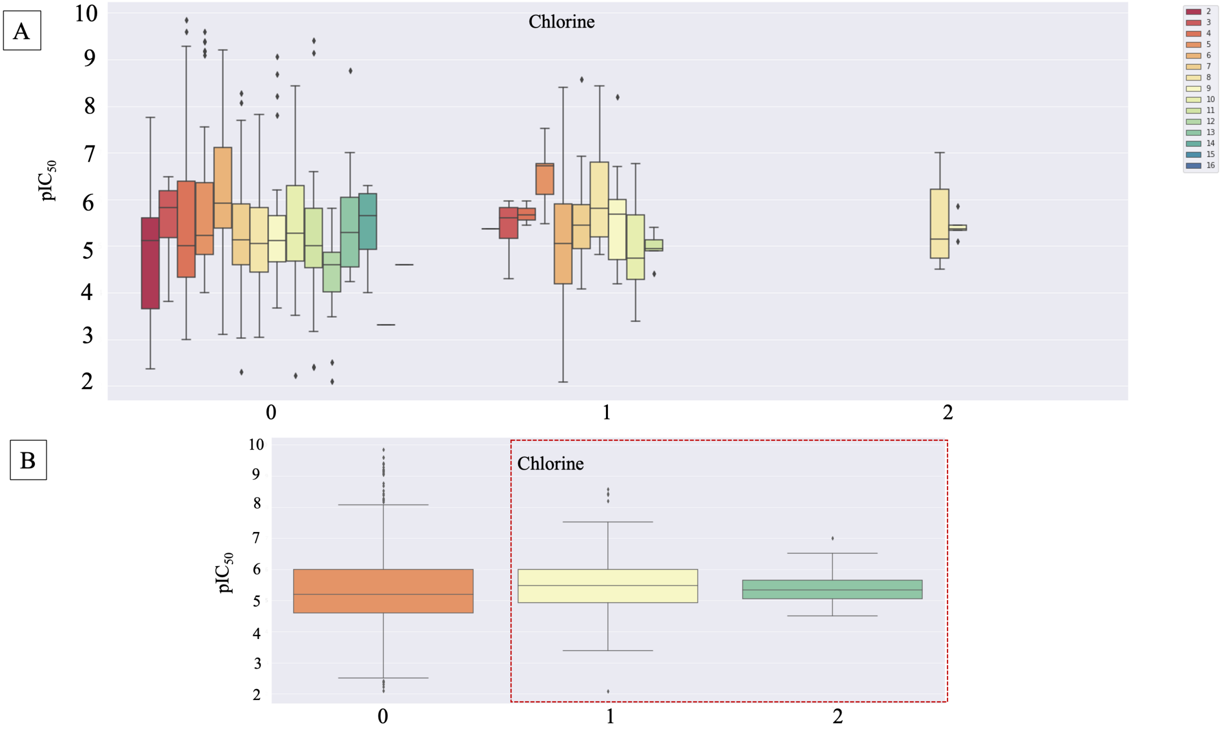

**Figure S2.** Influence of heteroatoms on the activity of the hERG blockers. (A) the effect of the number of chlorine atoms to the hERG blockage activity ($pIC_{50}$). (B) Detailed analyzes on the influence of the chlorine atoms on the $pIC_{50}$ values.

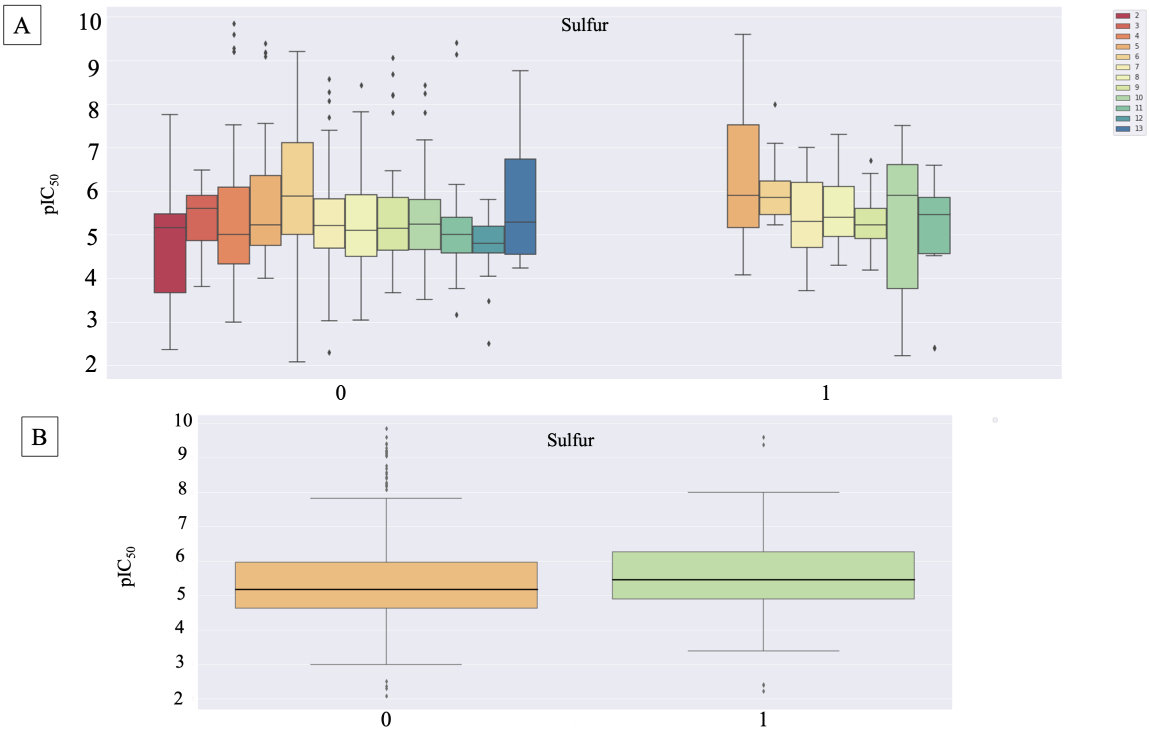

**Figure S3.** Influence of heteroatoms on the activity of the hERG blockers. (A) the effect of the number of sulfur atoms to the hERG blockage activity ($pIC_{50}$). (B) Detailed analyzes on the influence of the sulfur atoms on the $pIC_{50}$ values.

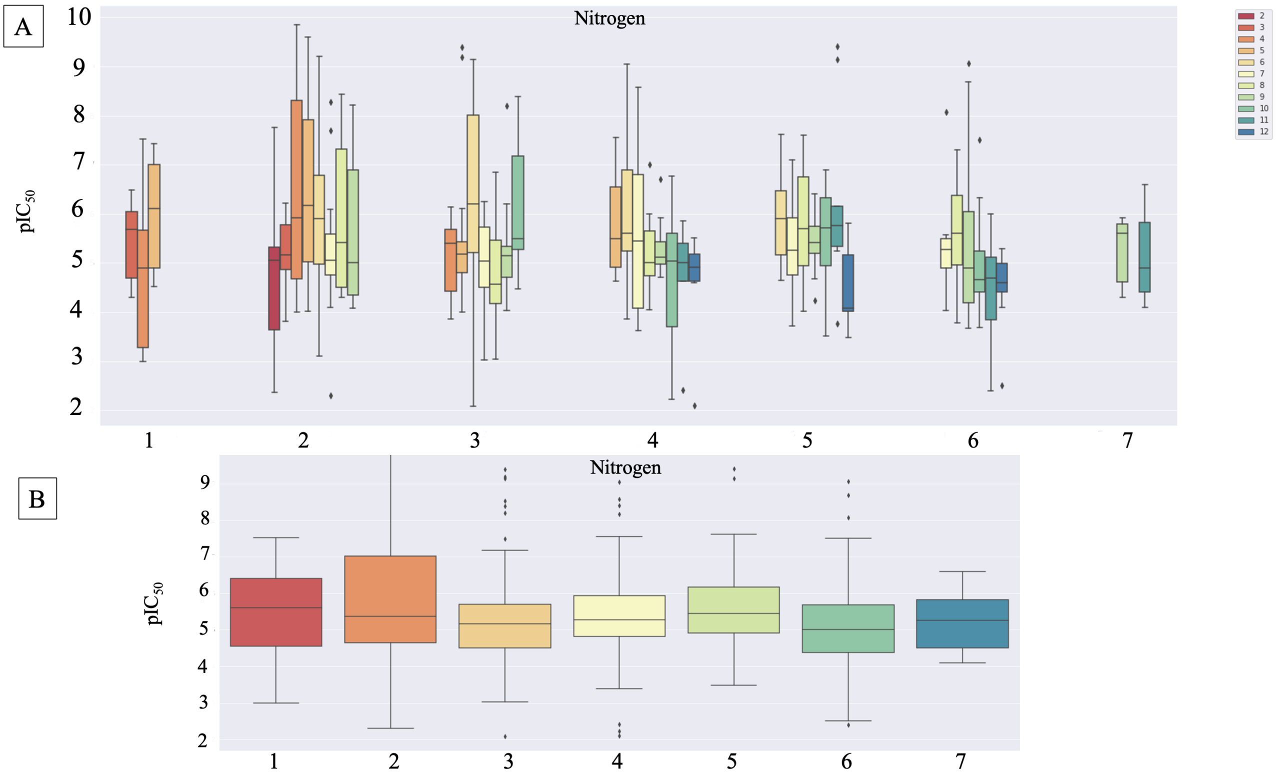

**Figure S4.** Influence of heteroatoms on the activity of the hERG blockers. (A) the effect of the number of nitrogen atoms to the hERG blockage activity ($pIC_{50}$). (B) Detailed analyzes on the influence of the nitrogen atoms on the $pIC_{50}$ values.

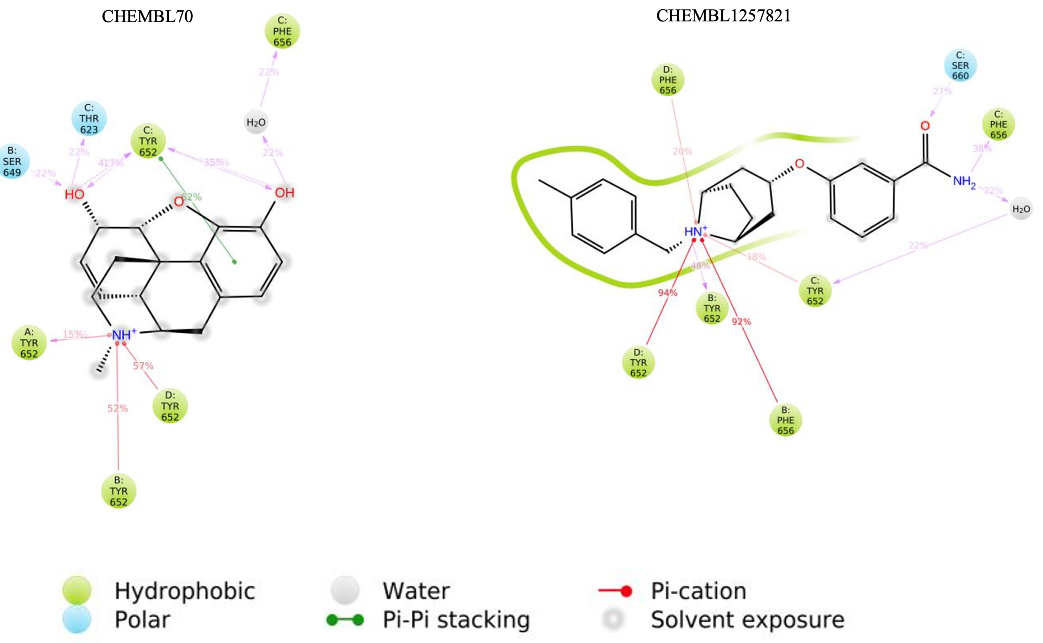

**Figure S5.** 2D protein-ligand interaction for pair 1 compounds.

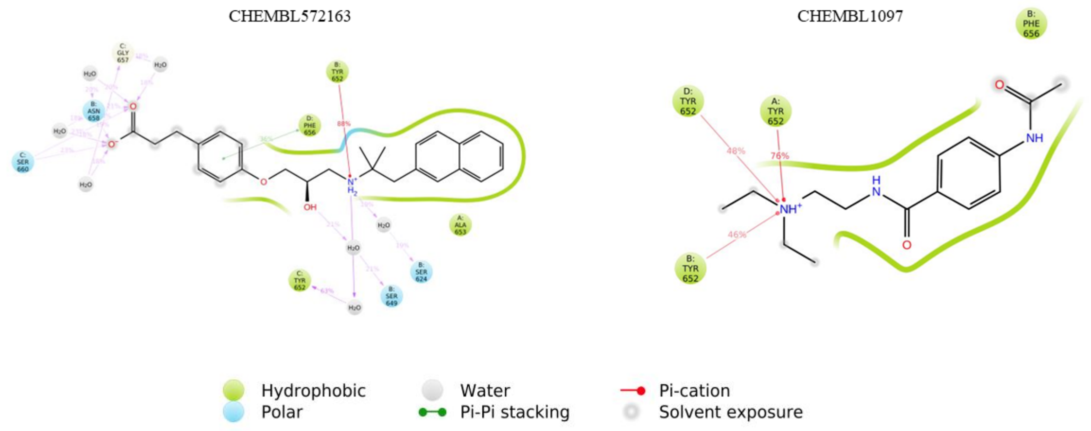

**Figure S6.** 2D protein-ligand interaction for pair 2 compounds.

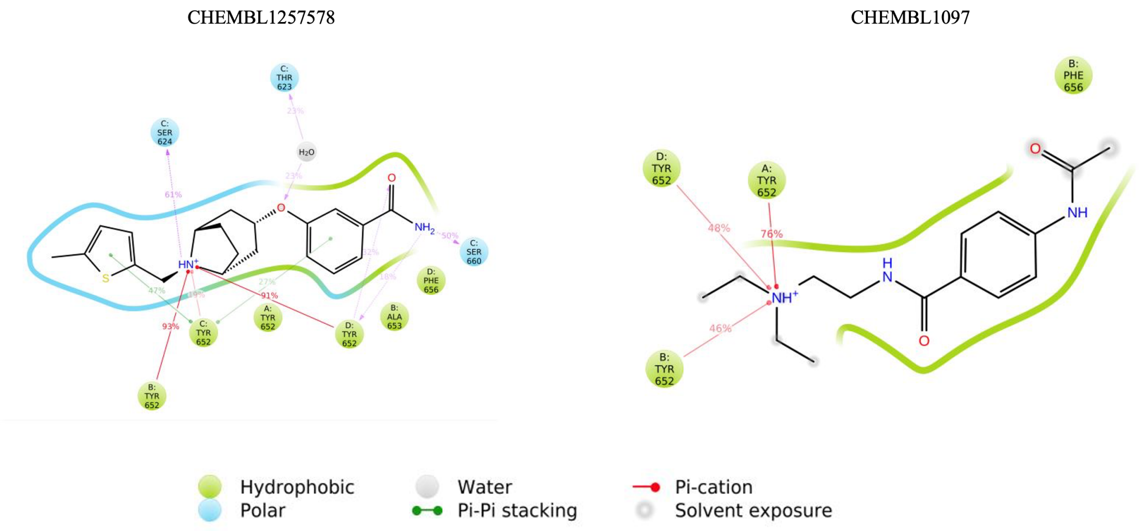

**Figure S7**. 2D protein-ligand interaction for pair 3 compounds.

CHEMBL1782574 CHEMBL3422978

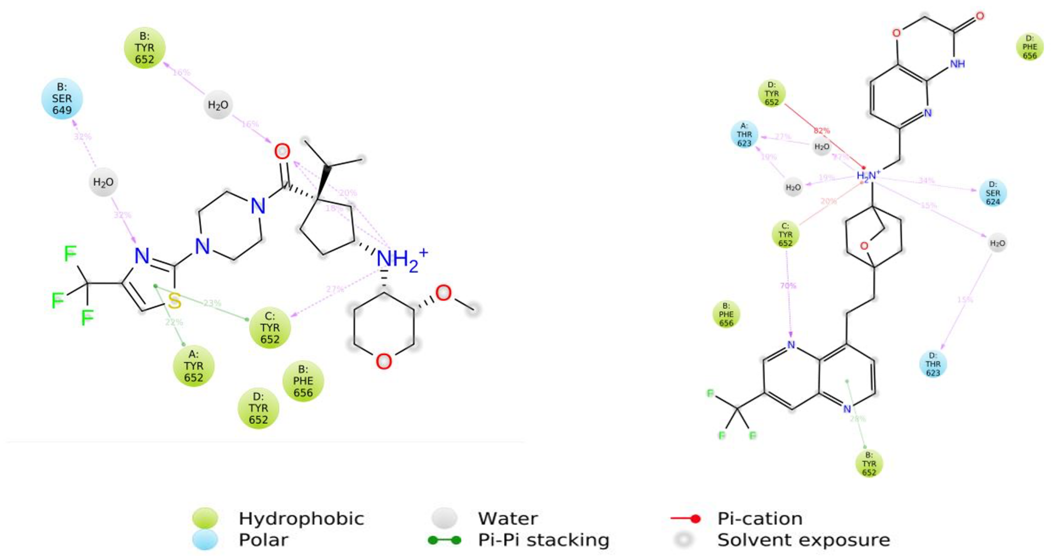

**Figure S8.** 2D protein-ligand interaction for pair 4 compounds.

CHEMBL2424928 CHEMBL3900649

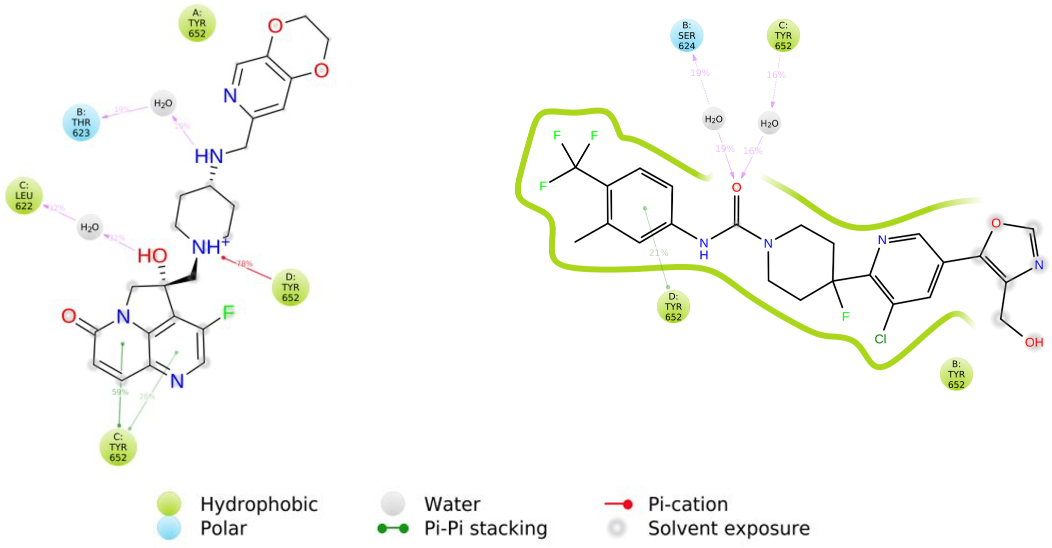

**Figure S9.** 2D protein-ligand interaction for pair 5 compounds.

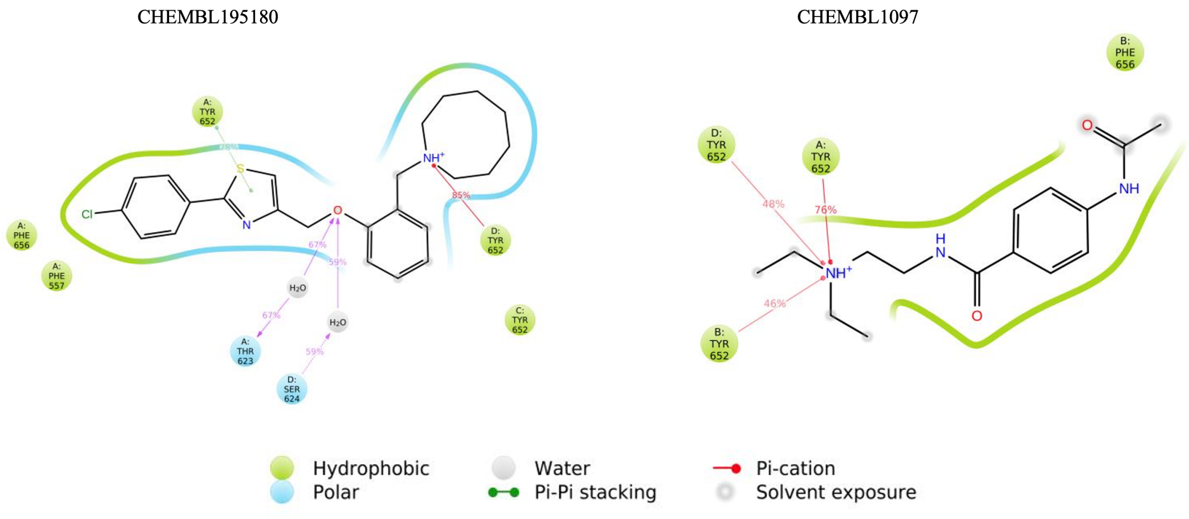

**Figure S10.** 2D protein-ligand interaction for pair 6 compounds.

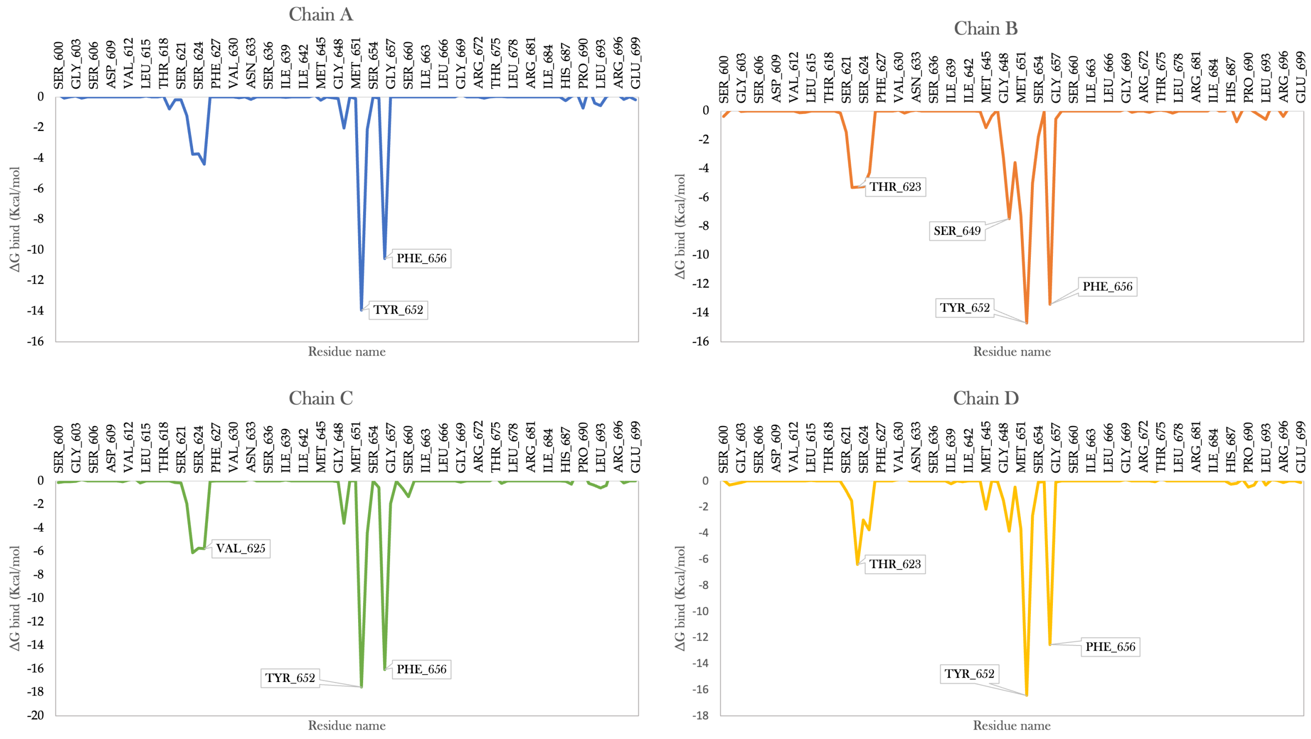

**Figure S11.** Per-residue MM/GBSA analysis for CHEMBL70 compound.

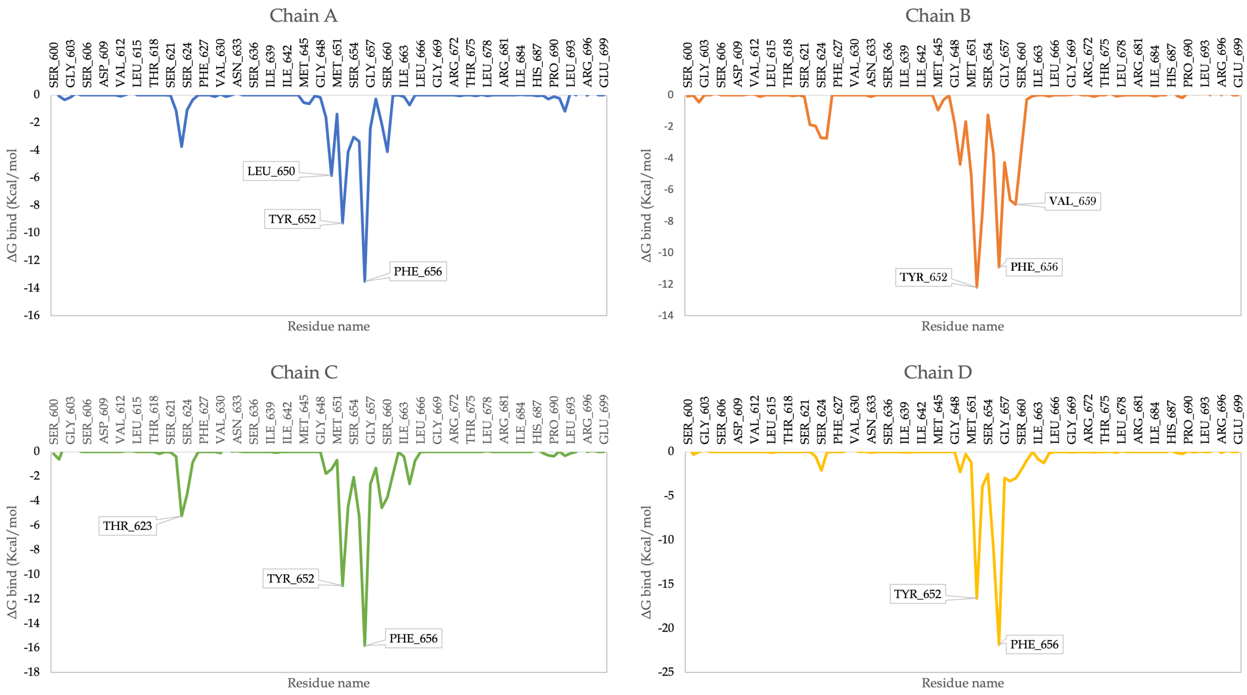

**Figure S12.** Per-residue MM/GBSA analysis for CHEMBL572163.

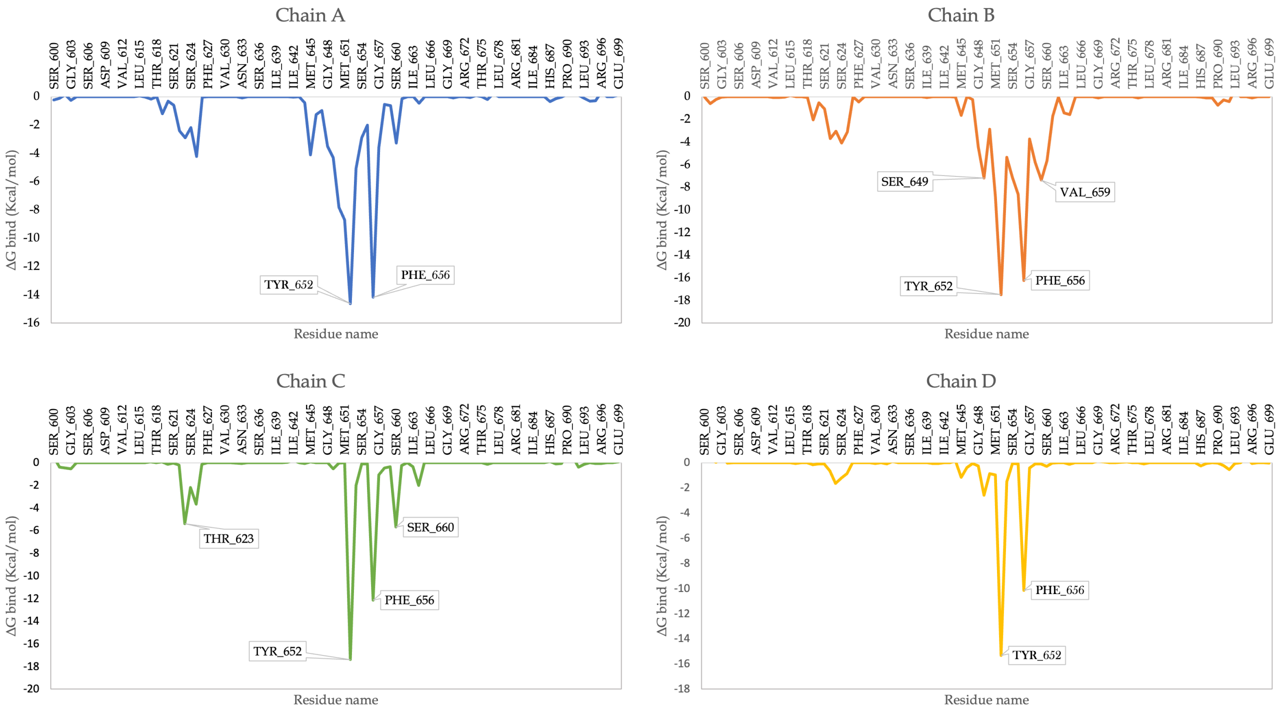

**Figure S13.** Per-residue MM/GBSA analysis for CHEMBL1097.

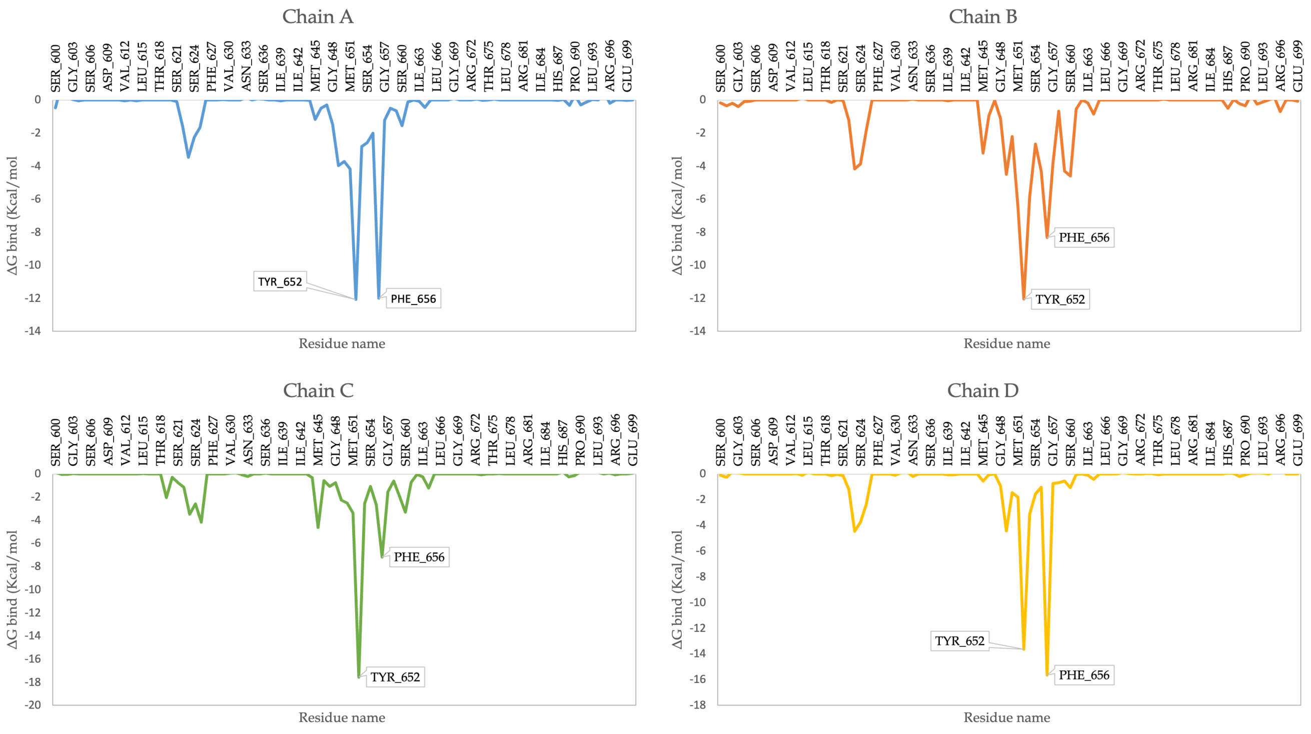

**Figure S14.** Per-residue MM/GBSA analysis for CHEMBL1782574.

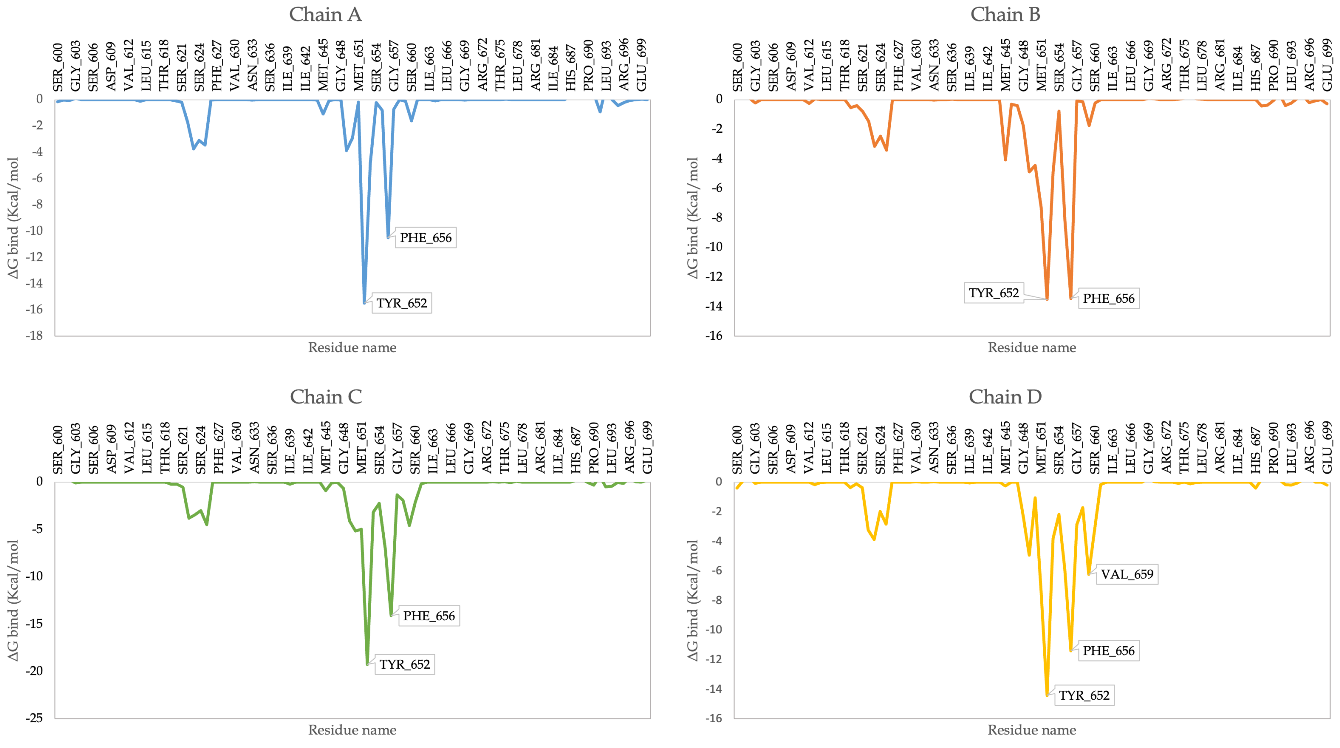

**Figure S15.** Per-residue MM/GBSA analysis for CHEMBL3422978.

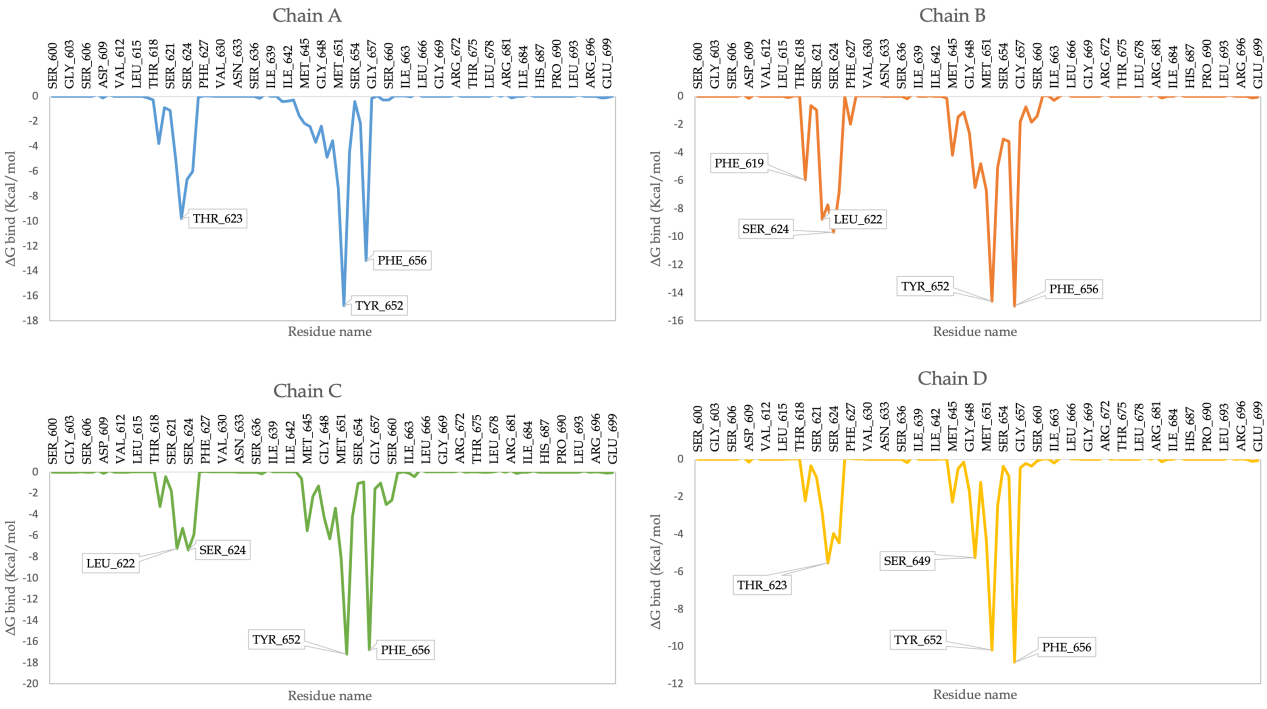

**Figure S16.** Per-residue MM/GBSA analysis for CHEMBL2424928.

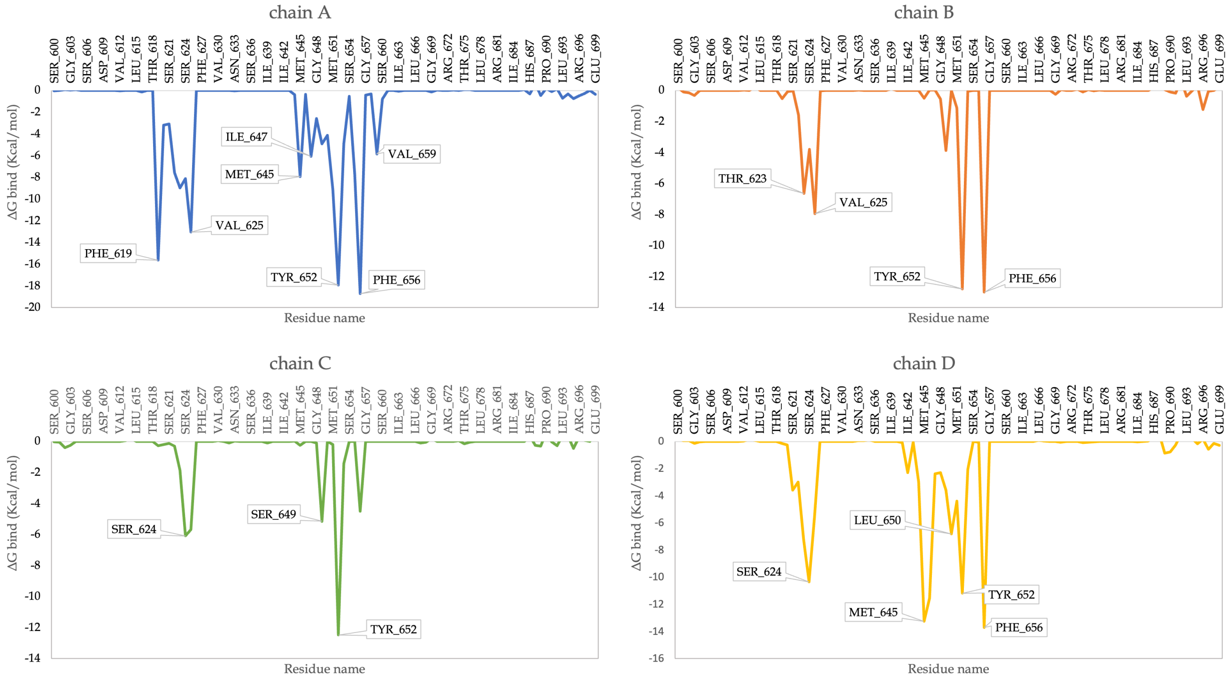

**Figure S17.** Per-residue MM/GBSA analysis for CHEMBL195180.

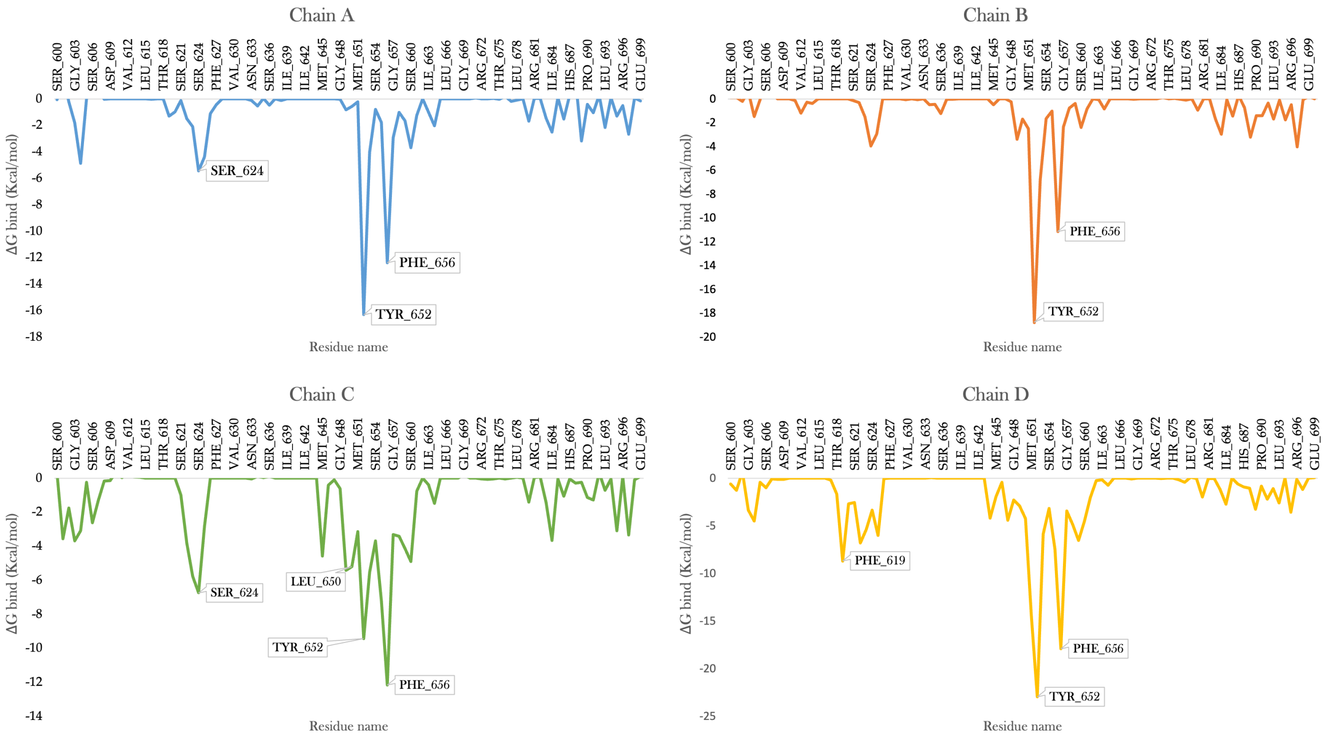

**Figure S18.** Per-residue MM/GBSA analysis for CHEMBL390649.

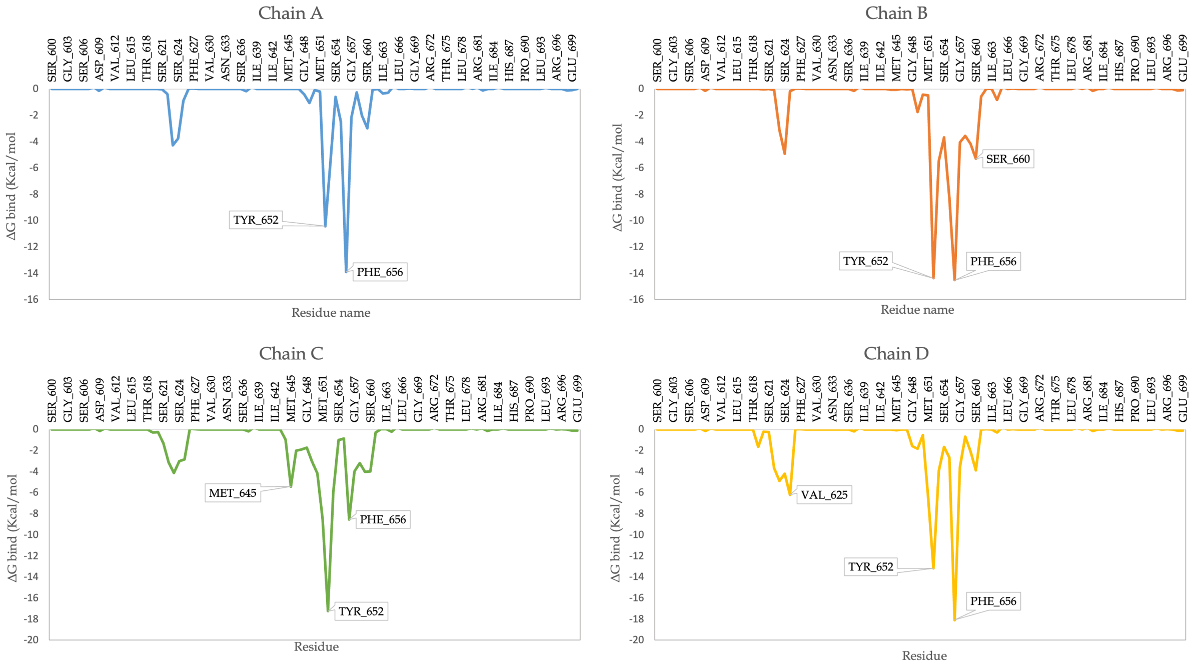

**Figure S19.** Per-residue MM/GBSA analysis for CHEMBL1257821.
